# A bone-derived protein primes rapid visual escape via GPR37 receptor in a subpopulation of VTA GABAergic neurons

**DOI:** 10.1101/2024.08.12.607510

**Authors:** Xuemei Liu, Juan Lai, Xiang Gao, Lina Wang, Yali Hu, Guoyang Liao, Shaohua Ma, Bo Feng, Liang Yang, Zhengjiang Qian, Liming Tan, Xiang Li, Liping Wang

## Abstract

Rapid escape against visual threats is critical for survival. Whether it requires a permissive mechanism is unknown. Here, we show that osteocalcin (OCN), a protein produced by bone and persisted in the brain, primes rapid visual escape response by increasing excitability of VTA GABAergic neuron subpopulation via OCN-GPR37-cAMP-THIK-1 (K_2P_13.1) pathway. Knock-out of OCN or its receptor GPR37, and conditional knock-out of GPR37 in VTA GABAergic or glutamatergic neurons caused delayed escape. Reconstituting OCN-GPR37 signaling specifically in VTA was sufficient to restore normal response. Single-cell transcriptomics combined with electrophysiology showed that OCN decreases potassium currents in a subpopulation of VTA GABAergic neurons via GPR37-induced cAMP reduction and subsequent THIK-1 suppression. This elevation of excitability in VTA neuron subpopulation can be recapitulated by HM4Di, an inhibitory chemogenetic GPCR commonly used to suppress neuronal activity. Our study demonstrated that visual behavior requires a bone-derived protein that tunes electrophysiology of central nervous system neurons.

**Graph Abstract:** *In brief:* The bone-brain axis regulates visual escape behavior. Osteocalcin (OCN), a small protein secreted by bone, plays a permissive role in rapid visual escape. This is achieved through GPR37 receptor expressed in a subpopulation of VTA GABAergic neurons. OCN-GPR37 signaling increases neuronal excitability by decreasing K+ current via THIK-1 channel suppression.

*Highlights:*

- The bone-derived protein osteocalcin and its receptor GPR37 in VTA are required for rapid visual escape response.
- GPR37 is expressed in a subpopulation of VTA GABAergic neurons and primes rapid visual escape.
- Osteocalcin-GPR37 signaling increases neuronal excitability by suppressing THIK-1 (K_2P_13.1) channel via cAMP reduction.
- Activation of HM4Di increases excitability of subpopulation of VTA GABAergic neurons and enhances rapid visual escape. 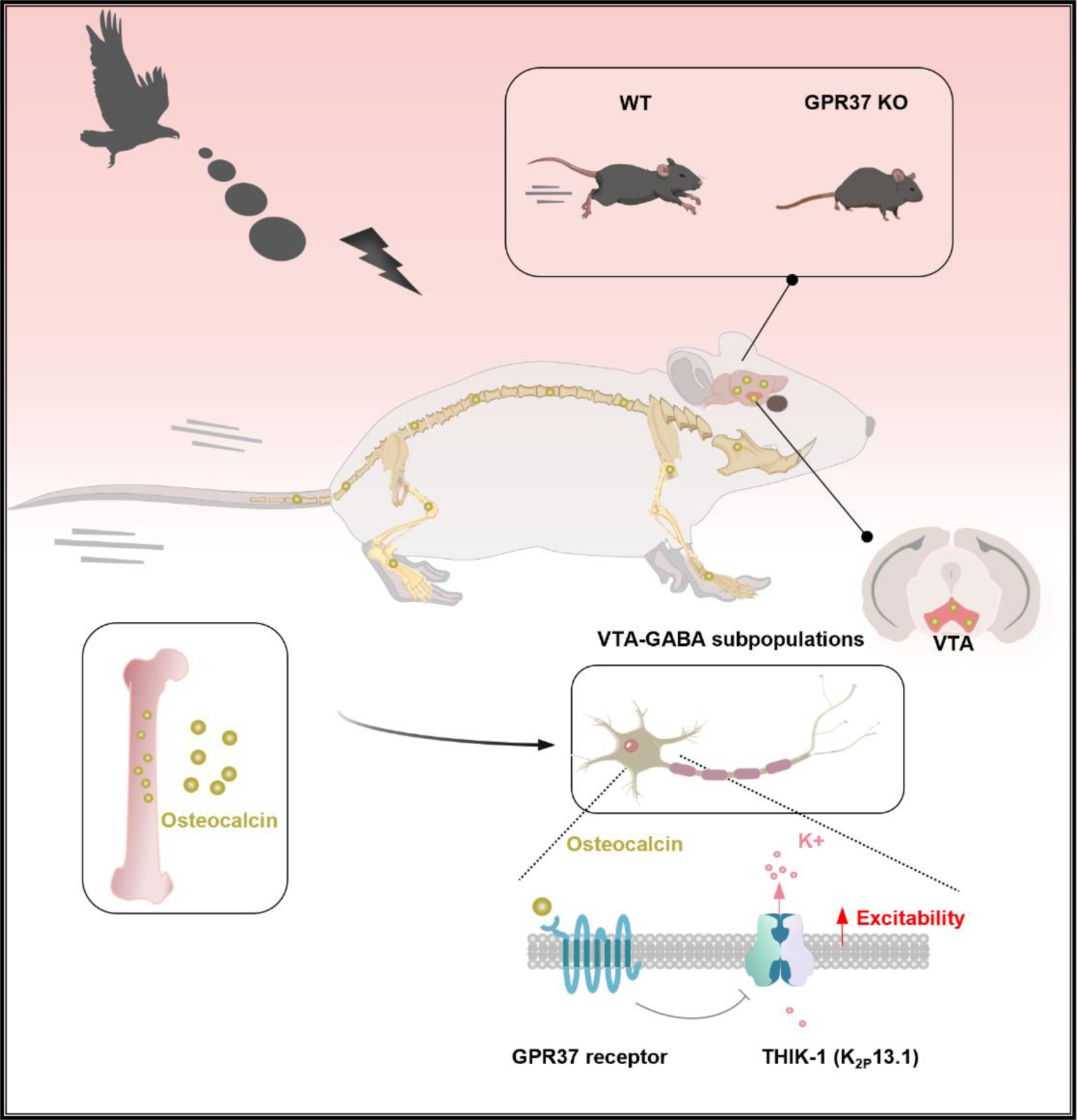

## INTRODUCTION

For vertebrates to survive in complex and life-threatening environment, the brain must coordinate with the body to rapidly transform salient sensory cue into appropriate defensive responses ^1–6^. The looming stimulus mimics a fast-approaching predator from above and is a well-established paradigm for studying diverse visually evoked defensive behaviors, including freezing, escape and attack. The neurobiological underpinnings of these defensive behaviors have been extensively studied, revealing distinct circuits comprising brain regions responsible for visual information detection, visual-motor transformation, and motor outputs ^7–15^. Of these brain regions, VTA, a midbrain nuclei traditionally viewed as a reward and motivation processing center ^16–20^, has been identified to be an information relay center for visual escape response to looming stimulus ^12,21^. VTA GABAergic neurons receive visual information from superior colliculus (SC), and send outputs to central nucleus of the amygdala (CeA) that elicit the motor response ^12^. However, whether this circuit is self-sufficient for rapid visual escape response to looming stimulus has been overlooked.

Chemicals and proteins that are secreted from different organs in the body, pass blood-brain barrier and get into the brain, have been shown to coordinate the brain and body for numerous defensive related behaviors. For instance, glucocorticoids and adrenaline are known to induce physiological changes like increased heart rate, blood pressure, and respiration rate, which are crucial for maintaining emotional and defensive responses ^22–25^. Furthermore, osteocalcin (OCN), a protein solely produced in bone and persisted in the brain and body ^26–29^, has recently been identified to regulate stress-induced defensive behaviors including reactions to electrical shock, tail suspension, and TMT exposure in adrenal insufficiency condition ^28,30^. Thus, bone, once viewed only as structural components of the body, have emerged as an endocrine organ that influence brain function and behavior ^26–33^. Nonetheless, whether and how OCN contributes to rapid visual escape response is unknown.

In this study, we demonstrated that the bone-derived protein OCN enables rapid visual escape via GPR37 receptor in a subpopulation of VTA GABAergic neurons. OCN-GPR37 pathway increases neuronal excitability by suppressing THIK-1 (tandem pore domain halothane inhibited K^+^ channel) channel activity via non-canonical GPCR signaling. Furthermore, we showed that chemogenetic activation of the Gi/o-coupled hM4Di DREADD also exerts excitatory effects on VTA GABAergic neuron subpopulation and promote rapid visual escape, in contrary to its well-accepted inhibitory effect on neuronal activity and behavior. Our findings established a new way of body-brain interaction by which the bone-derived protein OCN tunes electrophysiology of selective VTA neurons via GPR37 receptor to prime rapid visual escape, a key behavior for animal survival.

## RESULTS

### Osteocalcin is required for rapid visual escape response

Osteocalcin (OCN) is identified as a protein hormone produced by bone. It enters the circulation system upon secretion and can pass blood-brain-barrier. Previous studies have demonstrated OCN is necessary for an acute stress response ^30^. To ask whether OCN plays a role in responding to visual threat, we first measured the circulating level of undercarboxylated, Gla-OCN, and looming-evoked innate defensive responses (Figure 1A-B) in 3-month-old (young) and 7-month-old (old) wild-type (WT) mice. We found serum OCN declined 52% in old mice compared to young mice (Figure 1C). When presented an overhead looming stimulus, the old mice exhibited higher response latency to the stimuli and longer return time to nest compared to young mice (Figure 1D). Next, we performed an intraperitoneal (i.p.) injection of osteocalcin in old mice. Administration of osteocalcin, but not saline, enabled rapid visual escape in old mice (Figure 1B,1E). These results indicated that circulating OCN levels are associated with rapidness of visual escape response. This conclusion was further validated using young *OCN* mutant mice. Compared with WT and *OCN*+/- mice, *OCN*-/- mice had significantly higher response latency, longer return time and less time spent hiding in the nest (Figure 1F). Taken together, the bone-derived OCN is essential for efficient innate defensive responses to looming stimulus.

**Figure 1.**
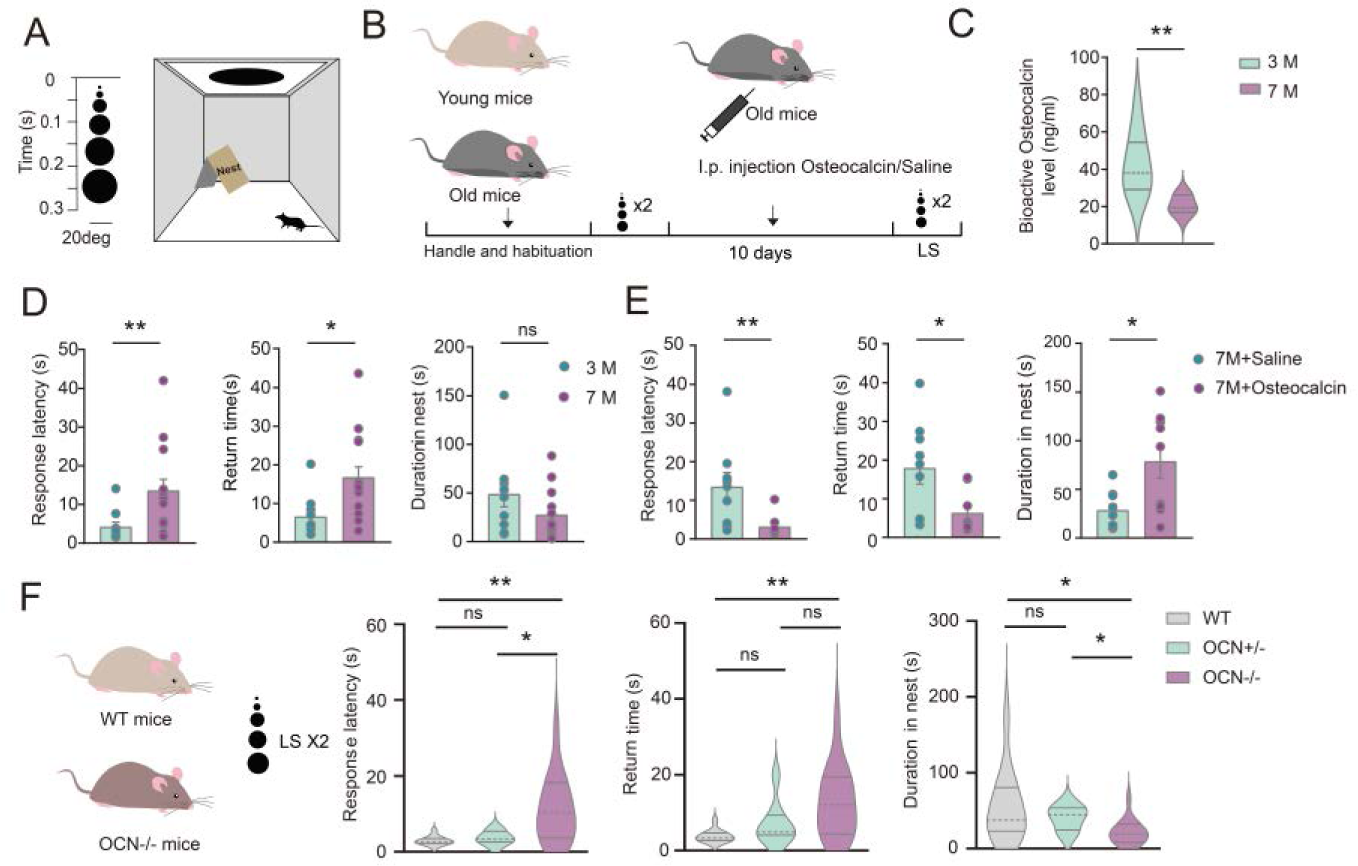
Osteocalcin Mediates Innate Defensive Responses to Looming Stimulus. **(A)** Schematic of behavior paradigm. **(B)** Experimental design for results in panels D and E. **(C)** Serum osteocalcin level in young (n=8) and old (n=9) WT mice (Two-tailed unpaired t test, **p=0.0014). **(D)** Bar graphs showing metrics of looming responses of young (3-month-old WT mice, n=10) and old (7- month-old WT mice, n=15) mice; . Left: response latency (Mann Whitney U test, **p=0.0054); middle: return time (Two-tailed unpaired t test, *p=0.0105); right: duration in the nest (Two-tailed unpaired t test, p=0.125). **(E)** As in D, but for old mice (7-month-old) after i.p. injection (saline group, n=9; Osteocalcin group, n=9). Left: response latency (Mann Whitney U test, **p=0.0040); middle: return time (Two-tailed unpaired t test, , *p=0.0197); right: duration in the nest (Two-tailed unpaired t test, *p=0.0148). **(F)** Bar graphs showing metrics of looming responses of WT, *OCN*+/- and *OCN*-/- mice (WT, n=12; *OCN*+/-, n=8; *OCN*-/- mice, n=15). Left: response latency (One-way ANOVA, **p=0.0014); middle: return time (One-way ANOVA, **p=0.0046); right: duration in the nest (One-way ANOVA, *p=0.0211).

### Rapid visual escape requires GPR37 receptor in the VTA

Previous studies have identified GPR158 and GPR37 as receptors for OCN in the brain ^28,34^. To investigate the potential roles of GPR37 and GPR158 in looming-evoked defensive response, we tested *GPR37* and *GPR158* mutant mice. We found that *GPR37*-/- mice exhibited significantly higher response latency and longer return time compared to *GPR37* heterozygous, while no significant differences were observed between *GPR158*-/- and heterozygous mice Figure 2A-2B, Figure S1). Thus, GPR37 receptor, but not GPR158 receptor, plays an essential role in the defensive response to looming stimuli.

**Figure 2.**
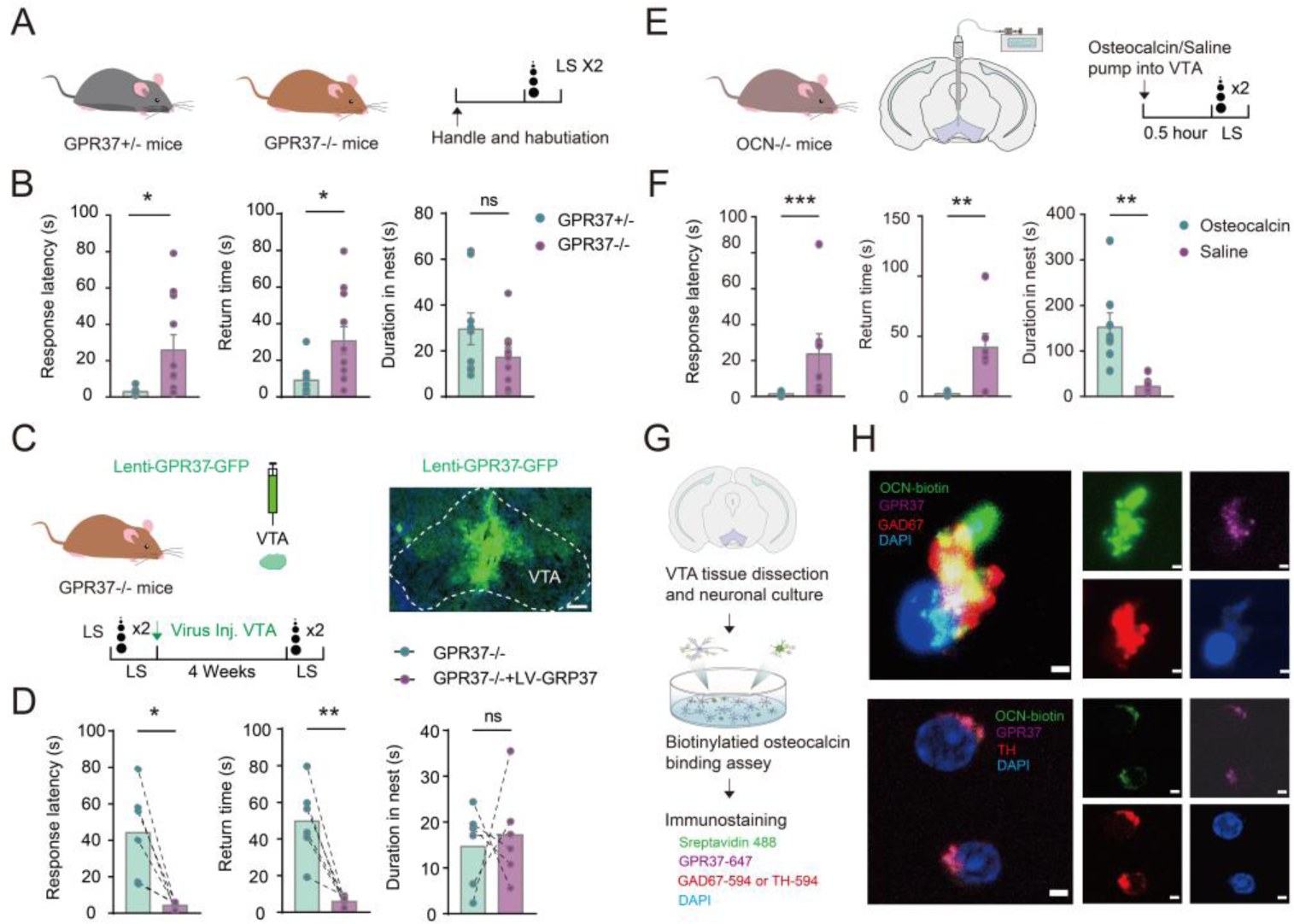
GPR37 in VTA regulates Looming-Evoked Flight Behaviors. **(A)** Schematic of looming test for *GPR37*+/- and *GPR37*-/- mice. **(B)** Bar graphs showing metrics of looming responses of GPR37+/- (n=9) and GPR37-/- (n=11) mice. Left: response latency (Two-tailed unpaired t test, *p=0.0239); middle: return time (Two-tailed unpaired t test, *p=0.0235); right: duration in the nest (Two-tailed unpaired t test, p=0.1331). **(C)** Experimental design for testing the effect of GPR37 restoration in VTA in *GPR37* mutant mice. **(D)** As in B, but for *GPR37*-/- mice with (n=6) or without (n=6) virus injection. Left: response latency (Two-tailed paired t test, * p=0.0109); middle: return time (Two-tailed paired t test, ** p =0.0044); right: duration in the nest (Two-tailed paired t test, p=0.7533). **(E)** Experimental design for testing the effect of OCN restoration in VTA in *OCN*-/- mice. **(F)** As in B, but for *OCN*-/- mice injected with OCN (n=7) or saline (n=8). Left: response latency (Mann Whitney U test, ***p=0.0003); middle: return time (Two-tailed unpaired t test, **p=0.0025); right: duration in the nest (Two-tailed unpaired t test, **p=0.0020). **(G)** Schematic showing VTA neuronal culture experiment. **(H)** Representative images showing immunostaining of osteocalcin-biotin and GPR37 receptors in GABAergic neurons and DA neurons. Green: osteocalcin-biotin and steptavidin-488; Blue: DAPI; Purple: GPR37, Red: TH or GAD67. Scale bar= 2 μm. Error bars represent SEM. p<0.05 is significant.

Previous studies indicated OCN mainly functions in CA3 region of the hippocampus, dorsal raphe, and VTA^29^. Additionally, our group previously identified VTA as a circuit hub for looming-evoked defensive responses, which receives input from visual processing center SC and sends output to CeA to elicit escape behavior ^12^. To test whether GPR37 receptor in VTA is involved in looming-induced escape response, we reintroduced GPR37 expression into the VTA of *GPR37*-/- mice via Lenti-GPR37-GFP injections. Specific restoration of GPR37 in VTA led to a remarkably faster flee to the nest (Figure 2C-2D). These results indicated that GPR37 receptor in the VTA is required for rapid looming-induced escape. Next, we administered OCN into VTA in *OCN*-/- mice to restore OCN level. The restoration of OCN specifically in VTA, led to much shorter response latency and return time, as well as longer duration spent in the nest (Figure 2E-2F, Figure S2). These results indicated that VTA is the target area of OCN in regulating looming responses.

Furthermore, we showed that OCN binds to GPR37 receptor in VTA neurons by treating primary neuronal culture from VTA with biotinylated OCN. Immunostaining results showed that Streptavidin 488, which binds to biotinylated OCN, colocalized with GPR37 receptors in both tyrosine hydroxylase (TH) positive neurons and GAD67 positive neurons (Figure 2G- 2H). These data suggested that OCN binds to the GPR37 receptor in both VTA GABAergic neurons and VTA dopaminergic neurons.

### GPR37 in VTA GABAergic and glutamatergic neurons regulates rapid visual escape

To identify through which VTA neuron type OCN-GPR37 signaling regulates rapid visual escape, we selectively removed GPR37 receptor in VTA dopaminergic (DA), glutamatergic and GABAergic neurons, respectively. And tested looming-induced escape responses in conditional knock-out mice. GPR37 ablation in all DA neurons in the brain did not cause any significant changes in looming responses (Figure S3). Consistently, selective ablation of GPR37 in VTA DA neurons also did not result in significant differences in looming responses. (Figure 3C-3D).

**Figure 3.**
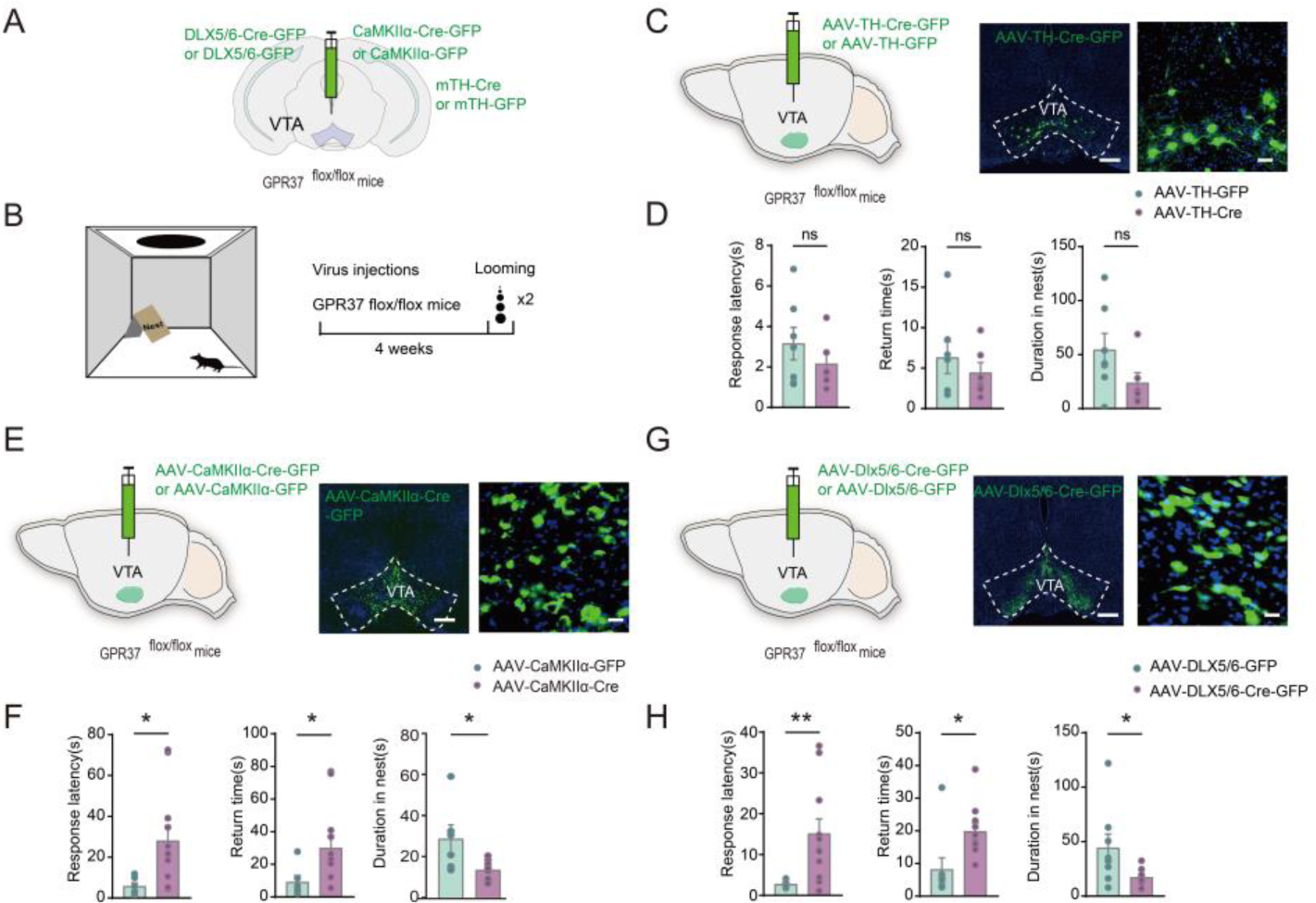
GPR37 receptor in VTA GABA or glut neurons but not DA neurons mediates rapid visual escape. **(A)** AAV expressing Cre with specific neuronal promoter injected into *GPR37*^LoxP/LoxP^ mice. **(B)** Looming test design. **(C)** Conditional deletion of GPR37 in VTA TH+ neurons. **(D)** Bar graph showing looming response in mice with conditional deletion of GPR37 in VTA dopaminergic neurons (n=7) and control mice (n=6). Left: response latency (Two-tailed unpaired t test, p=0.3353); middle: return time (Two-tailed unpaired t test, p=0.4622); right: duration in the nest (Two- tailed unpaired t test, p= 0.1308). **(E)** Conditional deletion of GPR37 in VTA CaMKⅡα+ neurons. **(F)** As in D, but for mice with conditional deletion of GPR37 in VTA CaMKⅡα+ neurons (n=13) and control mice (n=6). Left: response latency (Two-tailed unpaired t test, *p=0.0468); middle: return time (Two-tailed unpaired t test, *p=0.0477); right: duration in the nest (Two-tailed unpaired t test, *p= 0.0114). **(G)** Conditional deletion of GPR37 in VTA GABAergic neurons. **(H)** As in D, but for mice with conditional deletion of GPR37 in VTA GABAergic neurons (n=10) and control mice (n=8). Left: response latency (Mann-Whitney U test, **p=0.0044); middle: return time (Two-tailed unpaired t test, *p=0.0162); right: duration in the nest (Two-tailed unpaired t test, *p=0.0331).

To selectively ablate GPR37 receptor in VTA glutamatergic or GABAergic neurons, we first tested the specificity and efficiency of our labeling method^35^. We injected AAV-CaMKⅡα-GFP combined with AAV-DIO-mcherry into the VTA of vesicular glutamate transporter 2 (VGluT2)-ires-Cre mice. 90% of mcherry+ cells were labeled with GFP (Figure S4A-C). We injected AAV- Dlx5/6-GFP combined with AAV-DIO-mchery into the VTA of glutamate decarboxylase 2 (GAD2)-ires-Cre mice. 98% of mcherry+ cells were labeled by GFP, and 84% of GFP+ cells were labeled by mcherry. Very few cells (<1%) that were co-labeled by mcherry and GFP were positive for DA marker, TH (Figure S4D-F).

Next, we used AAV-CaMK Ⅱα -Cre-GFP or AAV-Dlx5/6-Cre-GFP virus injected into VTA of *GPR37*^flox/flox^ mice to explore the functional participation of GPR37 receptor in VTA glutamatergic or GABAergic neurons in looming responses. Specific ablation of GPR37 in VTA glutamatergic neurons (Figure 3E-3F) or GABAergic neurons (Figure 3G-3H) resulted in significant impairment of looming-induced escape responses.

Taken together, GPR37 receptor in VTA glutamatergic and GABAergic neurons, but not VTA-DA neurons, regulates looming-induced escape responses.

### Characterization of GPR37 Expression in Distinct Neuronal Populations in the Ventral Tegmental Area

The VTA is a well-known heterogenous nucleus implicated in reward, motivation, alerting and various neuropsychiatric disorder ^20^. To understand the cellular and molecular mechanisms underlying the function of GPR37 receptor in VTA on the visual escape response, it is crucial to comprehensively characterize the cellular composition of VTA. To achieve this, we used 10x Chromium platform to perform snRNA-seq of two biological replicates, with each containing VTA area from multiple male mice (STAR Methods). The resulting gene expression matrices were filtered to remove low-quality cells and doublets^36^. In total, we obtained 23,029 high-quality nuclear transcriptomes (Figure S5A, B).

We used dimensionality reduction (uniform manifold approximation and projection, UMAP) and unsupervised clustering to derive a cell-type atlas in VTA. Cells were first classified into neurons (n=11,063) and non-neuronal cells (n=11,685) via differential expression of *Snap25* (Figure S5A, C). Neurons were comprised of Glut (n=2,038), GABA (n=1,966), GABA-Glut (n=5,736) and dopaminergic (n=1,323) types, identified by canonical markers (Figure 4A; Figure S5A and C). Notably, neurons that express both GABAergic and glutamatergic markers (GABA-Glut neurons) make up more than half of all neurons, and more than 70% of neurons expressing either GABAergic or glutamatergic markers (Figure 4B).

**Figure 4.**
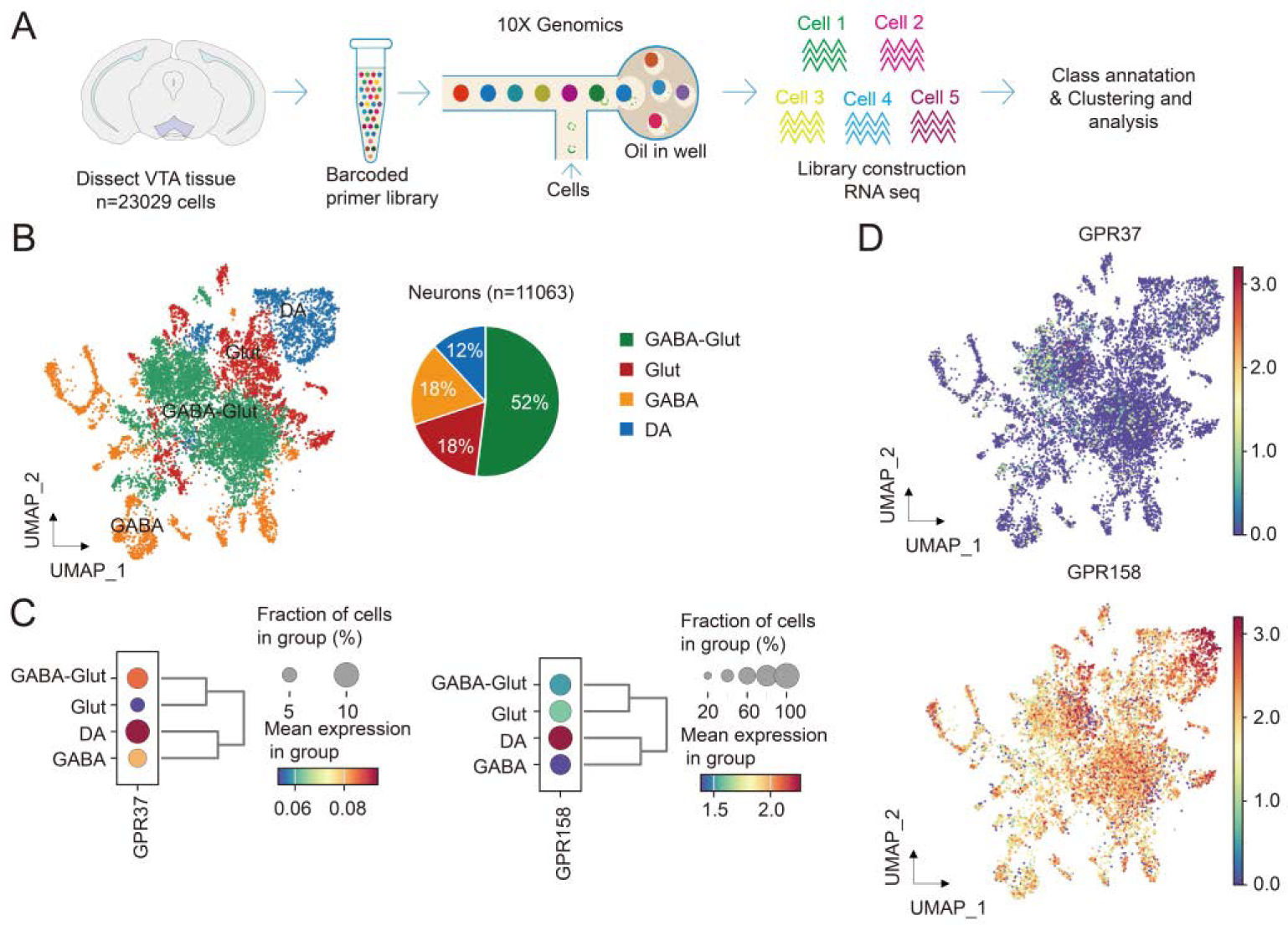
Single nucleus RNA-Seq characterized GPR37 expression in distinct neuronal populations in the Ventral Tegmental Area. **(A)** VTA snRNA-seq workflow. **(B)** UMAP (Uniform manifold approximation and projection) and plot of proportion of neuronal populations within the VTA. **(C)** Dot plots showing the expression levels of GPR37 and GPR158 in different neuronal populations within VTA. **(D)** GPR37 and GPR158 expression shown in the UMAP.

We next examined expression of GPR37 and GPR158 in the four neuronal types. These two genes showed distinct expression patterns across neuronal types. GPR158 is broadly and highly expressed in all types. In contrast, GPR37 is expressed in dopaminergic and GABAergic neurons (GABA-Glut and pure- GABA population) at lower levels, but not expressed in Glut-only neurons (Figure 4C, D; Figure S5D). Since we showed that GPR37 in VTA GABAergic and glutamatergic, but not VTA-DA neurons is required for rapid escape response, we focus our further study on VTA GABAergic neurons, within which 70% also express glutamatergic marker and higher level of GPR37.

### OCN-GPR37 signaling increases excitability in subpopulation of VTA GABAergic neurons by suppressing potassium current

To understand the neurophysiological mechanisms of OCN via the GPR37 receptor in VTA GABAergic neurons, we conducted whole-cell recordings in VTA in brain slices. We sought to assess effects of repeated OCN infusion (10ng/ml) specifically in VTA GABAergic neurons by injecting AAV-DIO-GFP in the VTA of GAD2-ires-Cre mice. We found that OCN elicits excitatory or inhibitory responses in distinct subpopulations of VTA GABAergic neurons. Among 37 cells we recorded, the spontaneous firing rate of 18 cells (Type-Ⅰ VTA GABAergic neurons) increased (Figure 5A-C). Additionally, in response to a series of current injections, the spike rate of these 18 cells increased after osteocalcin infusion (Figure S6A-D). By contrast, spontaneous firing rate of the other 19 cells (Type-Ⅱ VTA GABAergic neurons) decreased upon OCN infusion (Fig 5D). The resting membrane potential (RMP) decreased in Type-I neurons and increased in Type-II neurons in response to OCN, respectively. Other electrophysiological properties were similar between the two types of neurons, except that the holding current decreased in Type-I neurons (Figure S7A-D). Considering potassium ion channels are crucial for regulation of VTA neuronal excitability^37–39^, we performed patch clamp recording on VTA GABAergic neurons to test whether K^+^ channel-mediated current is regulated by OCN- GPR37 signaling. Consistent with results in Figure 5C and D, K+ current was decreased in Type-Ⅰ VTA GABAergic neurons (Fig 5E-F), and increased in Type-Ⅱ VTA GABAergic neurons by OCN (Fig 5G-H). Thus, OCN increases or decreases excitability in two distinct populations of VTA GABAergic neurons, and correspondingly suppresses or enhances K+ current in these neurons.

**Figure 5.**
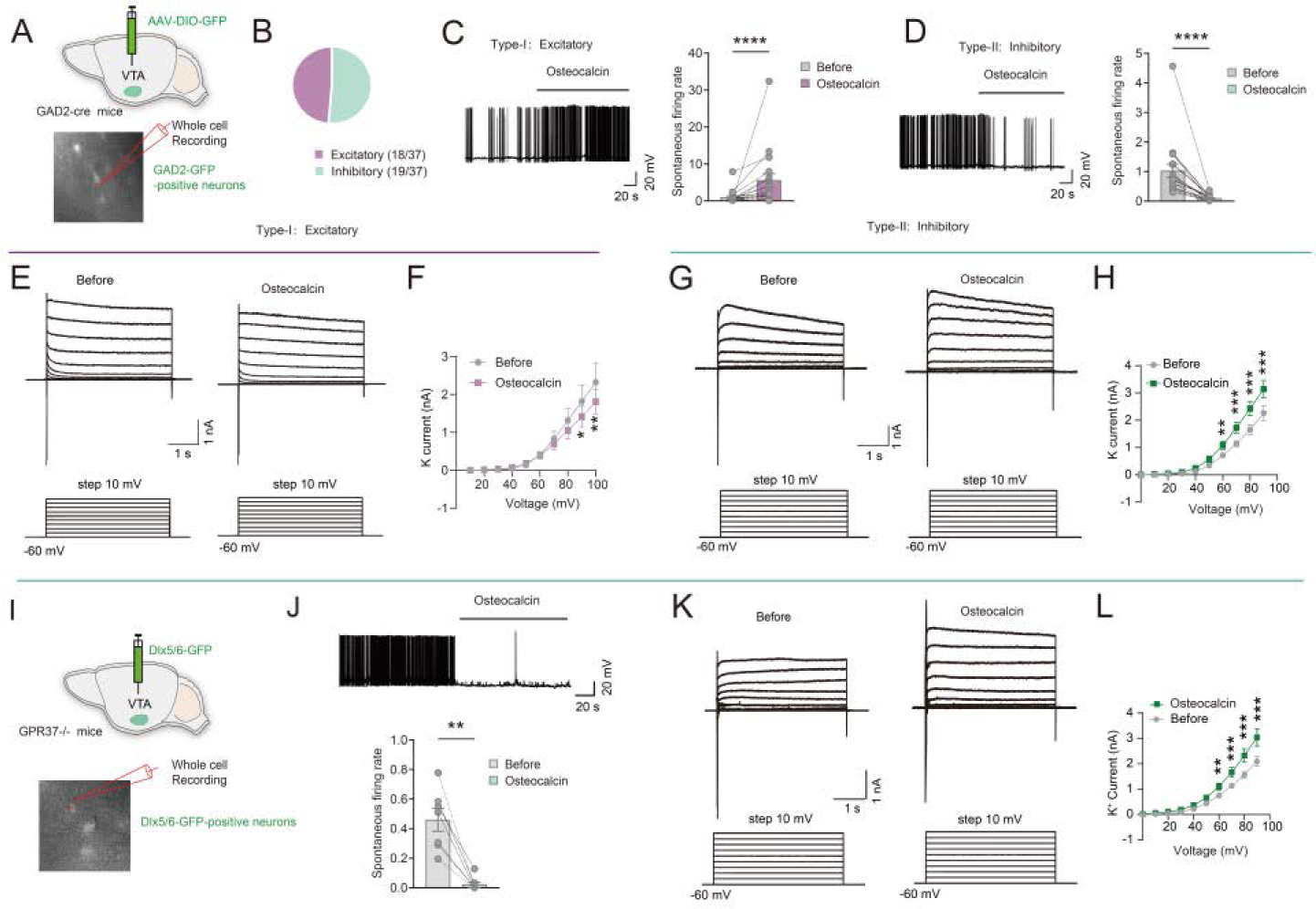
OCN elicits excitatory or inhibitory responses in distinct subpopulations of VTA GABAergic neurons with opposite K+ currents changes. **(A)** Schematic showing recording of VTA GAD2 positive neurons using patch-clamp. **(B)** Proportions of excitatory and inhibitory effects induced by OCN. **(C)** Spontaneous firing rate of Type-I neurons were activated following treatment with OCN (10ng/ml). (Wilcoxon matched-pair signed rank test, ****p<0.0001, 18 cells from 13 mice) **(D)** Spontaneous firing rate of Type-II neurons were inhibited following treatment with OCN (10ng/ml). (Wilcoxon matched-pair signed rank test, ****p<0.0001, 19 cells from 13 mice) **(E-F)** Sample traces and quantification showing K+ channel-mediated currents decreased in Type-I neurons after infusion of OCN. (Two-way ANOVA multiple comparisons, *p=0.0335, **p=0.0032, 6 cells from 5 mice). **(G-H)** Sample traces and quantification showing K+ channel-mediated currents increased in Type-II neurons after infusion of OCN. (Two-way ANOVA multiple comparisons, ***p=0.0009, **p=0.0088, 8 cells from 6 mice). **(I)** Schematic showing recording of VTA GABAergic neurons in *GPR37*-/- mice. **(J)** As in D, but for GABAergic neurons in *GPR37*-/- mice. (Wilcoxon matched-pair signed rank test, **p=0.0078, 8 cells from 6 mice). **(K-L)** As in G-H, but for GABAergic neurons in *GPR37*-/- mice. (Two-way ANOVA multiple comparisons, ***p<0.001, **p=0.0094, 9 cells from 6 mice).

GPR158 has been shown to control neuronal excitability by regulating potassium current ^40–42^. We next examined whether GPR37 plays a role in increasing excitability and decreasing potassium current in VTA GABAergic neurons upon OCN infusion. We performed whole cell recording in brain slices of *GPR37*-/- mice infected with AAV-Dlx5/6-GFP in VTA. Spontaneous firing rate of Dlx5/6- GFP positive neurons solely deceased by repeated osteocalcin infusion (Fig 5I- J). Also, the spike rate of these neurons decreased after osteocalcin infusion in response to a series of current injections (Fig S6E-G). Consistent with changes in firing rate, K+ current was solely increased by OCN in VTA GABAergic neurons in *GPR37*-/- mice (Fig 5K-L). These results argued that the increased firing rate and decreased potassium current upon OCN infusion in Type-Ⅰ VTA GABAergic neurons depend on GPR37 receptor. In parallel, the decreased firing rate and increased potassium current upon OCN infusion in Type-Ⅱ VTA GABAergic neurons probably depend on GPR158.

### OCN increases excitability of VTA GABAergic neuron subpopulation via GPR37-K_2P_13.1 axis

Since we observed concomitant changes on neuronal excitability and potassium current by OCN in VTA GABAergic neurons, we next sought to examine the causal relationship between these two changes by targeting potassium channels. To do this, we first identified potassium channels that have the highest correlation in gene expression pattern with *GPR37* and *GPR158* in VTA GABAergic neurons. We focus our bioinformatic analysis on GABA-Glut neuron type to reduce noise due to higher GPR37 expression level compared to GABA type. Unsupervised clustering further divided GABA-Glut neurons into three subtypes, each with a distinct set of marker genes (Figure 6A, B). GPR37 had the highest expression in subtype 2 (Figure 6C), while GPR158 was expressed highest in subtype 1 (Figure 6E). Some 71 genes encoding different potassium channels were expressed in VTA GABA-Glut neurons (Figure S8). The Pearson correlation of expression in the UMAP of each potassium channel with either GPRs were calculated, together with p-value and the slope and intercept of linear regression (Figure 6C-F; Figure S9 and S10; Table 1 and 2).

**Figure 6.**
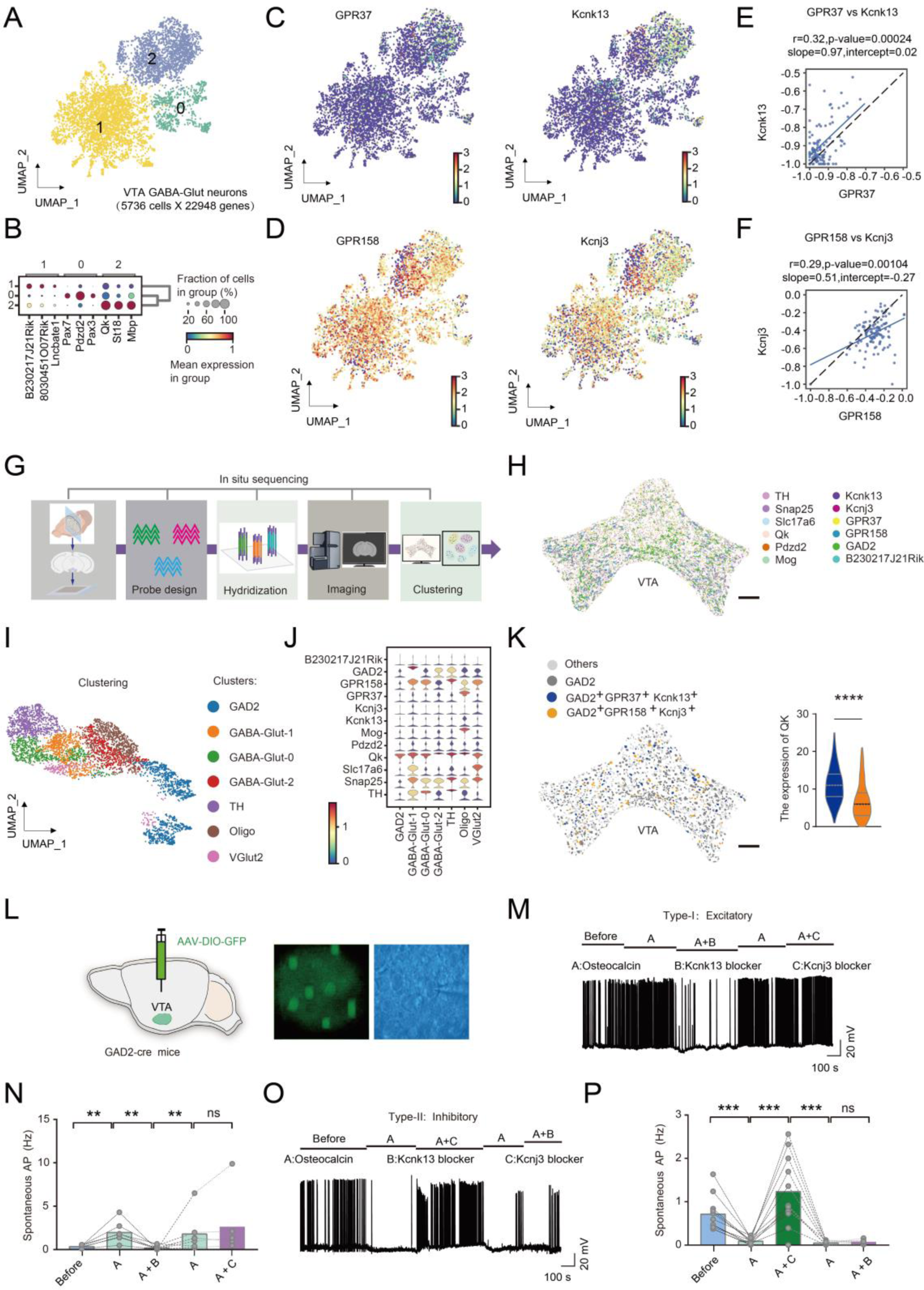
GPR37 increases excitability of a subpopulation of VTA GABAergic neurons via K_2P_13.1 encoded by *Kcnk13*. **(A)** Clustering of VTA GABA-Glut neurons identified 3 subtypes. **(B)** Marker gene expression of 3 subtypes in A. **(C)** Expression of *GPR37* and *Kcnk13* in 3 subtypes in UMAP plot. **(D)** Pearson correlation of *GPR37* and *Kcnk13* expression, with p-value, slope and intercept of linear regression. **(E)** Expression of *GPR158* and *Kcnj3* in 3 subtypes in UMAP plot.. **(F)** As in D, but for *GPR158* and *Kcnj3*. **(G)** Workflow of the *in situ* sequencing. **(H)** Spatial transcriptomics images in the VTA. Scale bar, 200 μm. **(I)** Heatmap of marker gene expression in cell clusters. **(J)** Violin plot of marker gene expression in each clusters. **(K)** Spatial transcriptomics expression of GPR37^+^ Kcnk13^+^ GAD2^+^ and GPR158^+^ Kcnj3^+^ GAD2^+^ subcluters. **(L)** Schematic showing recording of VTA GAD2 positive neurons using patch-clamp. **(M-N)** Sample traces and quantification of firing rate recorded in Type-I VTA GABAergic neurons after infusion of drug A, drug A+B, washout B, and drug A+C (**p<0.01, paired t-test, 7 cells from 6 mice. Drug A: osteocalcin, 10ng/ml; drug B: Kcnk13 blocker, gadolinium chloride, 150uM; drug C: Kcnj3 blocker, ifenprodil,15uM). **(O-P)** Sample traces and quantification of firing rate in Type-II VTA GABAergic neurons after infusion of drug A, drug A+C, washout C, drug A+B (***p<0.001 paired t-test, 11 cells from 6 mice. Drug A: osteocalcin,10ng/ml; drug B: Kcnk13 blocker, gadolinium chloride, 150uM; drug C: Kcnj3 blocker, ifenprodil,15uM).

**Table 1.**
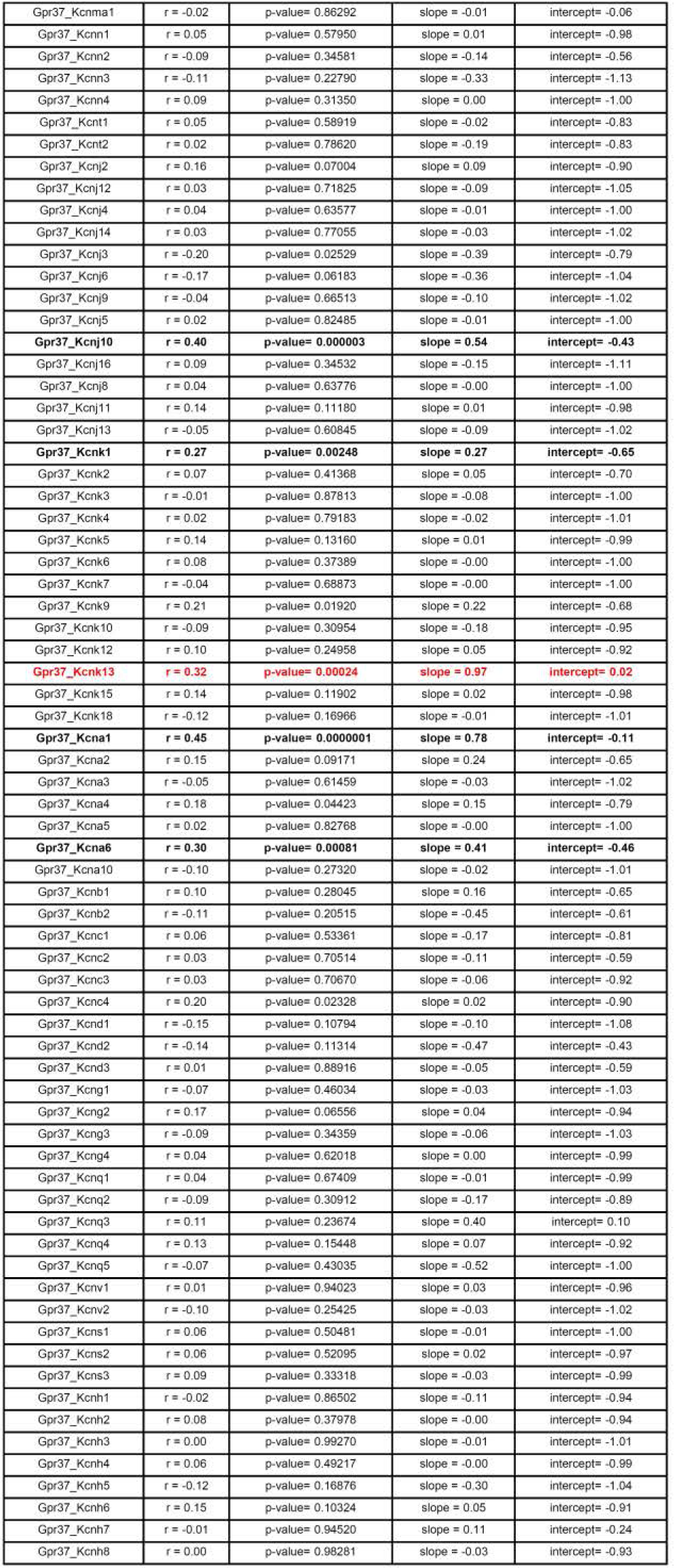
Pearson correlation of expression in the UMAP of each potassium channel with *GPR37*, together with p-value and the slope and intercept of linear regression, related to Figure 6.

**Table 2.**
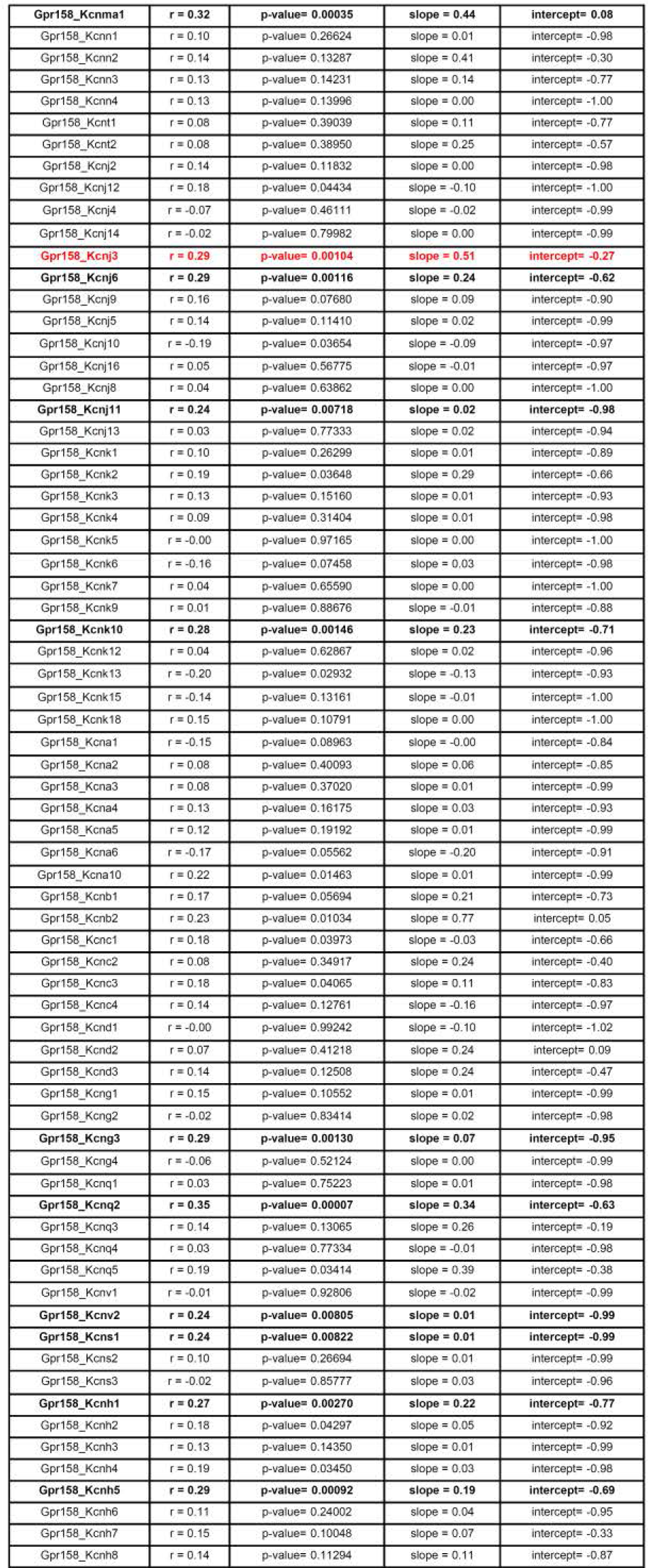
Pearson correlation of expression in the UMAP of each potassium channel with *GPR158*, together with p-value and the slope and intercept of linear regression, related to Figure 6.

Three criterions were used to identify the candidate potassium channels that most likely work downstream of the two GPRs based on these metrics: 1. The channel gene has positive correlation with the GPR; 2. P-value of correlation is lower than 0.01; 3. The slope of regression is closest to 1, while the intercept is closest to 0. Using these criteria, we identified Kcnk13 (K_2P_13.1) and Kcnj3 as the best candidate potassium channel targets for GPR37 and GPR158, respectively (Figure 6C-F).

To map the spatial organization of these neuronal subtypes, we concurrently evaluated the expression and spatial distribution of 12 genes (including maker genes, GPCRs and ion channel genes) (Figure 6G-H). We obtained 8133 neurons and 1079 oligodendrocyte from two biological replicate slices and the *in situ* sequencing data showed 7 transcriptionally and spatially defined cell clusters. Consistent with snRNA-seq result, we defined 3 GABA-Glut clusters (Figure 6I-J). And the expression level of *Qk*, the subtype 2 marker gene, is higher in GPR37+KCNK13+GAD2+ cells than in GPR158+KCNj3+GAD2+ cells (Figure 6K).

To test the causal relationship between changes in neuronal excitability and potassium current, we used specific blockers for the potassium channels that we identified in two sets of slice electrophysiology experiments on VTA GABAergic neurons (Figure 6L). In the first set, we first identified type-I neurons that potentially express GPR37. Then we applied Kcnk13 blocker together with OCN in the solution and observed a reduction in the firing rate. Washing off with OCN caused a re-elevation of firing rate, while application with Kcnj3 blocker together with OCN did not change the OCN effect (Figure 6M, N). In the second set, we first identified Type-II neurons that potentially only express GPR158. Then we applied Kcnj3 blocker together with OCN in the solution and observed an increase in the firing rate. Washing off with OCN caused a re-reduction of firing rate, while application with Kcnk13 blocker together with OCN did not change the OCN effect (Figure 6O, P). Moreover,

GPR37 is required for reduction of cAMP level ^43–47^ that is essential for rapid escape response (Figure S11A-E). Thus, GPR37 increases excitability of VTA GABAergic neurons by cAMP reduction via non-canonical Gi pathway and subsequent suppression of K_2P_13.1 encoded by *Kcnk13* (Figure S11F). This is consistent with previous finding that suppression of K_2P_13.1 led to increase of VTA neuron excitability ^48,49^. Our results also argued that GPR158 decreases excitability of VTA GABAergic neurons via Kcnj3. And these Type-I and Type-II neurons likely work in parallel pathways because the channel blockers did not exhibit crossing-effect. In summary, OCN increases excitability of GPR37- expressing VTA GABAergic neuron subpopulation by suppressing K_2P_13.1 to prime rapid visual escape response.

### HM4Di activation leads to dichotomous effect on excitability of VTA GABAergic neurons and promotes rapid visual escape

Since we identified GPR37 regulates neuronal excitability via non-canonical Gi pathway, we wondered whether G(i/o)-DREADDs (HM4Di) activation can lead to excitation of VTA GABAergic neurons. In our model, we hypothesized two signaling pathways upon G(i/o)-DREADD activation. One is the canonical pathway in which clozapine-N-oxide (CNO)-HM4Di signaling causes inhibition of neuronal activity by opening G protein inwardly rectifying K+ channels (GIRK) (Figure 7A). The other is non-canonical pathway that inhibits the production of cAMP, leading to decrease of potassium current that evoked neuronal excitation (Figure 7B).

**Figure 7.**
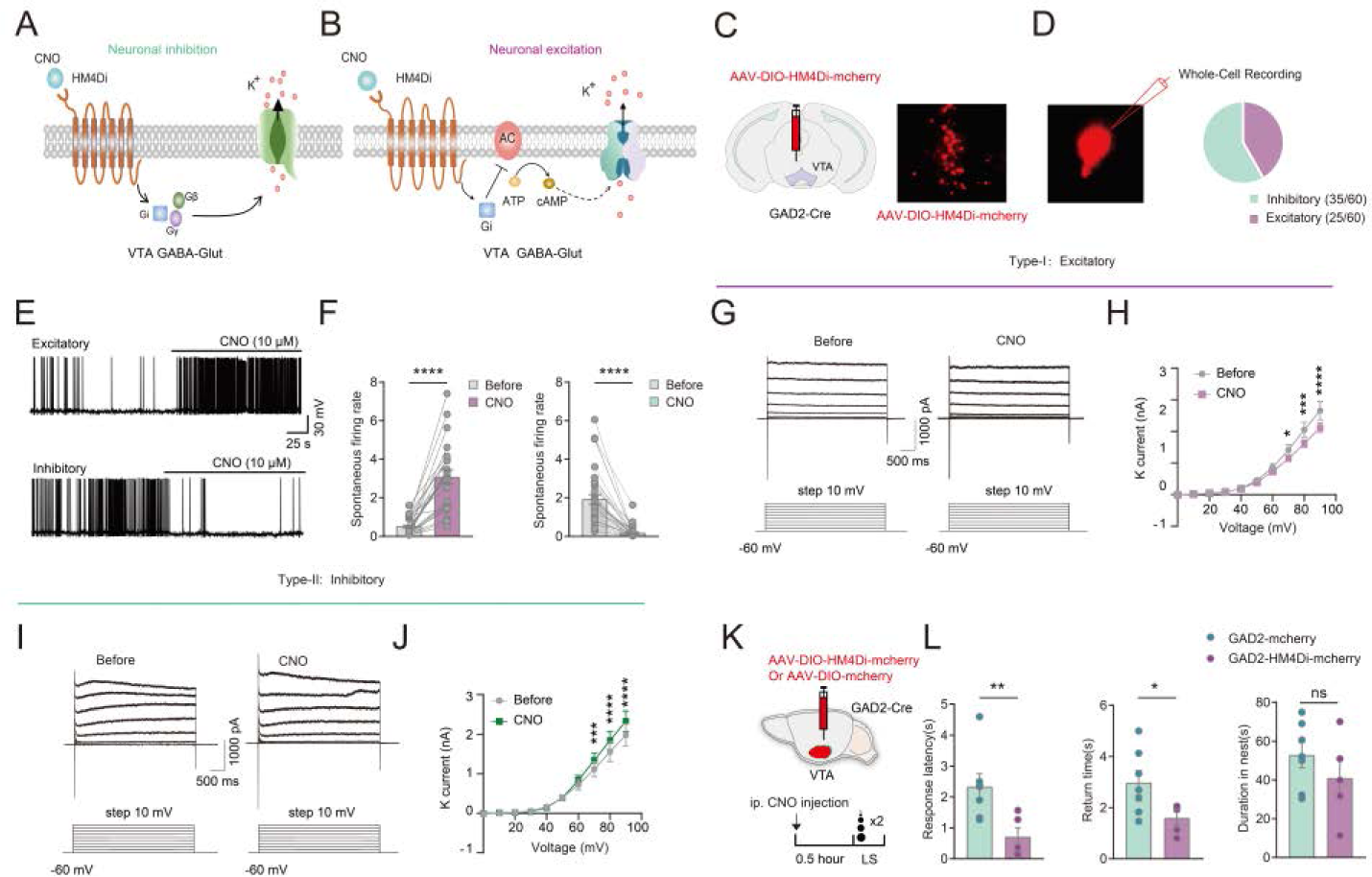
G(i/o)-DREADDs activation leads to neuronal excitation in VTA GABAergic neuron subpopulation and promoted rapid escape. **(A-B)** Model of G(i/o)-DREADDs signaling leading to inhibitory or excitatory effect. **(C)** Schematics of virus injection and validation of expression in VTA. **(D)** Example image showing HM4Di-mcherry expression in VTA GABAergic neuron. And proportion of neurons showing excitatory (Type-I, n=25) and inhibitory (Type-II, n=35) effect through whole-cell patch-clamp recording. **(E)** Example traces of action potentials upon CNO activation of HM4Di in VTA GAD2 positive neurons. **(F)** Quantification of firing rate recorded in HM4Di-expressing VTA GAD2 positive neurons treated by CNO (10μM) (****p<0.0001, paired t-test. Left: 25 cells, right: 35 cells, all from 27 mice). **(G-H)** Example traces and quantification showing decreased K+ currents in Type-I neurons upon CNO application. (Two-way ANOVA multiple comparisons, *p=0.0458, ***p=0.0003, ****p<0.0001, 6 cells from 6 mice) **(I-J)** Example traces and quantification showing increased K+ currents in Type-II neurons upon CNO application. (Two-way ANOVA multiple comparisons, ***p=0.0009, ****p<0.0001, 10 cells from 6 mice). **(K)** Experimental design. **(L)** Bar graph showing looming response in mice with (n=7) and without (n=5) HM4Di activation in VTA GABAergic neurons. Left: response latency (Two-tailed paired t test, **p=0.0032); middle: return time (Two-tailed paired t test, *p=0.0263); right: duration in the nest (Two-tailed paired t test, p>0.05).

To test these hypotheses, we conducted whole-cell recordings for VTA GABAergic neurons in acute brain slices. GAD2-ires-Cre mice were injected with AAV-DIO-HM4Di-mcherry in the VTA, and specific virus expression were validated in VTA (Figure 7C, D). Some 58% of recorded VTA GABAergic neurons (35 out of 60) showed suppressed neuronal firing, while the other 42% exhibited accelerated neuronal firing during CNO perfusion (10 μM) in the brain slices (Figure 7E, F). In addition, we also observed a decrease in K+ current in CNO-excited neurons (Figure 7G, H), and increase in K+ current in CNO- inhibited neurons (Figure 7I, J), consistent with model 1 and 2, respectively. These results mirrored excitability and corresponding K+ current changes of Type-I and Type-II neurons upon OCN infusion. Furthermore, HM4Di activation in VTA GABAergic neurons by CNO did not suppress rapid visual escape, as one would expect from the inhibitory effect of this chemogenetic GPCR. Instead, HM4Di activation promoted rapid visual escape (Figure 7K, L), consistent with the result that the excitability of a subset of VTA GABAergic neurons were elevated by HM4Di activation (Figure 7E, F), and our previous finding that activation of VTA GABAergic neurons evokes escape response ^12^. Taken together, G(i/o)-DREADD activation leads to excitation of VTA GABAergic neuron subpopulation and accelerated visual escape response.

## Discussion

In this study, we discovered a novel role for bone-derived protein OCN, which primes rapid visual escape by binding to GPR37 receptor in a subpopulation of VTA GABAergic neurons through a non-canonical GPCR signaling mechanism. This regulatory mechanism underscores a finely tuned brain-body interaction emerged during vertebrate evolution, which increases an animal’s survival in dangerous or unpredictable environments.

### OCN plays a permissive role on visual escape by modulating neuronal excitability in VTA

Our results demonstrated that osteocalcin can directly increase neuronal excitability through GPR37 receptor in a subset of GABAergic neurons by decreasing K+ current via THIK-1 channels and reducing cAMP levels. THIK- 1, belonging to two-pore domain K^+^ channels family, is responsible for maintaining the resting membrane potential (RMP) by facilitating the efflux of potassium ions^50–52^. Notably, previous study showed that rapid visual escape relies on instantaneous activation of VTA GABAergic neurons upon the presentation of looming stimulus^12^. Our new results indicated that OCN-GPR37 signaling decreased RMP to ensure that these neurons are in a state of readiness to rapidly respond to visual threats (Figure S6B).

Notably, the way by which OCN increases neuronal excitability is distinguished from classical neuromodulators like dopamine or choline, whose levels change acutely and dynamically upon environmental stimulation and animal behavior ^20,53,54^. Given the fact that VTA neurons respond to looming stimulus within tens of milli-seconds and the OCN is derived from the bone and get into the brain by passing through BBB, the OCN levels cannot increase before VTA neurons respond to looming. Thus, instead of playing an active role in modulating neuronal activity as classical neuromodulators, OCN plays a permissive role in maintaining heightened excitability of VTA GABAergic neuron subpopulation, making them ready to respond to sudden visual cues that reflect rapid approaching of predator.

The above mechanism underscores the importance of situational adjustments and long-term preparatory states in maintaining adaptive responses to environmental challenges. OCN does not directly drive escape (Fig 2F), but enhances the efficiency of transforming visual input into motor outputs, optimizing this process at the critical VTA hub that receives visual threat signal from SC and transmits to downstream motor regions. The effect of OCN on VTA GABAergic neurons may tune their adaptive responses to various stimuli and conditions, including predator, prey, and stress stimulus, providing insights into the relationship between bone-brain interactions and mental health. Elevated Kcnk13 levels in postmortem brains of Alzheimer’s and Parkinson’s patients^55^ underscore the therapeutic potential of targeting Kcnk13, as OCN does in specific VTA GABAergic subpopulation, highlighting the broader significance of our findings in understanding and treating neurological diseases.

### Bidirectional control of circuit activity by OCN

Our snRNA-seq results showed that over 70% of VTA GABAergic neurons express both GABAergic and glutamatergic markers, and potentially co-release these two neurotransmitters, highlighting the functional heterogeneity of VTA GABAergic neurons. Our findings demonstrates that GPR37 associates with THIK-1 channel, exhibiting excitatory effects on OCN signaling, whereas GPR158 associates with G protein-gated inwardly rectifying potassium GIRK1 channel (also known as Kir3.1) encoded by Kcnj3 gene, regulating inhibitory effect to OCN. We identified GPR37’s role in VTA GABA neurons, through which it facilitates the instant, energy-demanding escape via OCN signaling. The GPR158 receptor, recently serves as a metabotropic glycine receptor, has been implicated as a mediator of osteocalcin’s regulation of hippocampus-dependent memory and stress-induced depression ^56–60^. We found that GPR158 is expressed in virtually all GABAergic neurons, whereas GPR37 is only expressed in a subpopulation of these neurons. We demonstrate that the GPR158 receptor, known to associate with the RGS7-Gβ5 complex, interacts with the GIRK1 channel, which is distinctly expressed in VTA non-DA neurons, unlike the GIRK2/3 channels in VTA dopamine neurons^61^. Consistent with the modulation of GIRK channels by the RGS7-Gβ5 complex^62,63^, this GPR158-GIRK1 interaction leads to an inhibitory response to OCN, in contrast to the regulation of GABAB receptor-GIRK2/3 coupling in VTA dopamine neurons by RGS2^61^. Our finding indicates that the OCN-GPR158 signaling pathway may serve as a critical hub, coordinating GIRK1 channel, may mediate OCN’s long-term effects, enabling environmental recall and stress adaptation. Notably, our electrophysiology experiments suggests that the GPR37+ and GPR37- neurons are likely to be in two parallel circuits, since the channel specific blockers did not show crossing effects. Thus, it is possible that OCN displays bidirectional control on activity of distinct GABAergic circuits to differentially regulate distinct brain functions and behaviors.

Our findings that OCN bidirectionally modulates neuronal excitability adds to the growing body of evidence indicating that peripheral factors, such as glucocorticoids, insulin, and thyroid hormones, can regulate the activity of ion channels and neurotransmitter systems in the brain, thereby shaping behavioral and physiological responses to stress and environmental stimuli ^64–66^. The intricate crosstalk between the periphery and the central nervous system, as exemplified by our findings on the role of bone-derived OCN in modulating neuronal excitability, underscores the importance of adopting a holistic perspective of body-brain interaction to fully elucidate the physiological mechanisms governing innate defensive behaviors and their dysregulation in fear and anxiety disorders.

### Excitatory effect on VTA GABAergic subpopulation by inhibitory DREADD

Classically, G(i/o)-coupled receptors, such as hM4Di, are known for their inhibitory roles by activating GIRK channels ^67–69^. However, our study reveals a surprising excitatory effect of the Gi/o-DREADD on a subpopulation of VTA GABAergic neurons that further promote rapid visual escape. This finding highlighted the sophisticated regulation of excitability in VTA and demonstrated the importance of taking cellular heterogeneity within a cell type into account when applying chemogenetic approaches to study circuit basis for behavior. Besides GPR37, GPR158 and corresponding potassium channel heterogeneity within VTA GABAergic neurons, other mechanisms remain to be discovered that play similar roles in dichotomously regulating neuronal electrophysiology within particular neuronal types.

### Evolution of body-brain interactions for sensorimotor transformation

Our findings revealed how skeletal cues shape neural circuitry and behavior, offering a framework for brain-body integration. The effect of OCN on enhancing VTA GABAergic neuron excitability exemplifies the sophisticated regulation linking body and brain and could be an evolutionary strategy for rapid threat responses. The co-evolution of skeletal and visual systems in vertebrate ^70,71^underscores the importance of whole-body context for studying neural circuits and behaviors. Indeed, besides VTA, there are lots of other relay stations of sensory-motor transformation in the brain. It is possible that peripherally derived factors play a wide-spread role in regulating the activity of these relay stations, given the fact that many brain structures and body organs are co-evolved in vertebrates.

### Limitations of the study

While we showed VTA GABAergic neurons regulated by OCN-GPR37 mediate visual escape, the cell-type specific downstream synaptic target neurons remain further investigation. Additionally, the precise expression patterns of OCN receptors in other brain areas require further characterization. Integrating circuit-level and spatial transcriptomic approaches can address these limitations, allowing a more comprehensive understanding on the role of OCN and its receptors in regulating neural circuit function.

## Author Contributions

Conceptualization, X.L., X.M.L. and L.W.; methodology, X.M.L., X.L., L.T., J.L., X.G., L.N.W; Investigation, X.M.L., J.L., L.T., X.G., L.N.W., Y.L.H., G.L., S.M., B.F., L.Y., Z.Q.; writing-original draft, X.M.L., writing-review and editing, X.M.L., L.T ., X.L and L.W. funding acquisition, L.W., X.M.L., X.L. and L.T.; resources, L.W. and X.L.; supervision, XM.L., X.L., L.W. and L.T.

## Acknowledgments

We thank Dr. Helmut Kettenmann, Dr. Ming-Hu Han and Dr. Qiu-Fu Ma for comments on our manuscript. This work was partially sponsored by National Natural Science Foundation of China (32230042 to L.W., 32371069 to X.M.L., 31930047 to L.W.,32371019 to L.T.); Financial Support for Outstanding Talents Training Fund in Shenzhen (L.W.); STI2030-Major Projects-2022ZD0211700; Ten Thousand Talent Program(L.W.); Chang Jiang Scholars Program (L.W.); International Partnership Program of Chinese Academy of Sciences (CAS) 172644KYS820170004; CAS Key Laboratory of Brain Connectome and Manipulation (2019DP173024); the Strategic Priority Research Program of CAS (XDB32030100); the Guangdong Special Support Program; Key-Area Research and Development Program of Guangdong Province 2018B030331001; KQTD20210811090117032 to X.L., National Key R&D Program of China to L.T. (2022YFA1105503)

## STAR*METHODS

### KEY RESOURCES TABLE

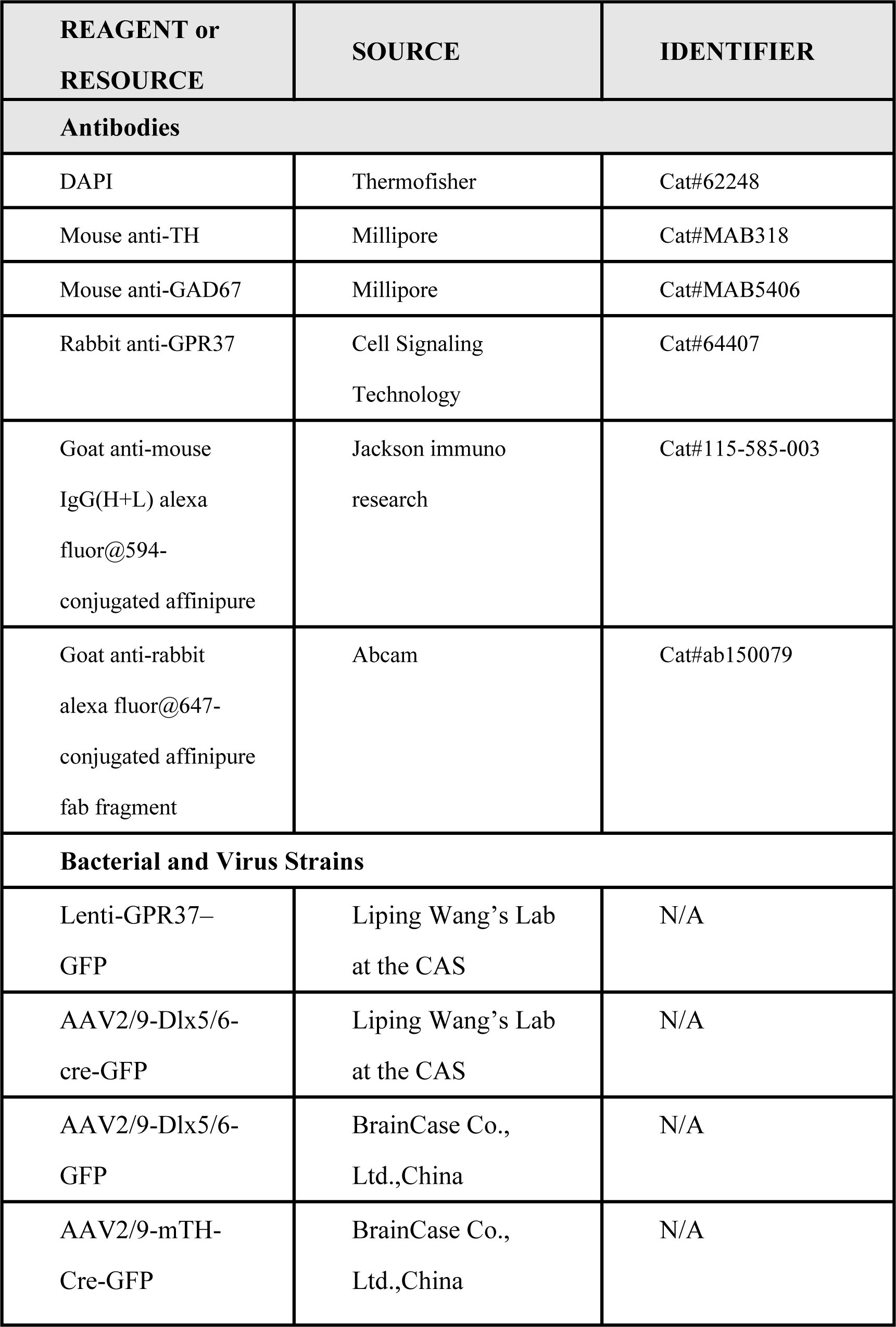
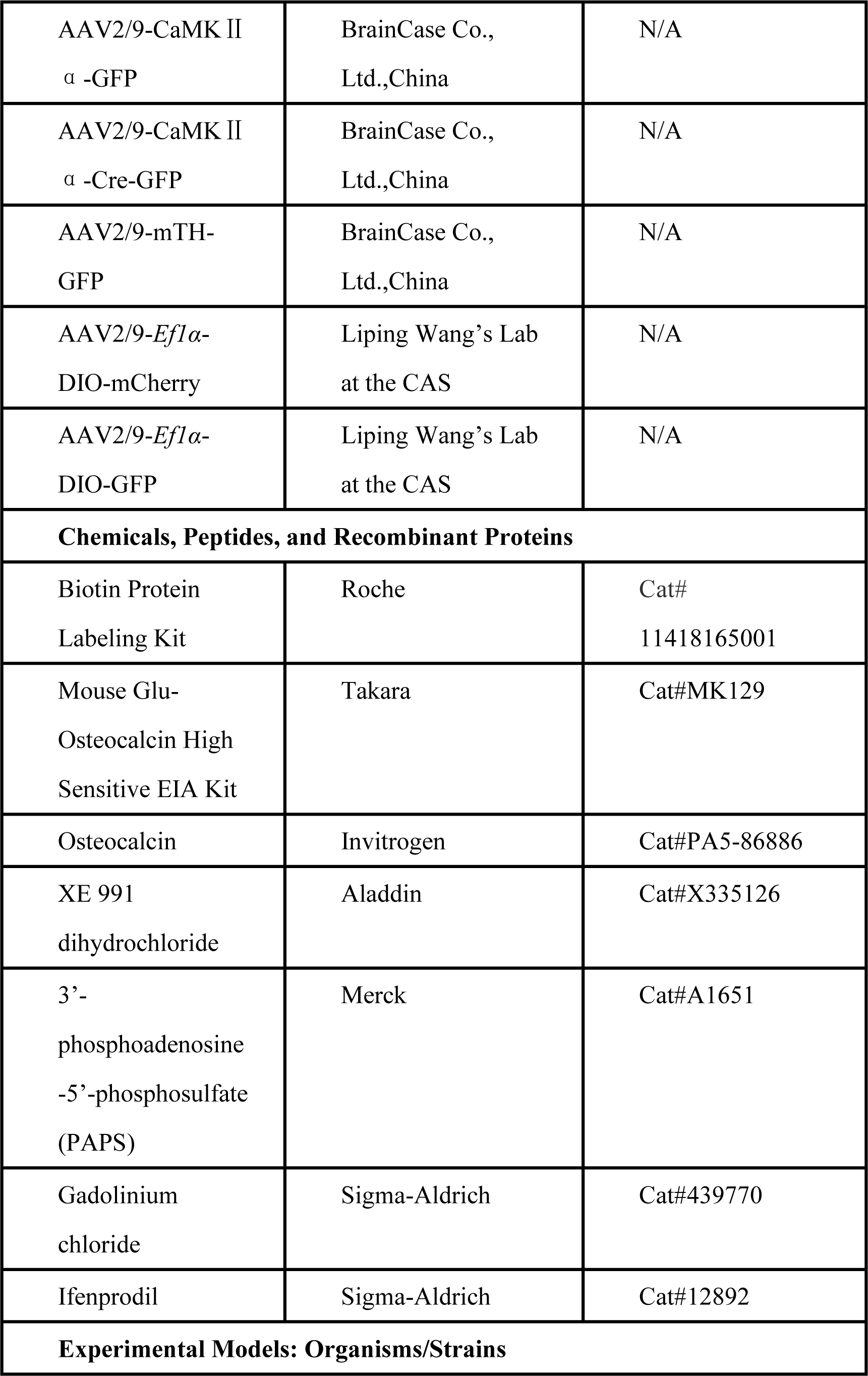
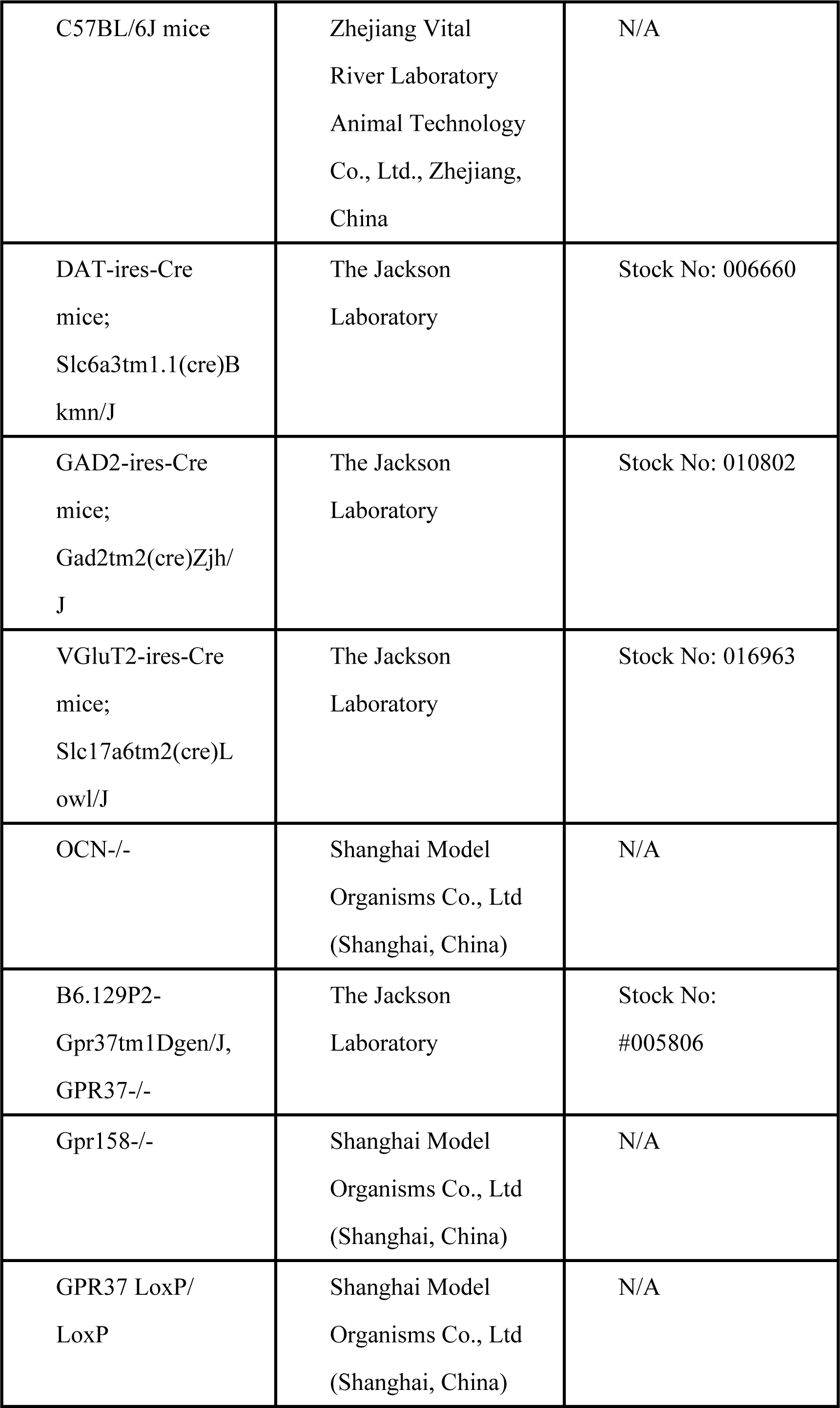
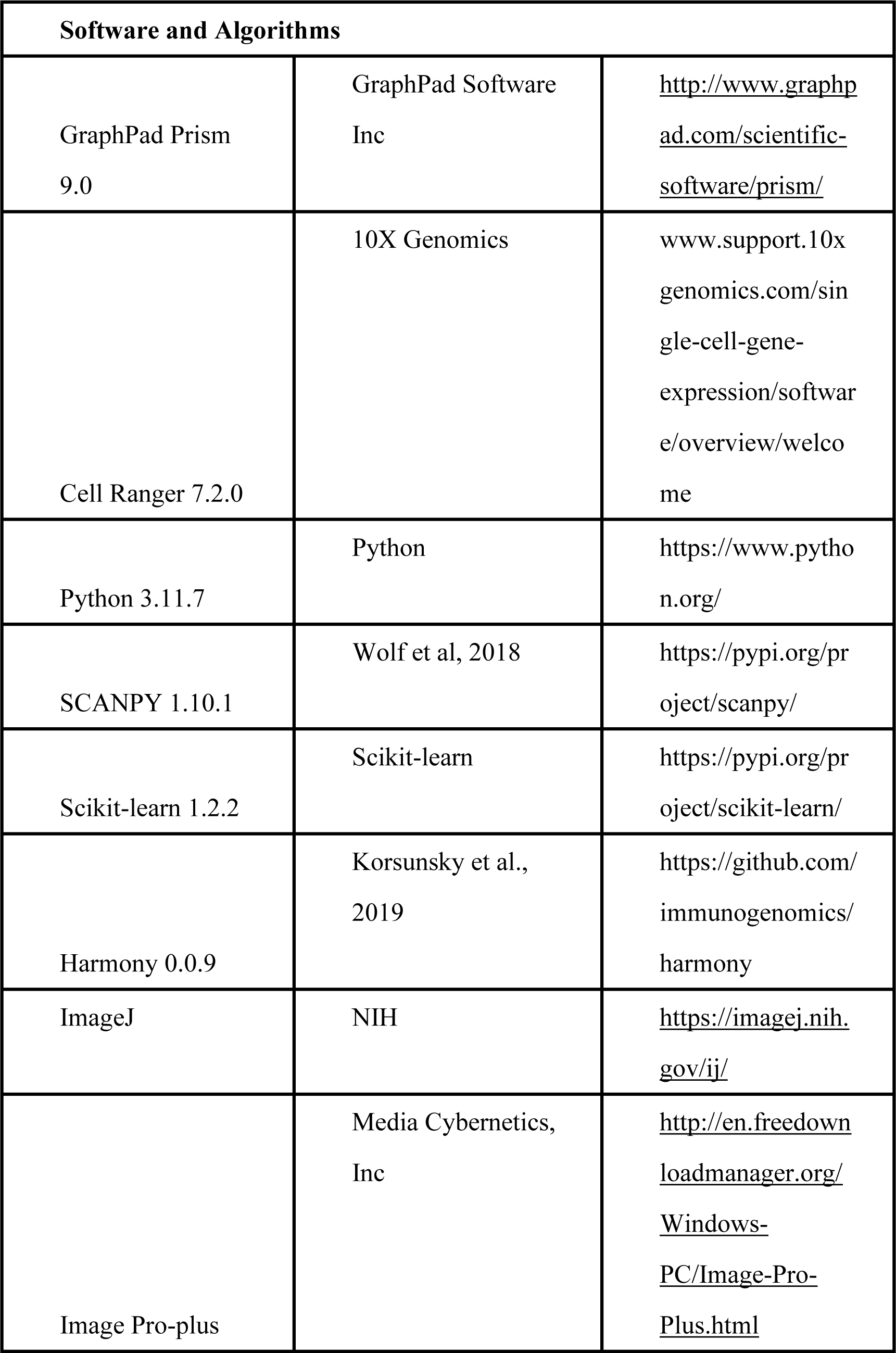
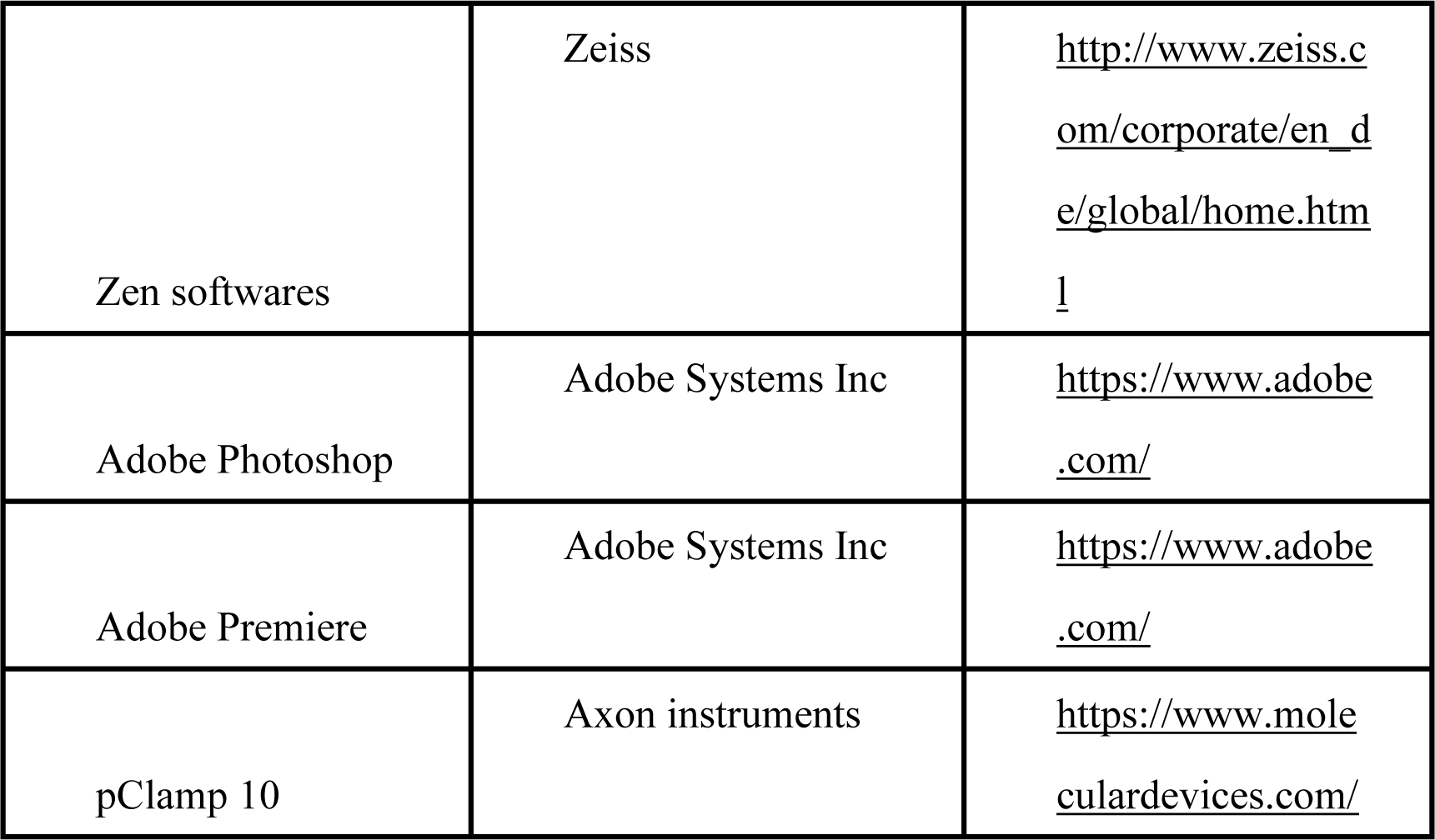

| **REAGENT or RESOURCE** | **SOURCE** | **IDENTIFIER** |
| --- | --- | --- |
| **Antibodies** |  |  |
| DAPI | Thermofisher | Cat#62248 |
| Mouse anti-TH | Millipore | Cat#MAB318 |
| Mouse anti-GAD67 | Millipore | Cat#MAB5406 |
| Rabbit anti-GPR37 | Cell Signaling Technology | Cat#64407 |
| Goat anti-mouse IgG(H+L) alexa fluor@594-conjugated affinipure | Jackson immuno research | Cat#115-585-003 |
| Goat anti-rabbit alexa fluor@647-conjugated affinipure fab fragment | Abcam | Cat#ab150079 |
| **Bacterial and Virus Strains** |  |  |
| Lenti-GPR37–GFP | Liping Wang’s Lab at the CAS | N/A |
| AAV2/9-Dlx5/6-cre-GFP | Liping Wang’s Lab at the CAS | N/A |
| AAV2/9-Dlx5/6-GFP | BrainCase Co., Ltd.,China | N/A |
| AAV2/9-mTH-Cre-GFP | BrainCase Co., Ltd.,China | N/A |
| AAV2/9-CaMK II<br>$\alpha$ -GFP | BrainCase Co.,<br>Ltd.,China | N/A |
| AAV2/9-CaMK II<br>$\alpha$ -Cre-GFP | BrainCase Co.,<br>Ltd.,China | N/A |
| AAV2/9-mTH-GFP | BrainCase Co.,<br>Ltd.,China | N/A |
| AAV2/9-*Efl*α-<br>DIO-mCherry | Liping Wang's Lab<br>at the CAS | N/A |
| AAV2/9-*Efl*α-<br>DIO-GFP | Liping Wang's Lab<br>at the CAS | N/A |
| **Chemicals, Peptides, and Recombinant Proteins** |  |  |
| Biotin Protein<br>Labeling Kit | Roche | Cat#<br>11418165001 |
| Mouse Glu-Osteocalcin High<br>Sensitive EIA Kit | Takara | Cat#MK129 |
| Osteocalcin | Invitrogen | Cat#PA5-86886 |
| XE 991<br>dihydrochloride | Aladdin | Cat#X335126 |
| 3'-<br>phosphoadenosine<br>-5'-phosphosulfate<br>(PAPS) | Merck | Cat#A1651 |
| Gadolinium<br>chloride | Sigma-Aldrich | Cat#439770 |
| Ifenprodil | Sigma-Aldrich | Cat#12892 |
| **Experimental Models: Organisms/Strains** |  |  |
| C57BL/6J mice | Zhejiang Vital<br>River Laboratory<br>Animal Technology<br>Co., Ltd., Zhejiang,<br>China | N/A |
| DAT-ires-Cre<br>mice;<br>Slc6a3tm1.1(cre)B<br>kmn/J | The Jackson<br>Laboratory | Stock No: 006660 |
| GAD2-ires-Cre<br>mice;<br>Gad2tm2(cre)Zjh/<br>J | The Jackson<br>Laboratory | Stock No: 010802 |
| VGluT2-ires-Cre<br>mice;<br>Slc17a6tm2(cre)L<br>owl/J | The Jackson<br>Laboratory | Stock No: 016963 |
| OCN-/- | Shanghai Model<br>Organisms Co., Ltd<br>(Shanghai, China) | N/A |
| B6.129P2-<br>Gpr37tm1Dgen/J,<br>GPR37-/- | The Jackson<br>Laboratory | Stock No:<br>#005806 |
| Gpr158-/- | Shanghai Model<br>Organisms Co., Ltd<br>(Shanghai, China) | N/A |
| GPR37 LoxP/<br>LoxP | Shanghai Model<br>Organisms Co., Ltd<br>(Shanghai, China) | N/A |

| Software and Algorithms |  |  |
| --- | --- | --- |
| GraphPad Prism<br>9.0 | GraphPad Software<br>Inc | http://www.graphpad.com/scientific-software/prism/ |
| Cell Ranger 7.2.0 | 10X Genomics | www.support.10xgenomics.com/single-cell-gene-expression/software/overview/welcome |
| Python 3.11.7 | Python | https://www.python.org/ |
| SCANPY 1.10.1 | Wolf et al, 2018 | https://pypi.org/project/scanpy/ |
| Scikit-learn 1.2.2 | Scikit-learn | https://pypi.org/project/scikit-learn/ |
| Harmony 0.0.9 | Korsunsky et al.,<br>2019 | https://github.com/immunogenomics/harmony |
| ImageJ | NIH | https://imagej.nih.gov/ij/ |
| Image Pro-plus | Media Cybernetics,<br>Inc | http://en.freedownloadmanager.org/Windows-PC/Image-Pro-Plus.html |
| Zen softwares | Zeiss | http://www.zeiss.com/corporate/en_de/global/home.html |
| Adobe Photoshop | Adobe Systems Inc | https://www.adobe.com/ |
| Adobe Premiere | Adobe Systems Inc | https://www.adobe.com/ |
| pClamp 10 | Axon instruments | https://www.moleculardevices.com/ |

## METHOD DETAILS

### Subjects

All experimental procedures were approved by the Animal Care and Use Committees at the Shenzhen Institute of Advanced Technology (SIAT), Chinese Academy of Sciences (CAS). Adult (6 to 8 weeks-old) male C57BL/6 (Zhejiang Vital River Laboratory Animal Technology Co., Ltd., Zhejiang, China), GAD2- *ires*-Cre (Jax No. #010802) mice, DAT-*ires*-Cre (Jax No. #006660), VGluT2- ires-Cre (Jax No. #016963), GPR37^-/-^ mice (B6.129P2-Gpr37tm1Dgen/J, Jax No. #005806) were used in this study. The OCN^-/-^ and GPR37 ^LoxP/LoxP^ mice were generated using CRISPR-Cas9 technique by Shanghai Model Organisms Co., Ltd (Shanghai, China). GPR37cKO and control animals (GPR37^flox/flox^) were obtained from crossed between DAT-*ires*-Cre animals; GPR37flx/flx and DAT- *ires*-Cre animals. All animals were housed at 22–25 °C on a circadian cycle of 12-hour light and 12-hour dark with ad-libitum access to food and water.

### Viral vector preparation

For specific VTA recovery GPR37, we used plasmids for lenti-viruses encoding GPR37–GFP, were packaged by Liping Wang Lab, Shenzhen. Viral vector titers were in the range of 3-6x10^12^ genome copies per ml (gc)/mL.

For specific neuron types knock out GPR37 receptor and anatomical labeling, AAV2/9 virus encoding mTH-cre-EGFP and the control virus mTH-EGFP were all packaged by BrainCase Co., Ltd., Shenzhen, and Dlx5/6-cre-GFP and the control virusDlx5/6-GFP, Ef1α-DIO-mcherry were all packaged by Liping Wang Lab, Shenzhen. Adeno-associated were purified and concentrated to titers at approximately 3×10^12^ v.g/ml and 1×10^9^ pfu/ml, respectively.

### Stereotaxic Surgery and virus injection

Stereotaxic surgeries were performed as described previously (Zhou et al., 2019). Briefly, Animals were anesthetized with pentobarbital (i.p., 80 mg/kg) and then positioned in a stereotaxic apparatus (RWD, China). Injections were conducted by a syringe (Hamilton #65460-05, 10ul ) under the control of a micro syringe pump (KDS, USA). The injection rate was 100nl/min. 10min after the end of virus injection, the syringe was retracted slowly to avoid backflow.

The coordinates of virus injection sites was used for virus injection: VTA (AP, – 3.20 mm, ML, ±0.25 mm, and DV, –4.4 mm) .

### Looming test

The looming test was performed as described previously (Zhou et al, 2019). The looming paradigm consisted of a 40ⅹ40ⅹ30 cm closed Plexiglas box and with a shelter nest in the corner. An LCD monitor was placed on the ceiling to present upper visual field looming stimulus, which was a black disc on a grey background expanding from a 2° to 20° visual angle, repeated 15 times, lasting 5.5 seconds. Behaviors were recorded using a Sony FDR-AX45 camera. Mice were handled and habituated for 10 min to the looming box the day before the test. During looming tests, mice was allowed freely explore the looming box for 3 minutes, and the looming stimuli were presented for 2 trials with the inter-trial intervals no less than 3 minutes.

### In vivo Treatments

For measurement of circulating bioactivated osteocalcin, blood was collected from orbital vessels after the 3 months old and 7 months old WT mice were anesthetized with sodium pentobarbital (80 mg/kg). Osteocalcin levels were quantified using the following kit: Mouse Glu-Osteocalcin High Sensitive EIA (ELISA) Kit. MK-129 (Takara, USA). All measurements were conducted following the manufacturer instructions.

To test the innate defensive response to looming of old mice with administration of osteocalcin, the 7 months old WT mice were injected intraperitoneally (i.p.) with either vehicle or osteocalcin (50ng/g, PA5-86886, Invitrogen) every day for 10 days (Qian et al., 2021).

### Cannula implants and pharmacology

OCN-/- mice were used for pharmacological experiments. The cannulas were implanted 0.2 mm above the VTA (AP, –3.20 mm, ML, ±0.25 mm, and DV, –4.2 mm). Mice were allowed to recover for at least 2 weeks before experiments. After undergoing handle and habituation protocol, osteocalcin was infused into the drug cannulas 0.5 hour before a looming test to assess effect on visual predatory stress in VTA recovery osteocalcin in OCN-/- mice.

### Electrophysiological recordings

Acute brain slices of VTA were prepared as done in previous studies. Male C57BL/6J and GPR37-/- mice were anaesthetized with isoflurane and perfused immediately with ice-cold aCSF (artificial cerebrospinal fluid), which contained (in mM): 125 NaCl, 2.5 KCl, 1.3 NaH2PO4, 1.3 Na-ascorbate, 0.6 Na-pyruvate, 2 CaCl2, 2 MgSO4, 10 glucose, 25 NaHCO3 (oxygenated with 95% O2 and 5% CO2, pH 7.35, 295–305 mOsm). Acute brain slices (270 µm) containing the VTA were cut using a cutting solution containing (in mM): 110 Choline Chloride, 2.5 KCl, 1.3 NaH2PO4, 1.3 Na-ascorbate, 0.6 Na-pyruvate, 0.5 CaCl2, 7 MgCl2, 25 glucose, 25 NaHCO3, and saturated by 95% O2 and 5% CO2. Slices were maintained in the holding chamber for 1 hour at 34°C. Slices were transferred into a recording chamber fitted with a constant flow rate of aCSF equilibrated with 95% O2/5% CO2 (2.5 ml/min) and at 34°C. Glass microelectrodes (2-4 MΩ) filled with an internal solution containing (mM): 130 K-gluconate, 10 KCl, 10 HEPES, 1 EGTA, 0.3 Na-GTP, 2 Mg-ATP, 2 MgCl2 (pH 7.2, 285 mOsm).

Electrophysiological properties were evaluated from whole-cell recordings. Firing rate was recorded with whole-cell clamp configuration. To isolate voltage- gated K+ channel-mediated currents, 4 s pulses with 10 mV step voltages from 0 to +100 mV at –60 holding potential. Cell excitability was measured with 1 s incremental steps of current injections (25, 50, 75 and 100 pA). Series resistance was monitored during the experiments and membrane currents and voltages were filtered at 3 kHz (Bessel filter). Data acquisition was collected using a Digidata 1440A digitizer and pClamp 10.2 (Axon Instruments).

### Tissue dissection from adult VTA

Mice were anesthetized with pentobarbital (i.p., 80 mg/kg), and brains were rapidly removed and put in the ice cold DEPC-PBS. Brains were cut into 1 mm thick coronal sections on mouse brain matrix (RWD, China). The sections containing the VTA (ranging from -2.92 to -3.88 anterior posterior to Bregma) were micro-dissected from neighboring brain regions (n=20 mice). Every dissected VTA tissue were collected in ice cold centrifuge tubes and then transferred to vacuum cup filled with liquid nitrogen. Total dissected VTA were stored at -80°C for single nucleus RNA-seq processing and western blotting.

### Single cell nucleus RNA-seq

Dissected VTA tissues were collected and dissociated using an adult brain dissociation kit from Miltenyi Biotec (Bergisch Gladbach, Germany). Subsequent procedures were performed by AccuraMed (Shanghai, China). Briefly, single-cell gel beads in emulsions (GEMs) and Barcoding were firstly generated. Master Mix were prepared, and Single-Cell suspension ran in the Chromium Controller (10X Genomics). GEMs transferred and Post GEM-RT cleanup& cDNA amplification conducted following standard protocols. Gene expression library were constructed, and every library was sequenced on a HiSeq X Ten platform (Illumina).

### Bioinformatic analysis

#### Initial processing of snRNA-seq data

Using the official analysis software CellRanger (version 3.1.0) provided by 10x Genomics, the raw data was filtered, mapped, quantified, and identified and recovered cells, and finally the gene expression matrix of each cell was obtained. The specific implementation method are as follows: Extract 10x Barcode and UMI sequence at the R1 end, as well as the insert sequence at the R2 end for gene alignment. Use STAR aligner (Spliced Transcripts Alignment to a Reference) to map reads (R2 insert of 91bp extracted in the previous step) to the reference genome (GRCm38), and use genomic GTF annotation files to correct to distinguish exon regions, intron regions, and intergenic regions. The Barcode sequence information obtained by sequencing is compared with the known Barcode sequence in the database, and the barcode sequence (measured sequence) that exactly matches the known Barcode in the database is the real sequence. The sequenced UMI sequence can not used directly for subsequent analysis, and unqualified reads need to be filtered and corrected. Count all valid barcodes to obtain an unfiltered blast gene expression matrix, only read that contains valid barcodes and UMIs and are reliably aligned can be counted. A distinction between barcodes containing cells and background barcodes is required to extract formal single-cell data for downstream analysis.

#### Data QC and Filtering

We generated scatter plots of the number of transcript molecules per cell (n_counts), the percentage of transcripts from mitochondrially encoded genes (percent_mito), and the number of expressed genes (n_genes) to identify outlier cells. Cells meeting the following criteria were retained: 100 < n_genes < 8000, percent_mito < 1%, and n_counts < 40,000. Only genes detected in more than three cells were kept for further analysis. We used scanpy.external.pp.scrublet() to identify doublets, setting the doublet score threshold to 0.15 and retaining cells with a score below this threshold, labeled as ’False’. Cells were normalized for library size differences, with transcript counts in each cell rescaled to sum to 10,000, followed by log-transformation.

#### Merging Two Bio-replicates and Cell Class Annotation

We used scanpy.AnnData.concatenate() to merge two bio-replicates, identified highly variable genes (HVGs), and computed a reduced dimensional representation of the data using principal component analysis (PCA). Harmony was then applied to adjust the principal components, storing the result as ’X_harmony’. This representation was used to compute a nearest-neighbor graph of the cells, which was subsequently clustered using the Leiden algorithm and embedded in 2D via the Uniform Manifold Approximation and Projection (UMAP) algorithm.

After that, neurons and non-neurons were classified based on canonical marker genes, with the Leiden algorithm resolution parameter set to 0.5. Neurons were then re-ustered with the resolution parameter increased to 0.8. All neurons were annotated into four classes: Glut, GABA, GABA-Glut, and Dopa using canonical marker genes. Finally, based on the number of principal components from PCA that explained the most variance, the GABA-Glut cells were clustered into three clusters with a resolution set to 0.1.

The scanpy.tl.rank_genes_groups() function and the Wilcoxon rank-sum test with tie correction in the Scanpy package were then used to identify marker genes.

### Calculating Gene Expression Correlation

The GABA-Glut cells were assigned into specific bin according to the UMAP1 and UMAP2 axes with a step size of 1, and the average expression of Gpr37, Gpr158, and all K channel genes was calculated for each bin. The correlation coefficients and p-values for the expression of all K channel genes with Gpr37 and Gpr158 were computed using the spearmanr function in scipy.stats, with a p-value threshold set at 0.01. The slope and intercept of linear regression fitting were calculated using numpy.polyfit.

### In situ sequencing

The specific probes for target RNA were designed by spatial FISH Ltd. Samples were fixed by 4% paraformaldehyde, then covered with reaction chamber to perform the following reactions. After dehydration and denaturation of samples with methanol, the hybridization buffer with specific targeting probes was added to the chamber for incubation at 37℃ overnight. Then samples were washed three times with PBST, followed by ligation of target probes in ligation mix at 25℃ for 3 h. Next, samples were washed three times with PBST and subjected to rolling circle amplification by Phi29 DNA polymerase at 30℃ for overnight.

Subsequently, the fluorescent detection probes in hybridization buffer was applied to samples. Finally, samples were dehydrated with an ethanol series and mounted with mounting medium. After capturing images by Leica THUNDER Imaging Systems, signal dots were decoded to interpret RNA spatial position information.

### Image Alignment

We used ImageJ to process multi-channel z-stack images (containing z, y, x dimensions). The images were maximum projected along the z-axis to generate 2D multi-channel images, and contrast parameters were adjusted to enhance fluorescence signals. The processed results were saved in .tif format (e.g., ‘demo_A_a-gene_b-gene_c-gene.tif’). Next, we used the BigWarp plugin for image alignment. Corresponding points were selected for alignment, and the results were exported and named ‘demo_B_moving_d-gene_e-gene_f-gene.tif’.

### Fluorescence Signal Detection and Statistics

The bigwarp-calibrated tif images were input into the U-FISH system for standardized image output and automated analysis, with results saved accordingly. For dense or weak fluorescence signals, the RS-FISH plugin in ImageJ was used for manual single-channel image calling, parameter adjustment, and result saving. In U-FISH result files, axis-0 represents the channel, and axis- 1 and axis-2 represent the y and x coordinates, respectively; RS-FISH results include x, y, t, c, and intensity information. Gene information was assigned to the corresponding channels by adding a ‘gene’ column at the end of the file, and the results were merged into ‘all_gene_location.csv’.

### Cell Segmentation

In ImageFlow, after loading the required packages, the DAPI tif image was read using AICS:ReadImage, and cell nuclei were segmented using cellpose. The segmentation results were visualized using Viewer:Standalone Napari and saved as ‘mask.tif’. Then, the gene coordinate file was read using pandas:read_csv, and genes were assigned to cells using innereye:gene_to_cell, with the statistical results saved.

### Visualization Analysis

ClusterMap algorithm was used for visualization analysis in the Python environment. The coordinates of all successfully decoded gene points and the DAPI image were input. The “ClusterMap” function was used to segment points into individual cells, with parameters set to a radius of 15 and a window size of 60. Cells meeting quality standards were normalized and scaled, and batch effects were corrected using the combat function. PCA was performed using the “arpack svd_solver”, focusing on the top five principal components (PC1-PC5). Cells were clustered using the Louvain algorithm (resolution 0.6), and UMAP was generated for visualization. The “rank_genes_groups” function was used to identify differentially expressed genes (DEGs) specific to each cluster, using the Wilcoxon test method.

### Histology and immunostaining

Mice was overdosed with pentobarbital and perfused with 0.9% saline followed by 4% paraformaldehyde (PFA) in PBS. Brains were dissected and postfixed in 4% PFA at 4 °C for 24 h and transferred to 30% sucrose for 2 d. We cut 40-μm coronal slices of the entire rostrocaudal extent of the brain via a cryostat at – 15 °C and stored in 24-well plates containing cryoprotectant at 4 °C.

To visualize virus expression, cannula placements, floating sections blocked with 10% normal goat serum in PBS-T (0.3% Triton-X 100) and DAPI (1:50000, Cat#62248, Thermofisher). If needed, brains slices were kept at 4 °C overnight with the following antibodies mouse anti-TH, mouse anti-GAD67, rabbit anti- GPR37 (Cat#64407, Cell Signaling Technology). The secondary antibodies Alexa fluor 488 or Alexa fluor 594 or Alexa fluor 647 were used. Sections were mounted and covered slipped with Fluoromount aqueous mounting medium (Sigma-Aldrich, USA). Sections were then photographed by Olympus VS120 virtual microscopy slide scanning system or Zeiss LSM 880 confocal microscope. Images were analyzed with and ImageJ, Image Pro-plus, and Photoshop software.

### Neuronal culture and Biotinylated Osteocalcin treatment

Bilateral ventral tegmental area (VTA) were dissected from 2-month old mice. VTA from 5-6 mice were collected and dissociated with 0.25% Trypsin/EDTA for 30 min at 37°C. Cells were pelleted and resuspended in DMEM supplemented with 10% fetal bovine serum, penicillin/streptomycin and then plated on poly-d-lysine-coated glass coverslips in a 24-well plate at a density of 7 × 104 cells/well. Cultures were maintained for 2 hours at 37°C with 5% carbon dioxide atmosphere. Cells were then treated with biotinylated Osteocalcin (Biotin Protein Labeling Kit, Cat# 11418165001, Roche, USA) with different concentration for 30 min at 37°C.

For immunostaining, cells were fixed for 20 min in 4% paraformaldehyde. After fixation, cells were incubated in a blocking solution (3% normal goat serum in tris buffer saline solution with 0.1% Triton X-100) for 1 hour at room temperature. Primary antibodies mouse anti-TH antibody (MAB318, Millipore), mouse anti-GAD67 antibody (MAB5406, Millipore) were used to identify the cell type. They were co-stained with rabbit anti-GPR37 (Cat#64407, Cell Signaling Technology) and Alexa 488 conjugated streptavidin overnight at 4°C. Afterward, cells were incubated with fluorochrome-conjugated secondary antibodies for 2 hours at room temperature and mounted in mounting media. Z-stacked images were obtained with a Zeiss LSM 880 Airyscan confocal microscope (Zeiss, Germany).

### Cell counting

For counting cell in the VTA, 40 μm coronal sections from bregma -2.9 to -3.8 mm for each mouse were collected for immunohistochemistry. The brain sections were mounted and imaging using a Zeiss LSM 880 Airyscan confocal microscope and Olympus VS120 virtual microscopy slide scanning system. Then the immunostaining was analyzed and counted with image J, image Pro- plus and Photoshop software.

### Behavioral analysis

Behavioral analysis of the looming test was performed as described previously (Tseng et al, 2022; Zhou et al, 2019). Behavioral data were recorded and analyzed with Adobe Premiere. The following parameters were calculated to define the looming evoked defensive behavior: (ⅰ) flight latency: time from the looming presentation to onset the flight to the nest, (ⅱ) return time: time from the onset of looming until returning to the nest, (ⅲ) time in the nest: time spent in the nest following escape. Data obtained from mice with imprecise cannula placement or virus expression were not used for analyses.

## QUANTIFICATION AND STATISTICAL ANALYSIS

All statistics were performed in Graph Pad Prism (GraphPad Software, Inc.). Paired student test, unpaired student test, one-way ANOVA and two-way ANOVA were used where appropriate. In all statistical measures a P value <0.05 was considered statistically significant. Post hoc significance values were set as *P< 0.05, **P< 0.01, ***P< 0.001 and ****P< 0.0001; all statistical tests used are indicated in the figure legends.

---

**Figure S1.**
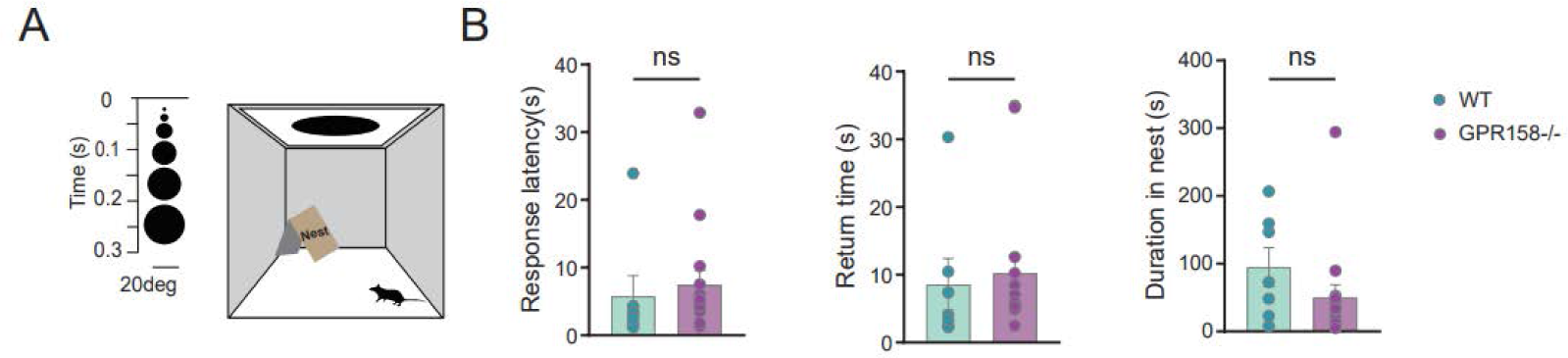
*GPR158* mutant mice do not exhibit significant changes of visual escape responses, related to Figure 1. **(A)** Schematic paradigm and timeline. **(B)** Bar graph showing WT mice and *GPR158*-/- mice respond to looming: 1) response latency (Two-tailed unpaired t test, p>0.05), 2) return time (Two-tailed unpaired t test, p>0.05) and 3) duration in the nest (Two-tailed unpaired t test, p>0.05) responding to LS (WT mice, n=7; *GPR158*-/- mice, n=15).

**Figure S2.**
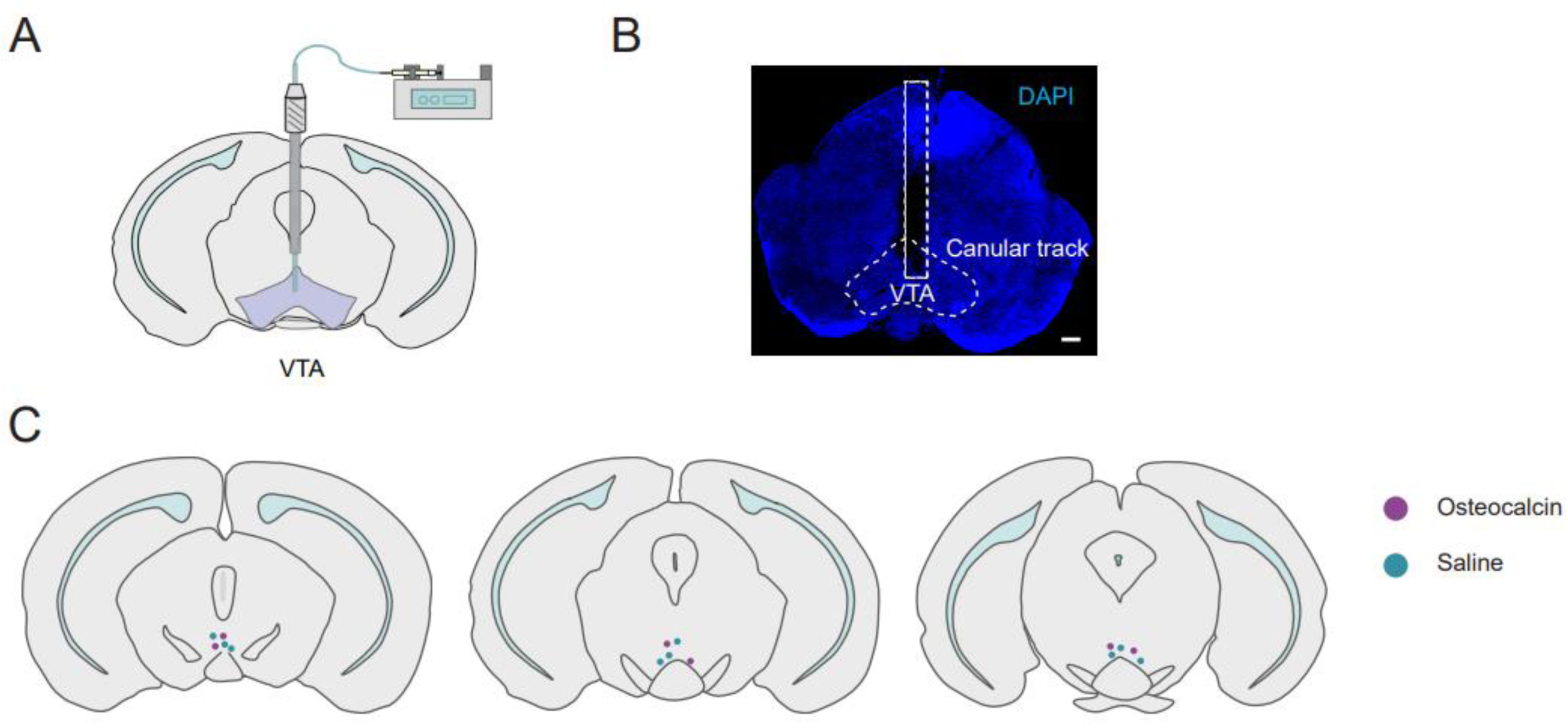
Validation of canular track for OCN administration in VTA, related to Figure 2. **(A)** Schematic showing OCN pumping into VTA. **(B)** Canular track location in VTA. **(C)** Schematics showing the canular location in VTA in osteocalcin and saline injection group.

**Figure S3.**
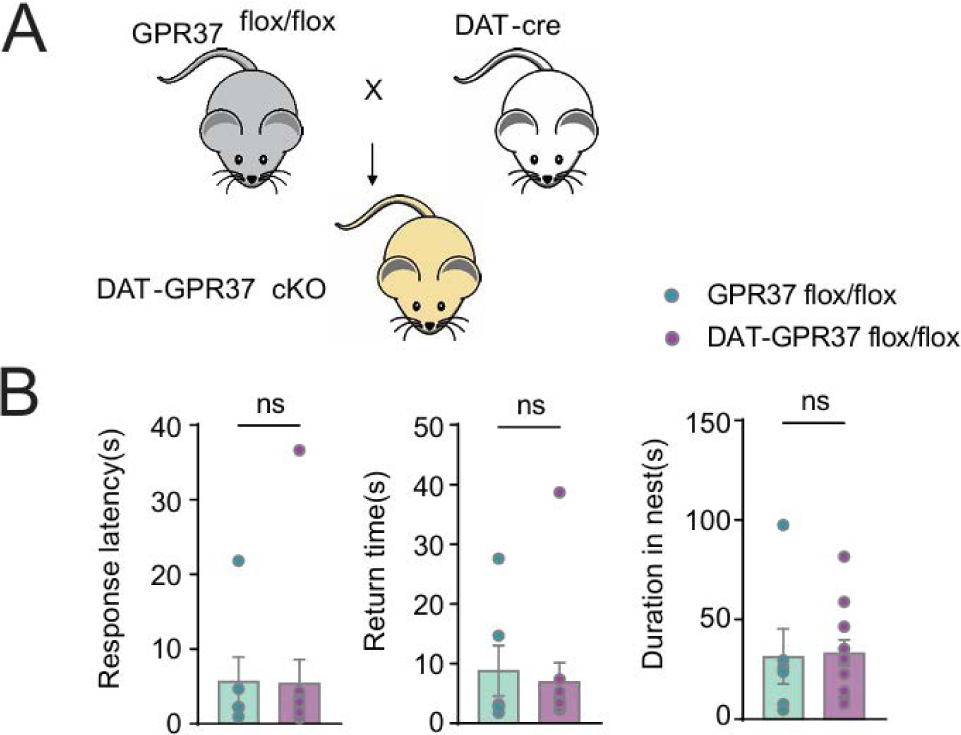
GPR37 receptor ablation in DA neurons do not exhibit significant changes of visual escape responses, related to Figure 3. **(A)** Schematic showing construction of mice with GPR37 deletion in DA neurons. **(B)** Bar graph showing the looming response in DAT-ires-Cre; GPR37^LoxP/Lox^P mice (n=11) and control mice (n=6): 1) response latency (Two-tailed unpaired t test, p= 0.9653), 2) return time (Two-tailed unpaired t test, p= 0.7302) and 3) duration in the nest (Two-tailed unpaired t test, p= 0.8989).

**Figure S4.**
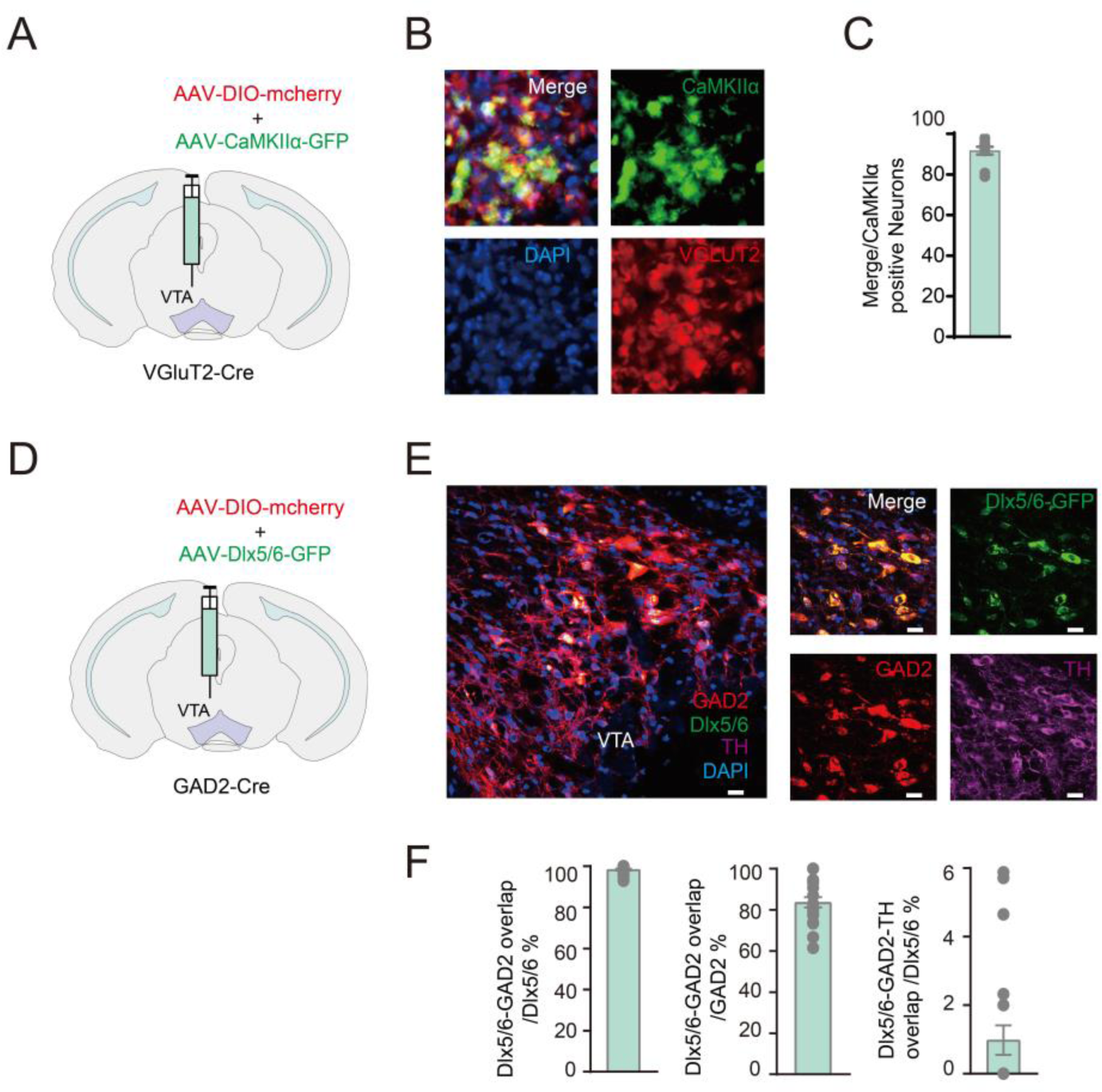
Validation of effectiveness and specificity in labeling VTA Glut or GABA neurons by using AAVs expressing neuronal type specific Cre, related to Figure 3. **(A)** Schematic of viral injection in VGluT2-ires-Cre mice. (**B-C**) Overlap between mcherry and GFP expression and immunostaining of TH in VTA (1433 cells from 3 mice). **(D)** Schematic of viral injection in the GAD2-ires-Cre mice. (**E-F**) Overlap between mcherry and GFP expression and immunostaining of GAD2 in VTA (1236 cells from 3 mice).

**Figure S5.**
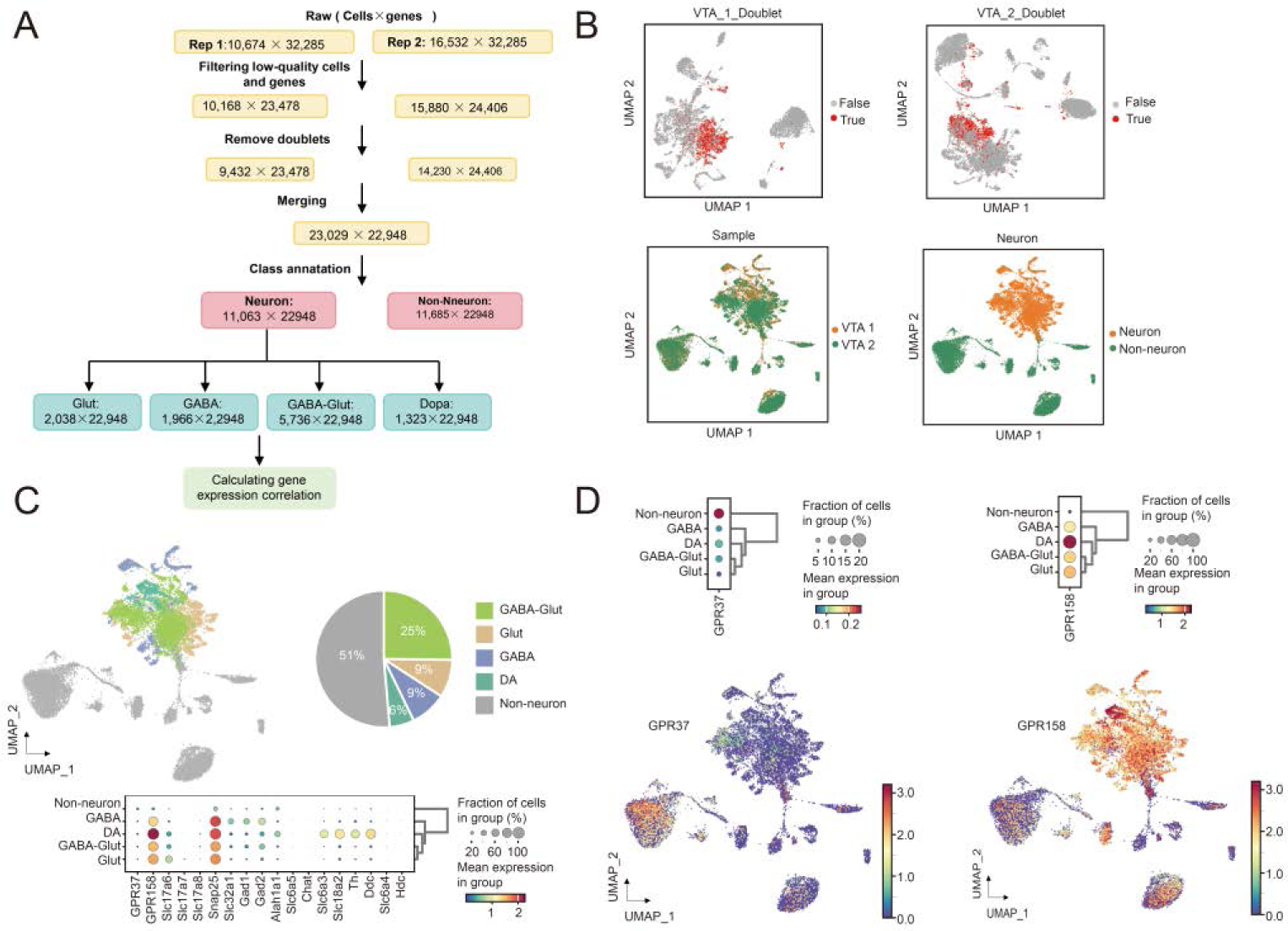
VTA snRNA-seq data analysis, related to Figure 4. **(A)** Analysis pipeline. **(B)** Analysis in two biological replicates to filterout doublets and low quality cells, and merging of two replicates to identify neuron and non-neuronal cells. **(C)** Five VTA cell types annotated based on canonical marker genes and their proportions, with dot plots showing expression of these marker genes and two GPRs. **(D)** GPR37 and GPR158 expression in all VTA cells shown in UMAP.

**Figure S6.**
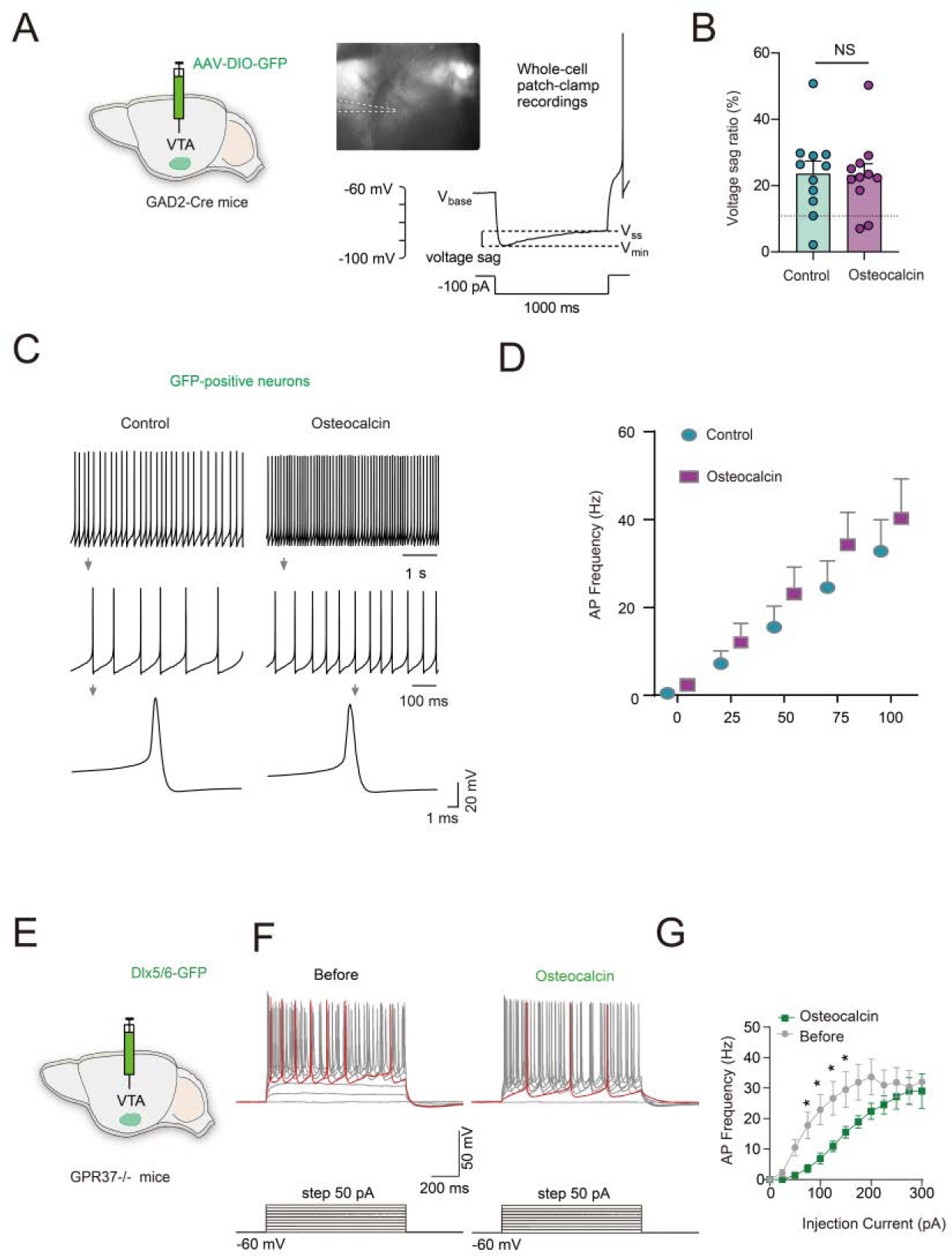
VTA GABAergic neurons exhibit opposite electrophysiological responses to osteocalcin contingent on the expression of GPR37 receptors, related to Figure 5. **(A)** Schematic showing recording of VTA GAD2 positive neurons using patch-clamp. **(B)** Voltage sag ratio of type I VTA GAD2 positive neurons before and after osteocalcin infusion (n=11 cells). (**C-D**) Type I VTA GAD2 positive neurons in responses to a series of current injections. **(E)** Labeling of GABAergic neurons in *GPR37*-/- mice. (**F-G**) Firing rate of VTA GABAergic neurons in *GPR37*-/- mice was solely decreased by OCN (n=9 cells).

**Figure S7.**
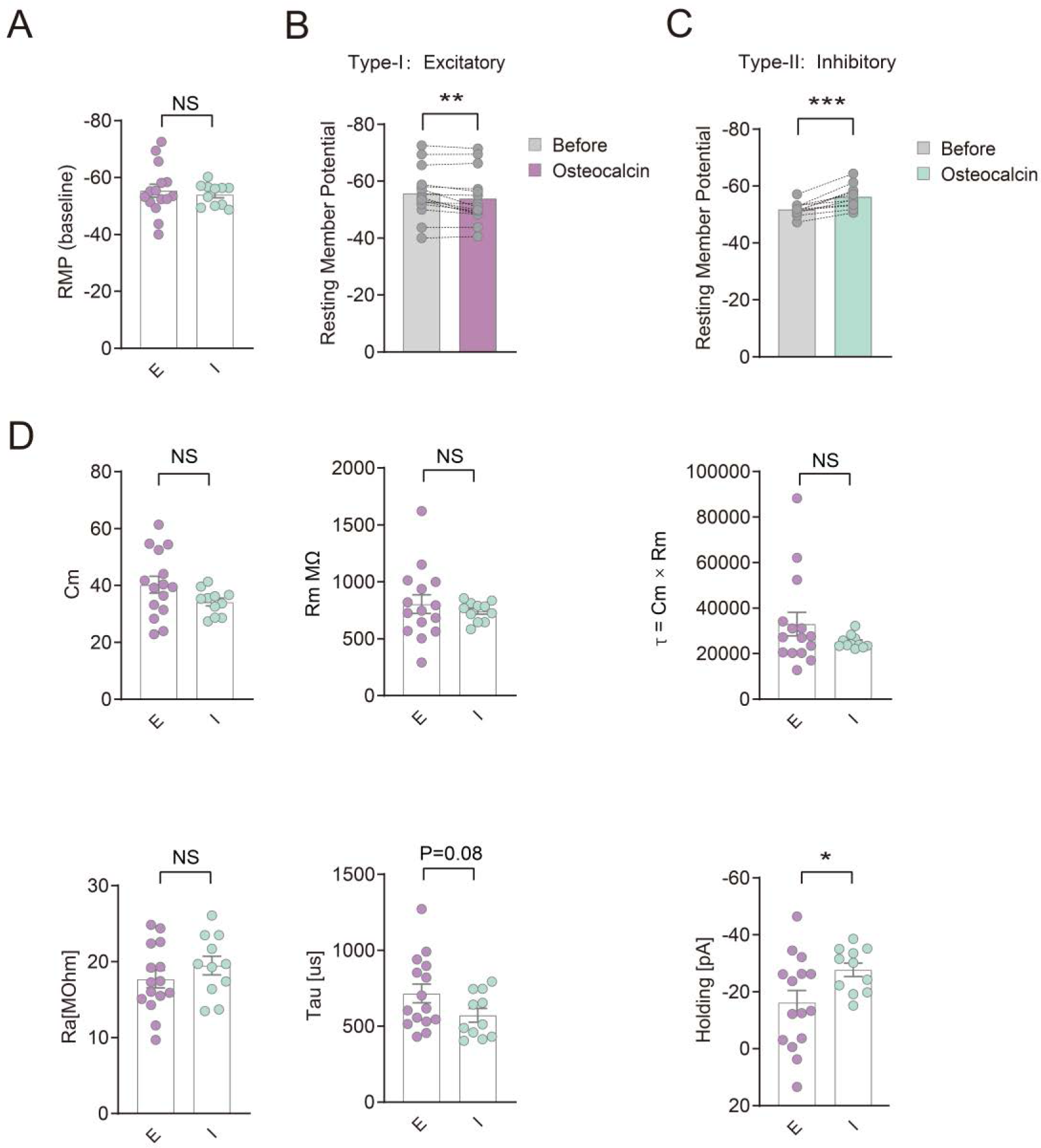
The basic electrophysiological parameters of the cells that are excited and inhibited by OCN, related to Figure 5. **(A)** RMP of the cells are excited (n=15) and inhibited (n=11) by OCN. **(B)** RMP of the cell that are excited before and after OCN treatment. **(C)** RMP of the cell that are inhibited before and after OCN treatment. **(D)** Basic electrophysiological parameters of the cells that are excited and inhibited by OCN (Cm, Rm, τ, Ra, Tau and Holding).

**Figure S8.**
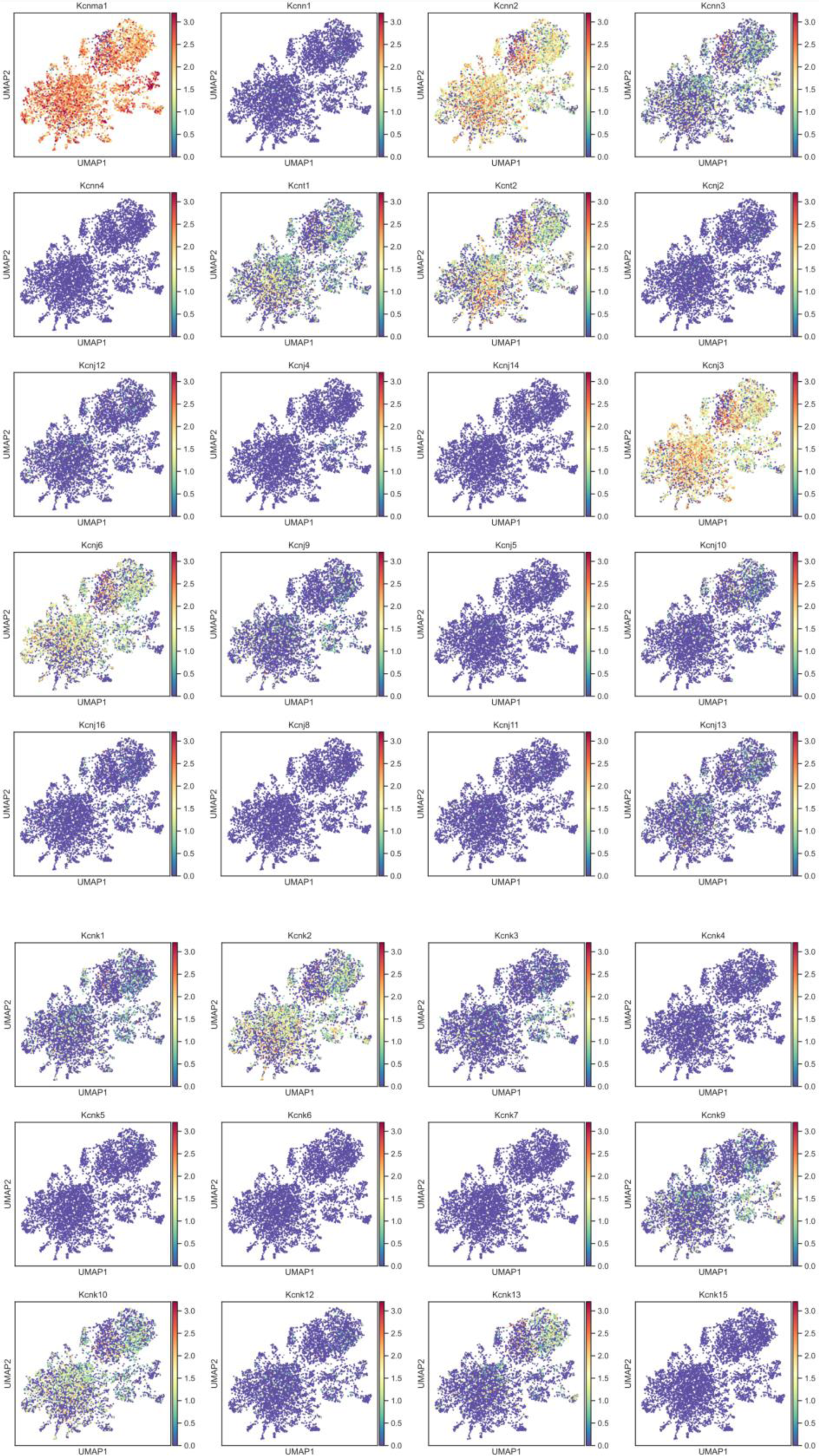
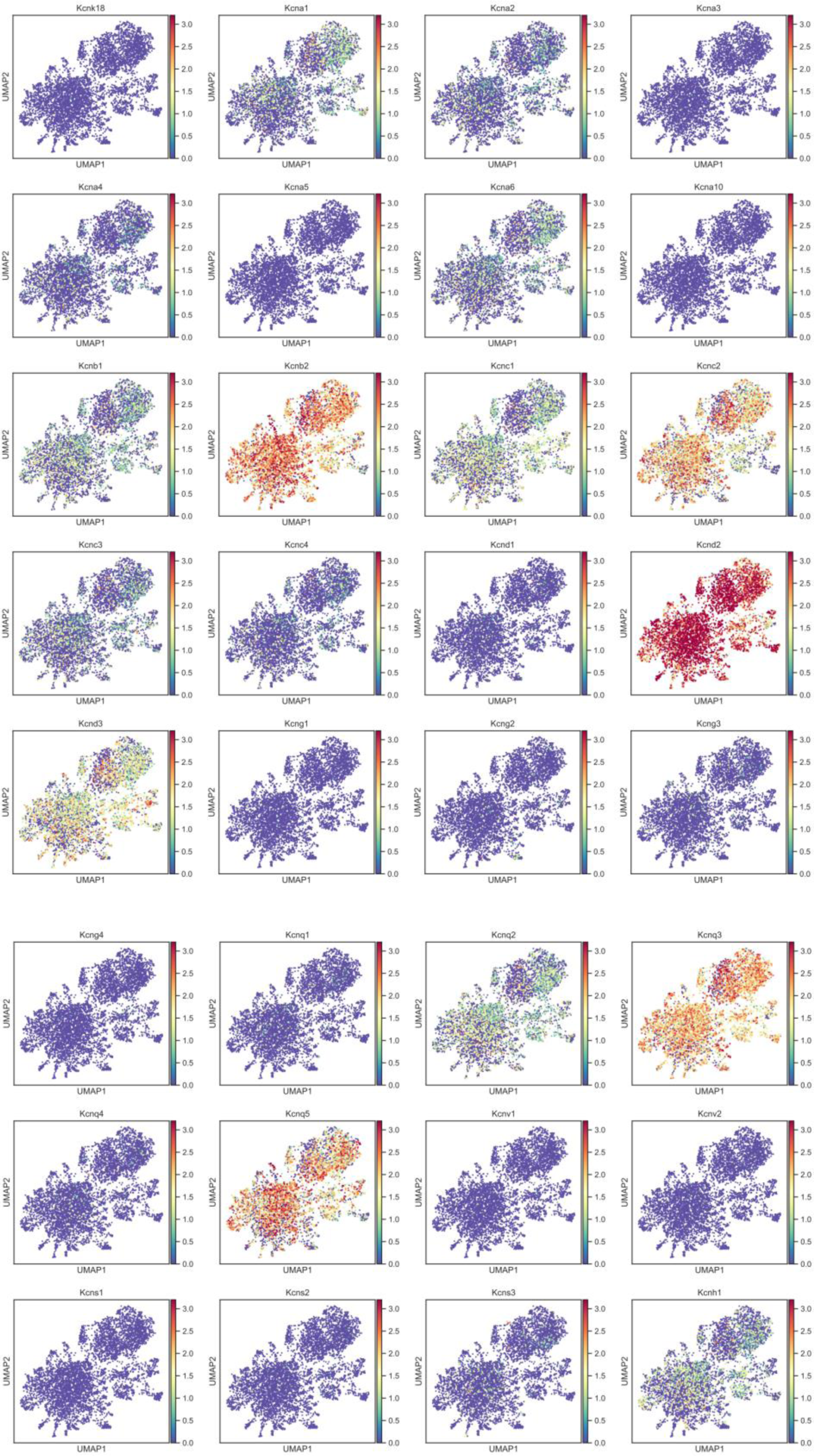
Expression of each potassium channel genes across the three VTA GABA-Glut subtypes shown in UMAP plot, related to Figure 6.

**Figure S9.**
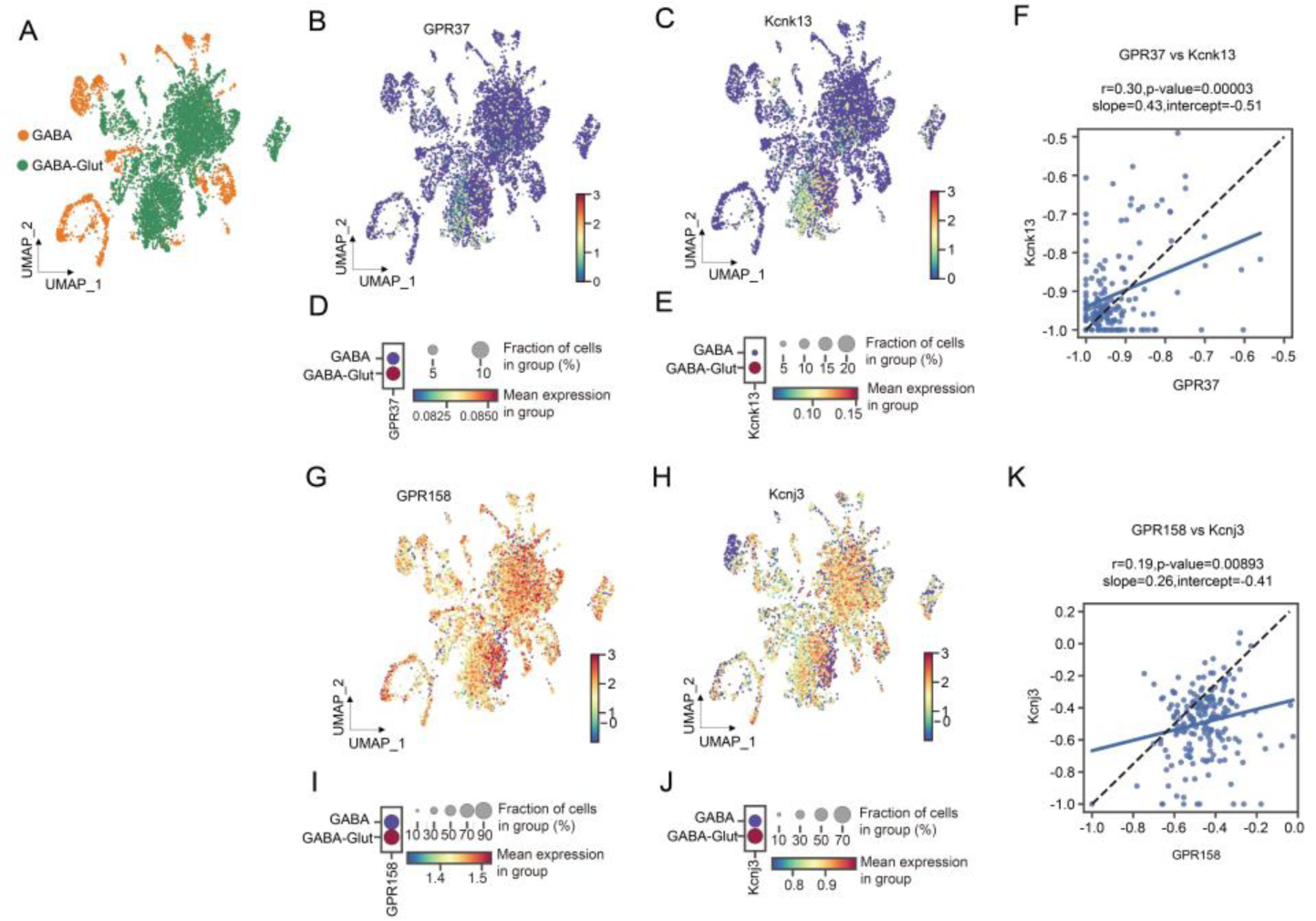
Expression and correlation of expression of genes encoding the two GPRs and potassium channels in all VTA GABAergic neurons, related to Figure 6. **(A)** UMAP of VTA GABAergic neurons, including GABA and GABA-Glut shown in Figure 4. **(B-E)** Expression of *GPR37* and *Kcnk13* in GABAergic neurons. **(F)** Pearson correlation of *GPR37* and *Kcnk13* expression, with p-value, slope and intercept of linear regression. **(G-J)** Expression of *GPR158* and *Kcnj3* in GABAergic neurons. **(K)** As in F, but for *GPR158* and *Kcnj3*.

**Figure S10.**
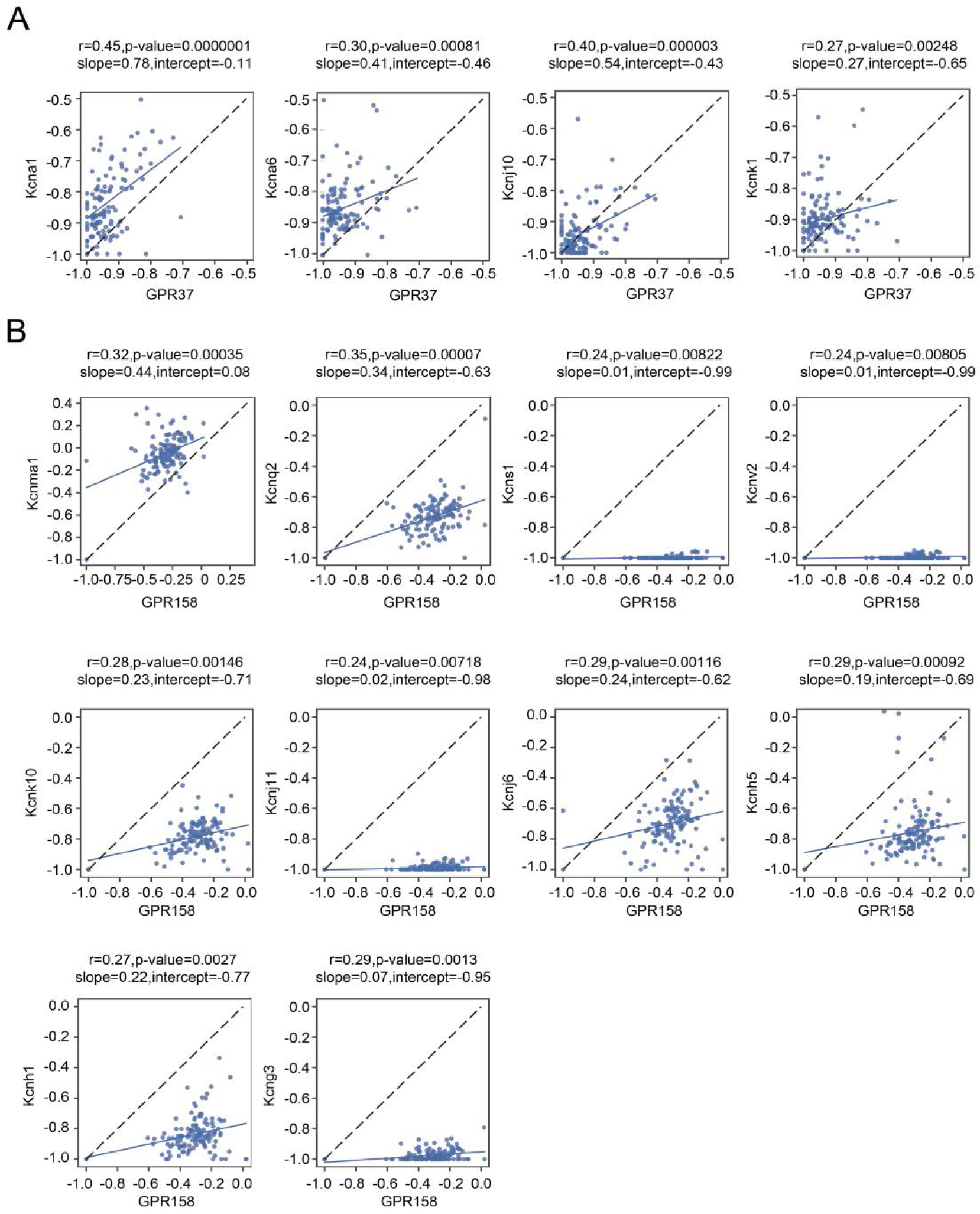
Pearson correlation of expression in the UMAP of selected potassium channels with either GPR37 or GPR158, related to Figure 6. **(A)** Pearson correlation of *GPR37* and four potassium channel gene expression, with p-value, slope and intercept of linear regression. Channels shown had p<0.01. **(B)** As in A, but for *GPR158*.

**Figure S11.**
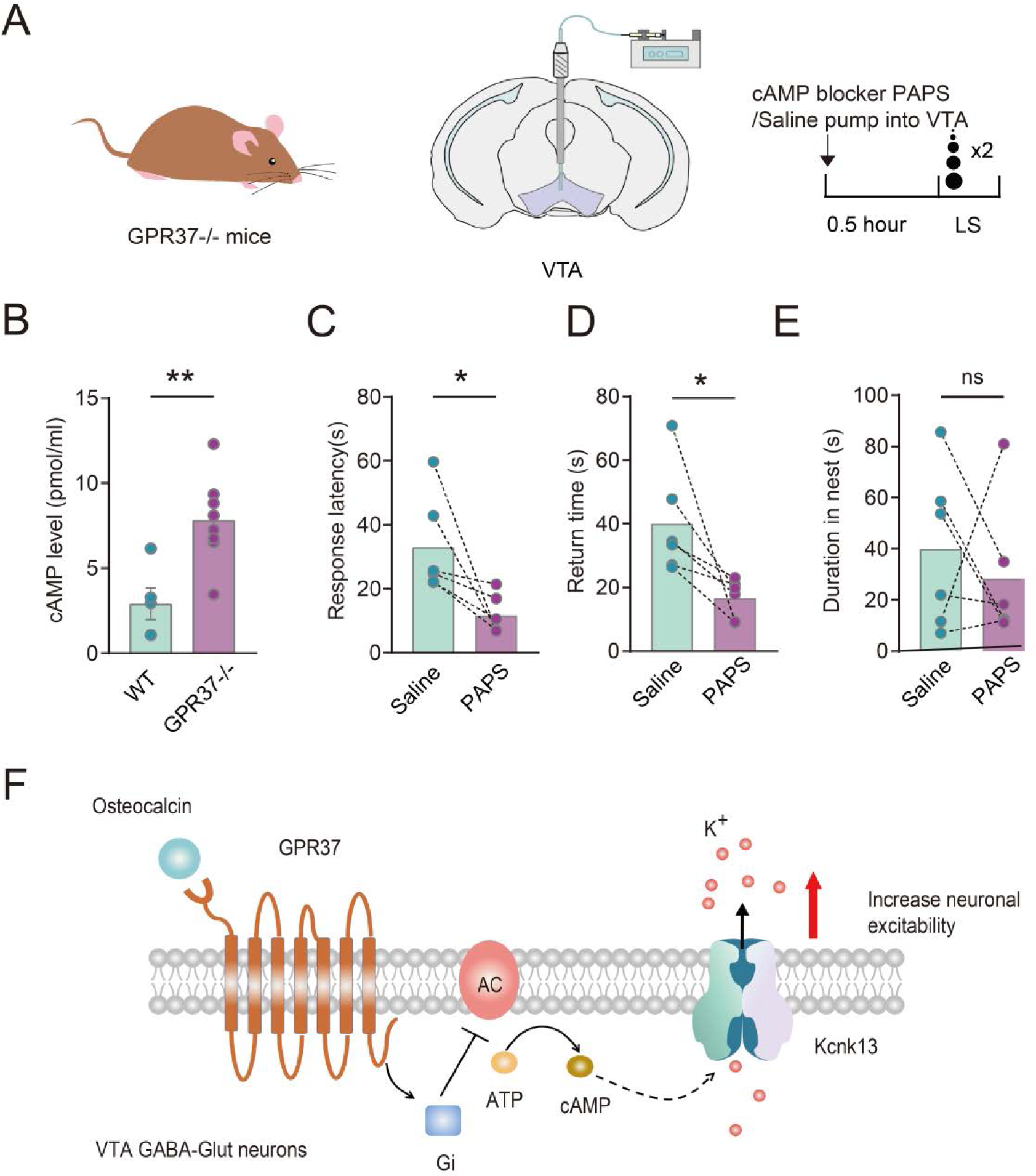
GPR37 is required for cAMP level reduction in the VTA that is essential for rapid visual escape. **(A)** Schematics showing cAMP blocker PAPS (0.3mM) pumped into VTA of *GPR37*-/- mice. **(B)** cAMP level in WT and *GPR37*-/- mice. **(C-E)** Bar graph showing changes on looming response after infusion of the cAMP blocker in *GPR37*-/- mice (n=6). Left: response latency (Two-tailed paired t test, *p=0.0414); middle: return time (Two-tailed paired t test, *p=0.0364); right: duration in the nest (Two-tailed paired t test, p=0.5684) . **(F)** Proposed model of OCN-GPR37 signaling in VTA GABAergic neuron subpopulation.

