## Supplementary figures and images for "A bone-derived protein primes rapid visual escape via GPR37 receptor in a subpopulation of VTA GABAergic neurons"

### S-Fig 1

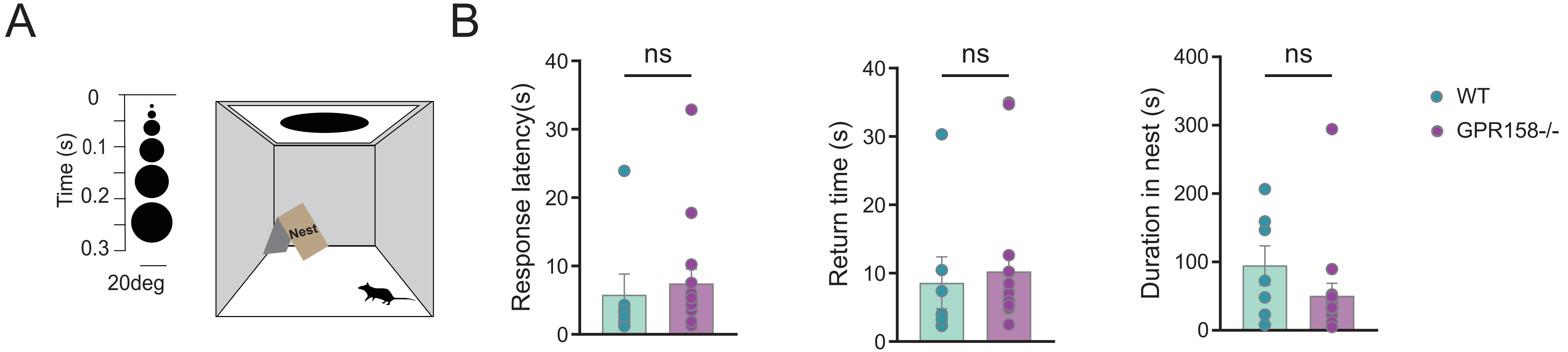

### S-Fig 2

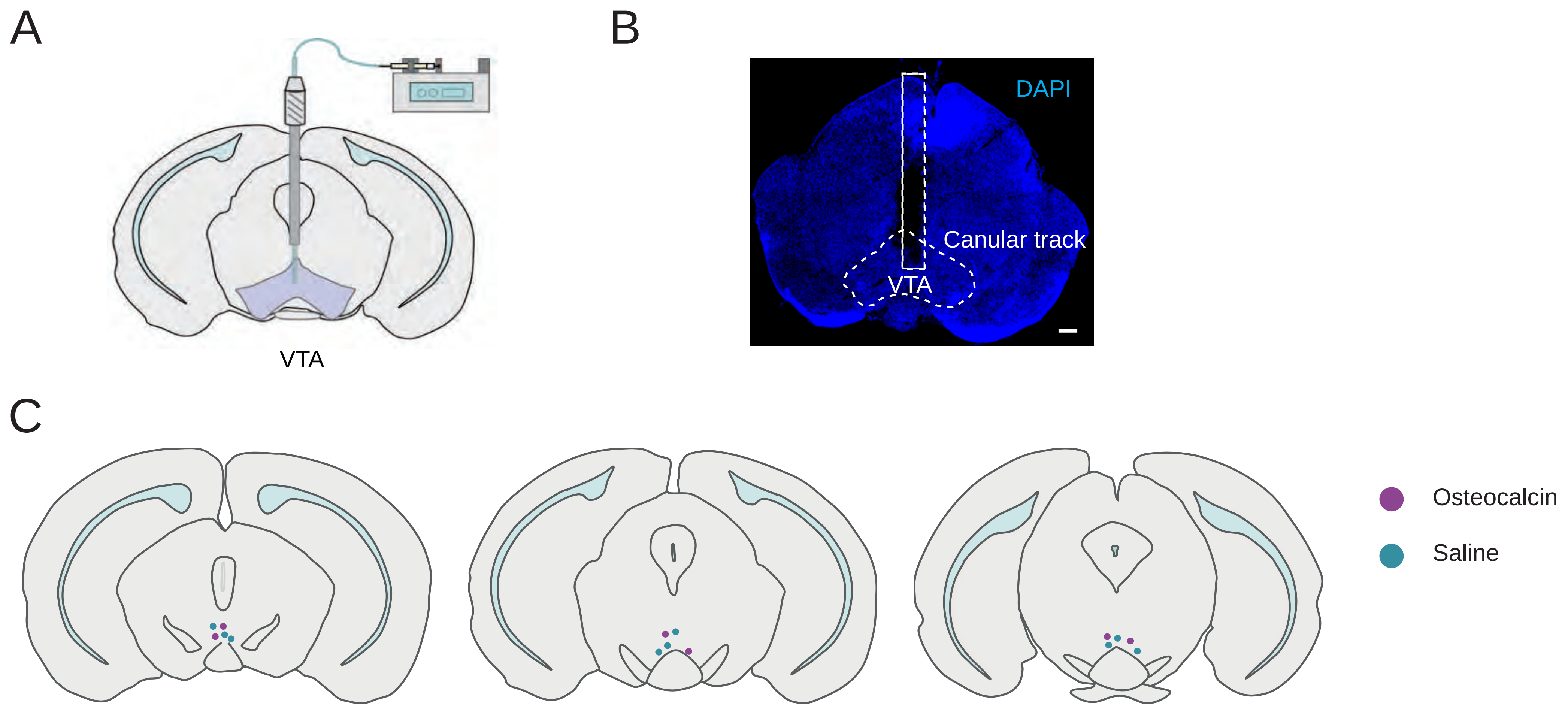

### S-Fig 3

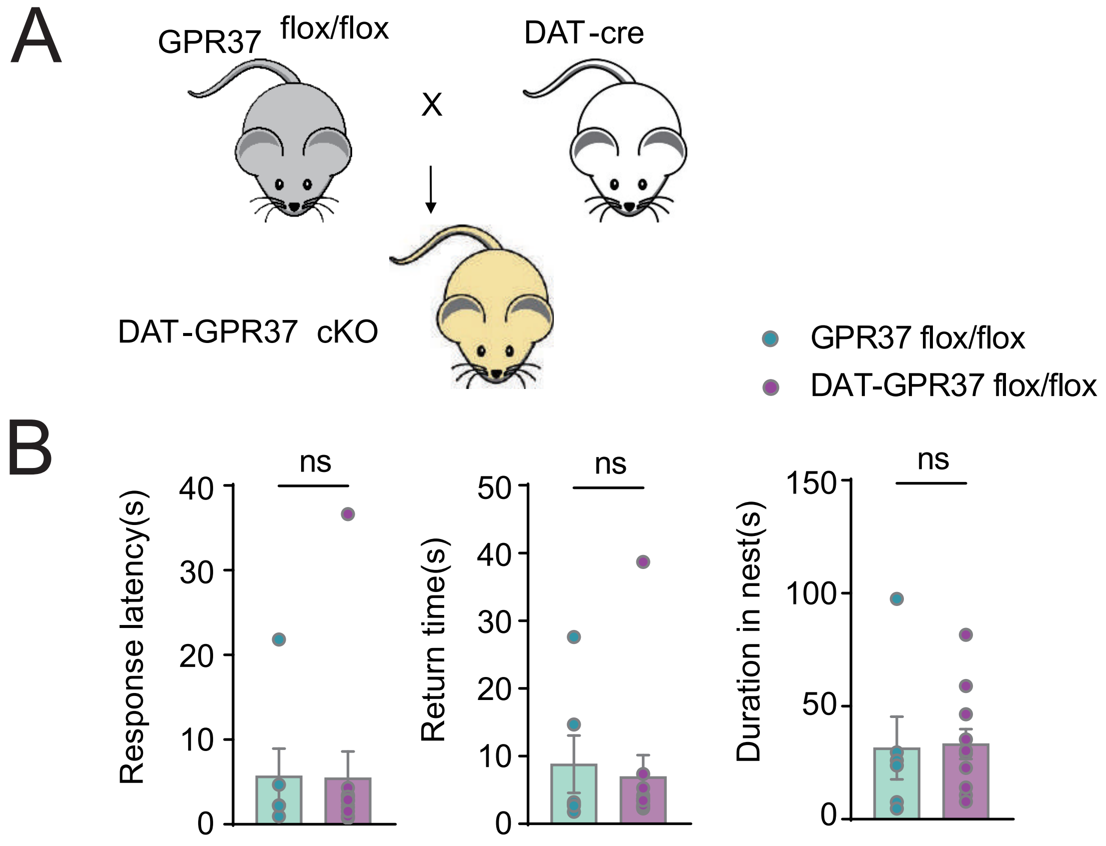

### S-Fig 4

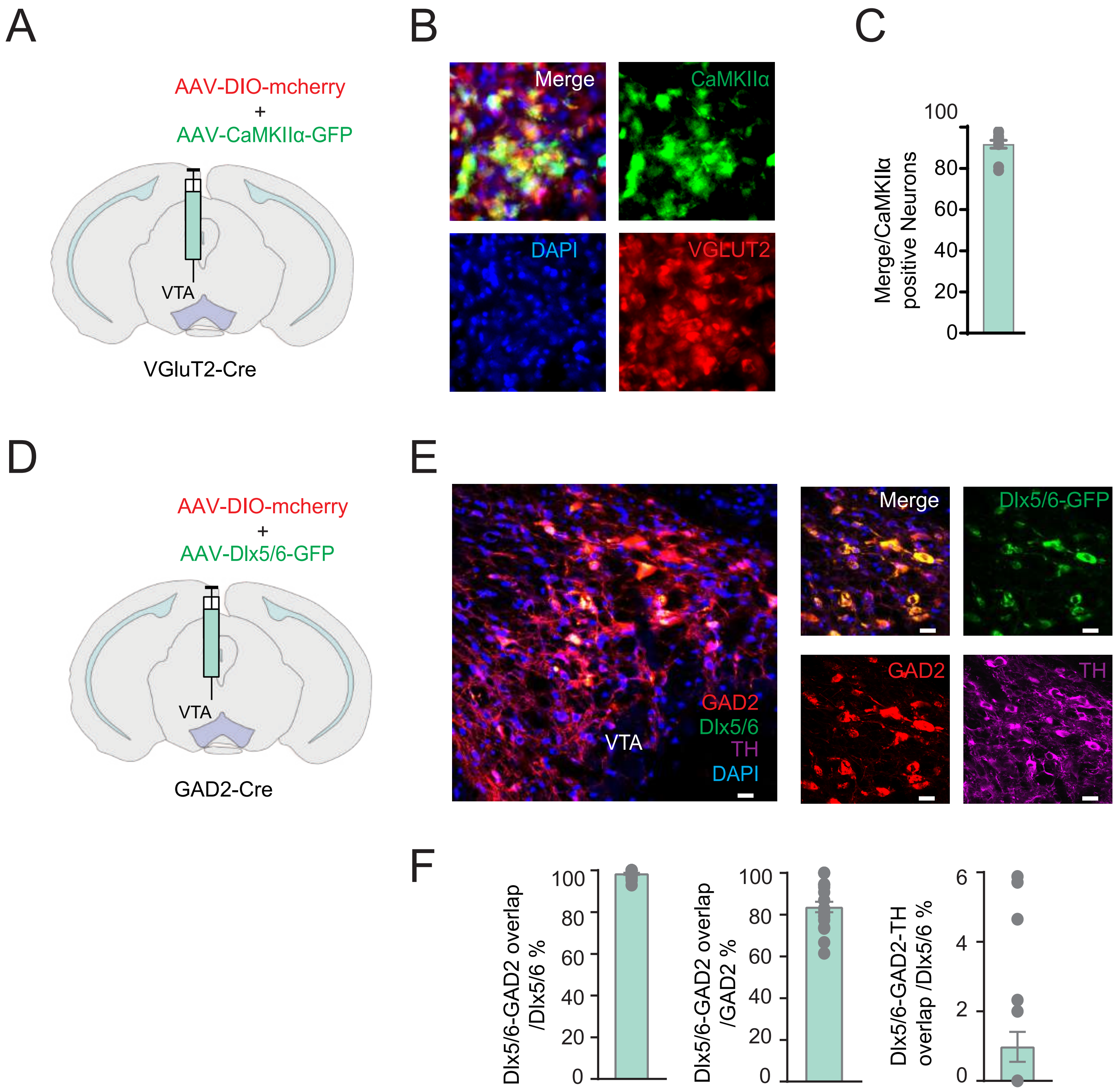

### S-Fig 5

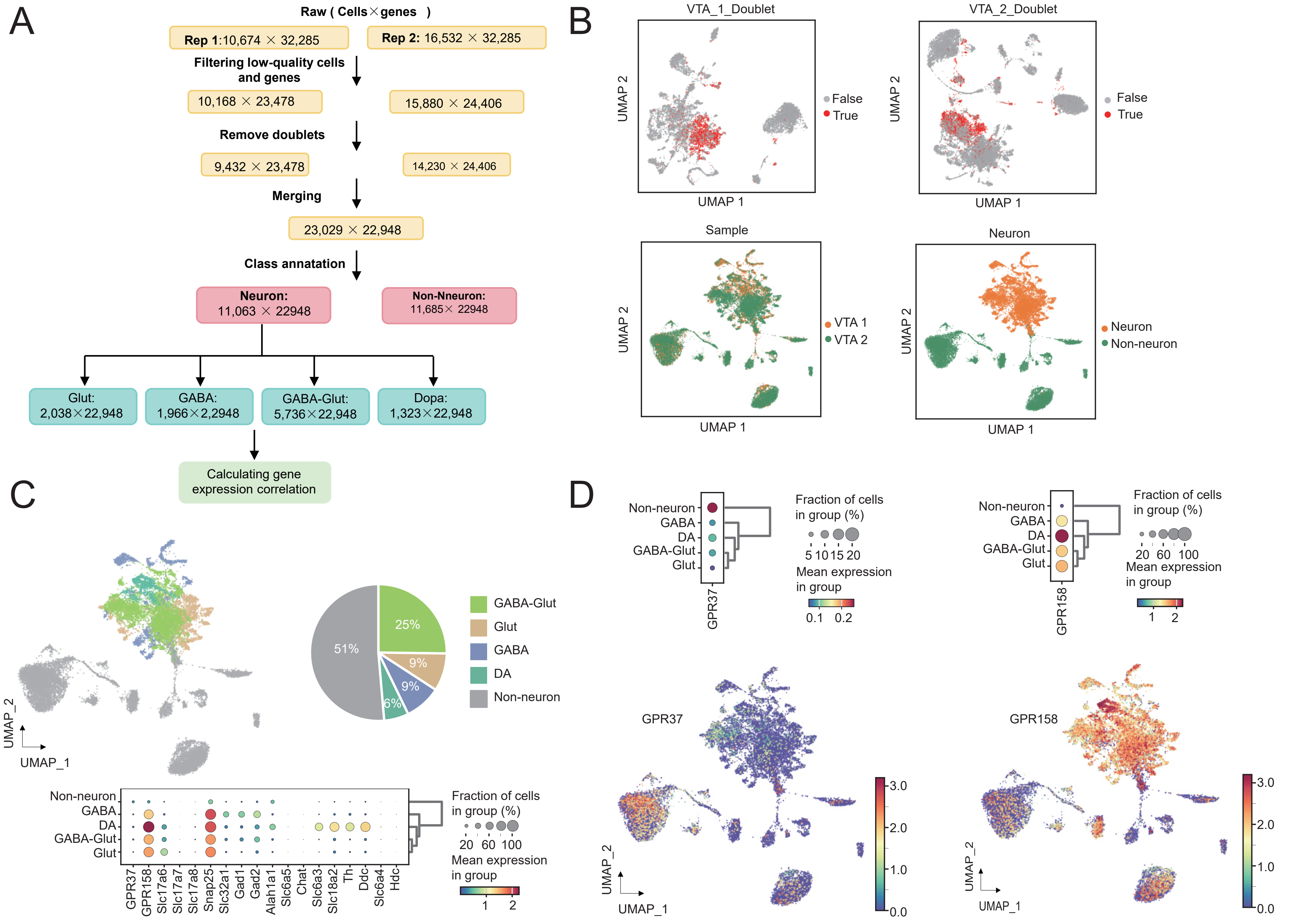

### S-Fig 6

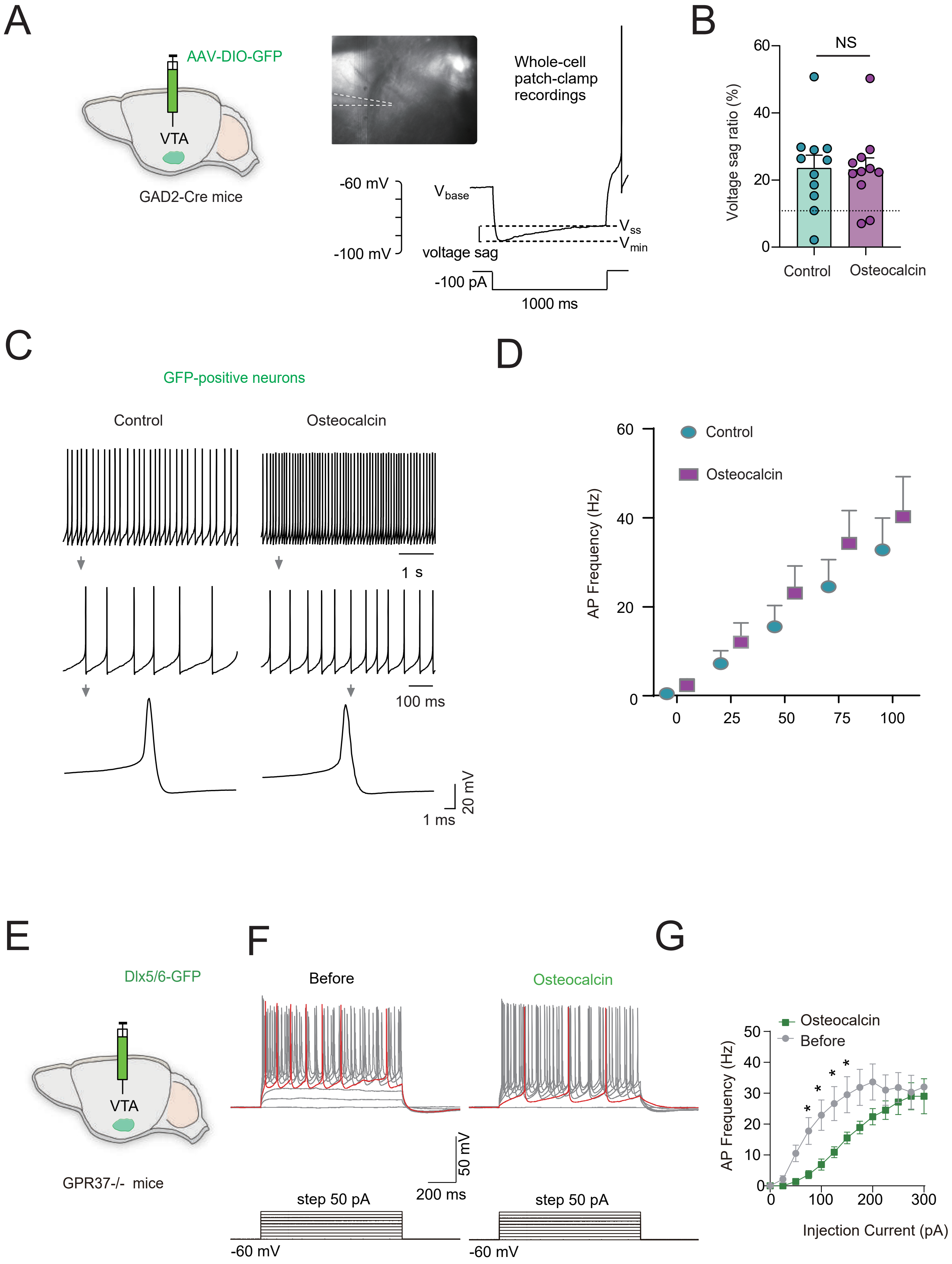

### S-Fig 7

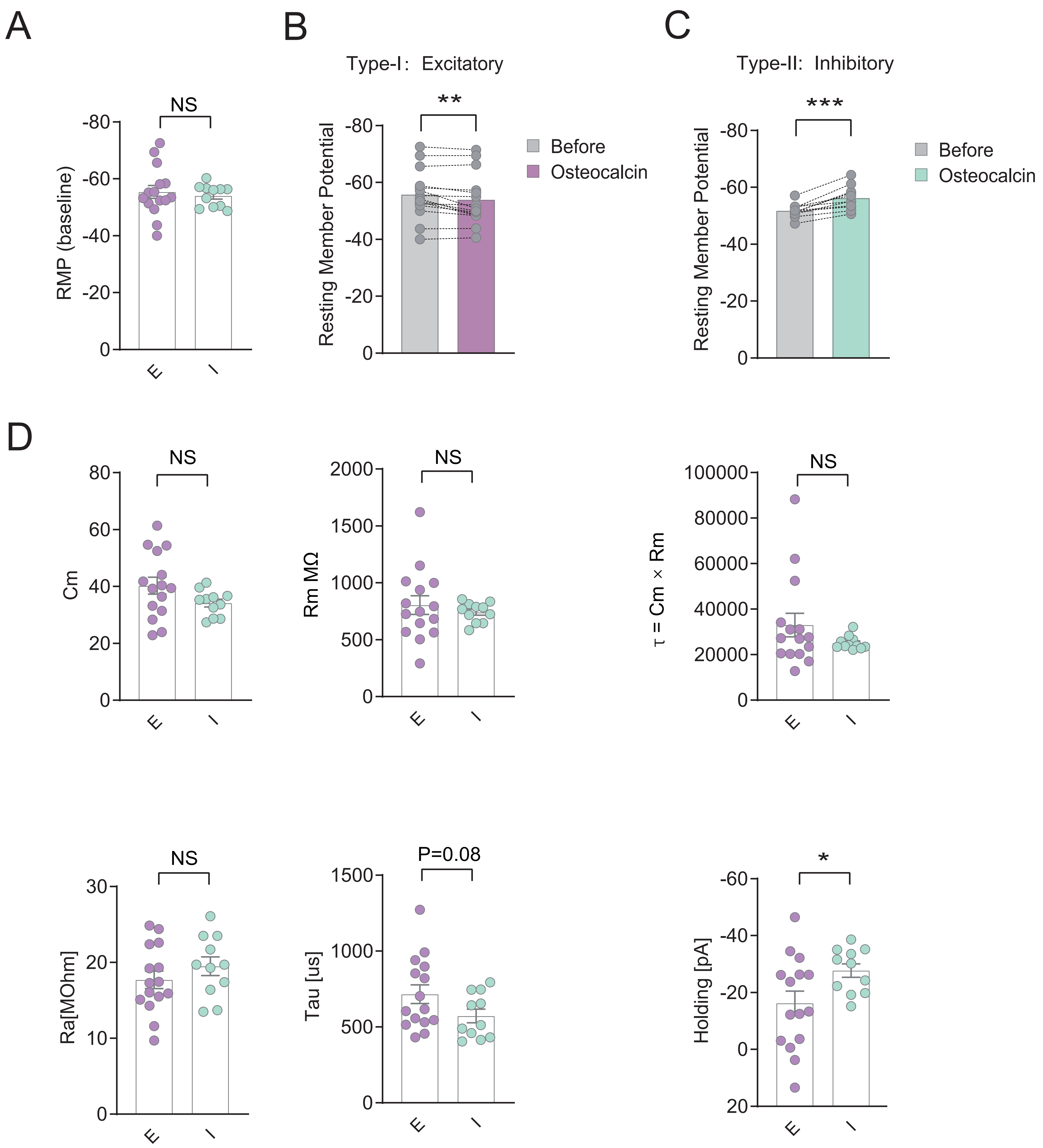

### S-Fig 8

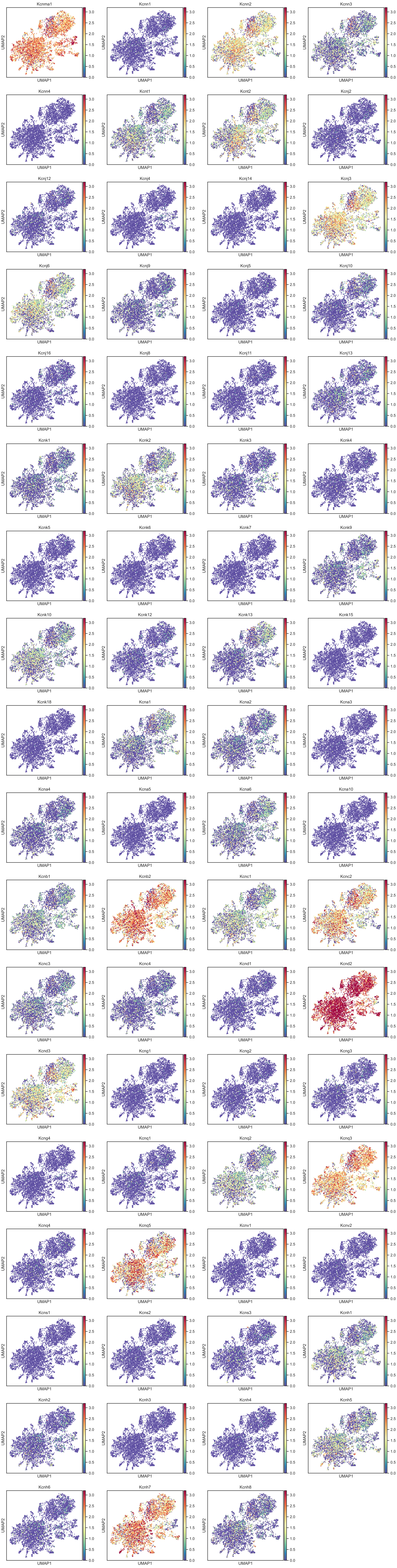

### S-Fig 9

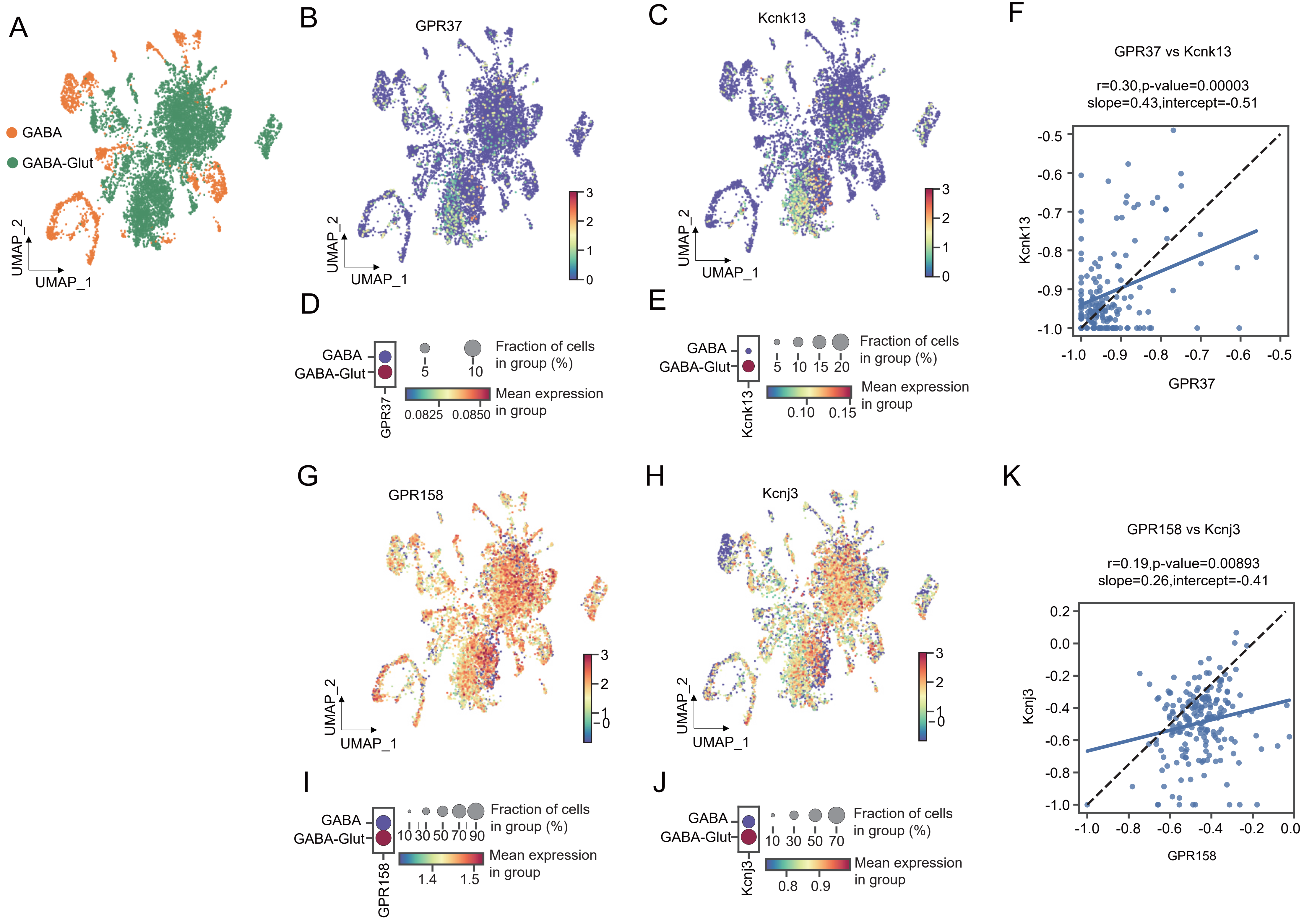

### S-Fig 10

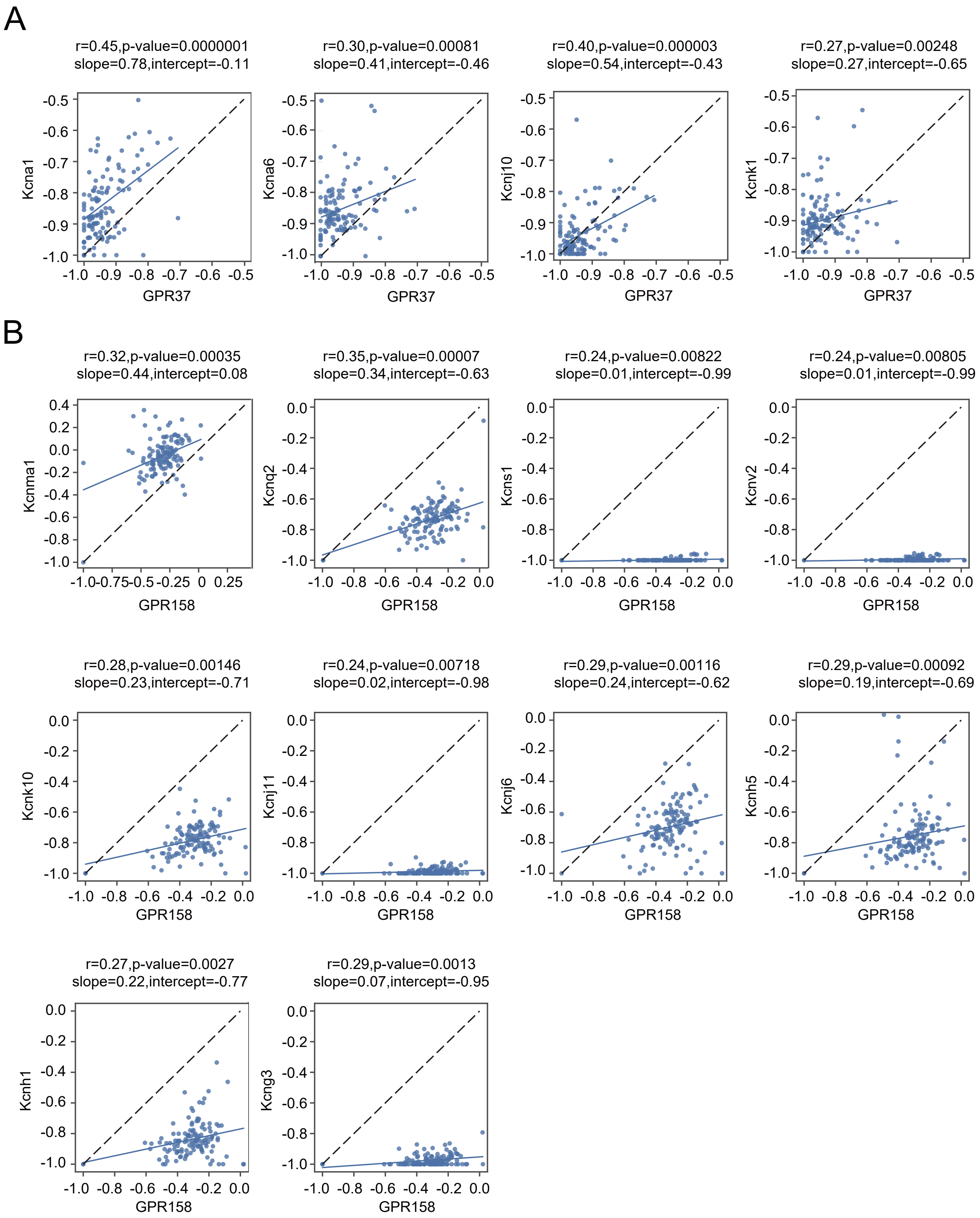

### S-Fig 11

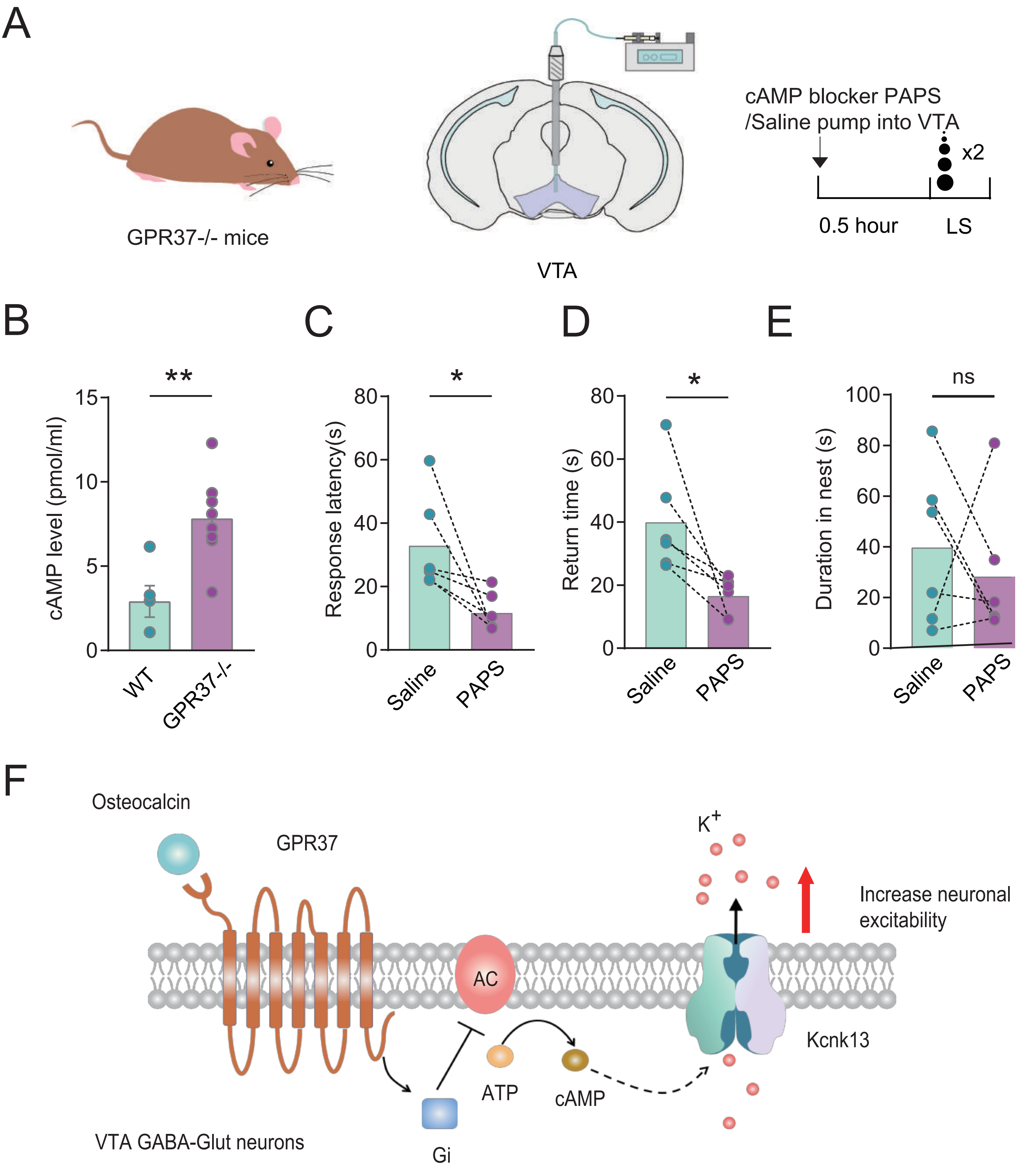
