## Supplementary material for "A bone-derived protein primes rapid visual escape via GPR37 receptor in a subpopulation of VTA GABAergic neurons": Table 1

| | | | | |
|---|---|---|---|---|
| Gpr37_Kcnma1 | r = -0.02 | p-value= 0.86292 | slope = -0.01 | intercept= -0.06 |
| Gpr37_Kcnn1 | r = 0.05 | p-value= 0.57950 | slope = 0.01 | intercept= -0.98 |
| Gpr37_Kcnn2 | r = -0.09 | p-value= 0.34581 | slope = -0.14 | intercept= -0.56 |
| Gpr37_Kcnn3 | r = -0.11 | p-value= 0.22790 | slope = -0.33 | intercept= -1.13 |
| Gpr37_Kcnn4 | r = 0.09 | p-value= 0.31350 | slope = 0.00 | intercept= -1.00 |
| Gpr37_Kcnt1 | r = 0.05 | p-value= 0.58919 | slope = -0.02 | intercept= -0.83 |
| Gpr37_Kcnt2 | r = 0.02 | p-value= 0.78620 | slope = -0.19 | intercept= -0.83 |
| Gpr37_Kcnj2 | r = 0.16 | p-value= 0.07004 | slope = 0.09 | intercept= -0.90 |
| Gpr37_Kcnj12 | r = 0.03 | p-value= 0.71825 | slope = -0.09 | intercept= -1.05 |
| Gpr37_Kcnj4 | r = 0.04 | p-value= 0.63577 | slope = -0.01 | intercept= -1.00 |
| Gpr37_Kcnj14 | r = 0.03 | p-value= 0.77055 | slope = -0.03 | intercept= -1.02 |
| Gpr37_Kcnj3 | r = -0.20 | p-value= 0.02529 | slope = -0.39 | intercept= -0.79 |
| Gpr37_Kcnj6 | r = -0.17 | p-value= 0.06183 | slope = -0.36 | intercept= -1.04 |
| Gpr37_Kcnj9 | r = -0.04 | p-value= 0.66513 | slope = -0.10 | intercept= -1.02 |
| Gpr37_Kcnj5 | r = 0.02 | p-value= 0.82485 | slope = -0.01 | intercept= -1.00 |
| **Gpr37_Kcnj10** | **r = 0.40** | **p-value= 0.000003** | **slope = 0.54** | **intercept= -0.43** |
| Gpr37_Kcnj16 | r = 0.09 | p-value= 0.34532 | slope = -0.15 | intercept= -1.11 |
| Gpr37_Kcnj8 | r = 0.04 | p-value= 0.63776 | slope = -0.00 | intercept= -1.00 |
| Gpr37_Kcnj11 | r = 0.14 | p-value= 0.11180 | slope = 0.01 | intercept= -0.98 |
| Gpr37_Kcnj13 | r = -0.05 | p-value= 0.60845 | slope = -0.09 | intercept= -1.02 |
| **Gpr37_Kcnk1** | **r = 0.27** | **p-value= 0.00248** | **slope = 0.27** | **intercept= -0.65** |
| Gpr37_Kcnk2 | r = 0.07 | p-value= 0.41368 | slope = 0.05 | intercept= -0.70 |
| Gpr37_Kcnk3 | r = -0.01 | p-value= 0.87813 | slope = -0.08 | intercept= -1.00 |
| Gpr37_Kcnk4 | r = 0.02 | p-value= 0.79183 | slope = -0.02 | intercept= -1.01 |
| Gpr37_Kcnk5 | r = 0.14 | p-value= 0.13160 | slope = 0.01 | intercept= -0.99 |
| Gpr37_Kcnk6 | r = 0.08 | p-value= 0.37389 | slope = -0.00 | intercept= -1.00 |
| Gpr37_Kcnk7 | r = -0.04 | p-value= 0.68873 | slope = -0.00 | intercept= -1.00 |
| Gpr37_Kcnk9 | r = 0.21 | p-value= 0.01920 | slope = 0.22 | intercept= -0.68 |
| Gpr37_Kcnk10 | r = -0.09 | p-value= 0.30954 | slope = -0.18 | intercept= -0.95 |
| Gpr37_Kcnk12 | r = 0.10 | p-value= 0.24958 | slope = 0.05 | intercept= -0.92 |
| **Gpr37_Kcnk13** | **r = 0.32** | **p-value= 0.00024** | **slope = 0.97** | **intercept= 0.02** |
| Gpr37_Kcnk15 | r = 0.14 | p-value= 0.11902 | slope = 0.02 | intercept= -0.98 |
| Gpr37_Kcnk18 | r = -0.12 | p-value= 0.16966 | slope = -0.01 | intercept= -1.01 |
| **Gpr37_Kcna1** | **r = 0.45** | **p-value= 0.0000001** | **slope = 0.78** | **intercept= -0.11** |
| Gpr37_Kcna2 | r = 0.15 | p-value= 0.09171 | slope = 0.24 | intercept= -0.65 |
| Gpr37_Kcna3 | r = -0.05 | p-value= 0.61459 | slope = -0.03 | intercept= -1.02 |
| Gpr37_Kcna4 | r = 0.18 | p-value= 0.04423 | slope = 0.15 | intercept= -0.79 |
| Gpr37_Kcna5 | r = 0.02 | p-value= 0.82768 | slope = -0.00 | intercept= -1.00 |
| **Gpr37_Kcna6** | **r = 0.30** | **p-value= 0.00081** | **slope = 0.41** | **intercept= -0.46** |
| Gpr37_Kcna10 | r = -0.10 | p-value= 0.27320 | slope = -0.02 | intercept= -1.01 |
| Gpr37_Kcnb1 | r = 0.10 | p-value= 0.28045 | slope = 0.16 | intercept= -0.65 |
| Gpr37_Kcnb2 | r = -0.11 | p-value= 0.20515 | slope = -0.45 | intercept= -0.61 |
| Gpr37_Kcnc1 | r = 0.06 | p-value= 0.53361 | slope = -0.17 | intercept= -0.81 |
| Gpr37_Kcnc2 | r = 0.03 | p-value= 0.70514 | slope = -0.11 | intercept= -0.59 |
| Gpr37_Kcnc3 | r = 0.03 | p-value= 0.70670 | slope = -0.06 | intercept= -0.92 |
| Gpr37_Kcnc4 | r = 0.20 | p-value= 0.02328 | slope = 0.02 | intercept= -0.90 |
| Gpr37_Kcnd1 | r = -0.15 | p-value= 0.10794 | slope = -0.10 | intercept= -1.08 |
| Gpr37_Kcnd2 | r = -0.14 | p-value= 0.11314 | slope = -0.47 | intercept= -0.43 |
| Gpr37_Kcnd3 | r = 0.01 | p-value= 0.88916 | slope = -0.05 | intercept= -0.59 |
| Gpr37_Kcng1 | r = -0.07 | p-value= 0.46034 | slope = -0.03 | intercept= -1.03 |
| Gpr37_Kcng2 | r = 0.17 | p-value= 0.06556 | slope = 0.04 | intercept= -0.94 |
| Gpr37_Kcng3 | r = -0.09 | p-value= 0.34359 | slope = -0.06 | intercept= -1.03 |
| Gpr37_Kcng4 | r = 0.04 | p-value= 0.62018 | slope = 0.00 | intercept= -0.99 |
| Gpr37_Kcnq1 | r = 0.04 | p-value= 0.67409 | slope = -0.01 | intercept= -0.99 |
| Gpr37_Kcnq2 | r = -0.09 | p-value= 0.30912 | slope = -0.17 | intercept= -0.89 |
| Gpr37_Kcnq3 | r = 0.11 | p-value= 0.23674 | slope = 0.40 | intercept= 0.10 |
| Gpr37_Kcnq4 | r = 0.13 | p-value= 0.15448 | slope = 0.07 | intercept= -0.92 |
| Gpr37_Kcnq5 | r = -0.07 | p-value= 0.43035 | slope = -0.52 | intercept= -1.00 |
| Gpr37_Kcnv1 | r = 0.01 | p-value= 0.94023 | slope = 0.03 | intercept= -0.96 |
| Gpr37_Kcnv2 | r = -0.10 | p-value= 0.25425 | slope = -0.03 | intercept= -1.02 |
| Gpr37_Kcns1 | r = 0.06 | p-value= 0.50481 | slope = -0.01 | intercept= -1.00 |
| Gpr37_Kcns2 | r = 0.06 | p-value= 0.52095 | slope = 0.02 | intercept= -0.97 |
| Gpr37_Kcns3 | r = 0.09 | p-value= 0.33318 | slope = -0.03 | intercept= -0.99 |
| Gpr37_Kcnh1 | r = -0.02 | p-value= 0.86502 | slope = -0.11 | intercept= -0.94 |
| Gpr37_Kcnh2 | r = 0.08 | p-value= 0.37978 | slope = -0.00 | intercept= -0.94 |
| Gpr37_Kcnh3 | r = 0.00 | p-value= 0.99270 | slope = -0.01 | intercept= -1.01 |
| Gpr37_Kcnh4 | r = 0.06 | p-value= 0.49217 | slope = -0.00 | intercept= -0.99 |
| Gpr37_Kcnh5 | r = -0.12 | p-value= 0.16876 | slope = -0.30 | intercept= -1.04 |
| Gpr37_Kcnh6 | r = 0.15 | p-value= 0.10324 | slope = 0.05 | intercept= -0.91 |
| Gpr37_Kcnh7 | r = -0.01 | p-value= 0.94520 | slope = 0.11 | intercept= -0.24 |
| Gpr37_Kcnh8 | r = 0.00 | p-value= 0.98281 | slope = -0.03 | intercept= -0.93 |
