## Supplementary material for "A bone-derived protein primes rapid visual escape via GPR37 receptor in a subpopulation of VTA GABAergic neurons": Table 2

| | | | | |
|---|---|---|---|---|
| **Gpr158_Kcnma1** | **r = 0.32** | **p-value= 0.00035** | **slope = 0.44** | **intercept= 0.08** |
| Gpr158_Kcnn1 | r = 0.10 | p-value= 0.26624 | slope = 0.01 | intercept= -0.98 |
| Gpr158_Kcnn2 | r = 0.14 | p-value= 0.13287 | slope = 0.41 | intercept= -0.30 |
| Gpr158_Kcnn3 | r = 0.13 | p-value= 0.14231 | slope = 0.14 | intercept= -0.77 |
| Gpr158_Kcnn4 | r = 0.13 | p-value= 0.13996 | slope = 0.00 | intercept= -1.00 |
| Gpr158_Kcnt1 | r = 0.08 | p-value= 0.39039 | slope = 0.11 | intercept= -0.77 |
| Gpr158_Kcnt2 | r = 0.08 | p-value= 0.38950 | slope = 0.25 | intercept= -0.57 |
| Gpr158_Kcnj2 | r = 0.14 | p-value= 0.11832 | slope = 0.00 | intercept= -0.98 |
| Gpr158_Kcnj12 | r = 0.18 | p-value= 0.04434 | slope = -0.10 | intercept= -1.00 |
| Gpr158_Kcnj4 | r = -0.07 | p-value= 0.46111 | slope = -0.02 | intercept= -0.99 |
| Gpr158_Kcnj14 | r = -0.02 | p-value= 0.79982 | slope = 0.00 | intercept= -0.99 |
| **Gpr158_Kcnj3** | **r = 0.29** | **p-value= 0.00104** | **slope = 0.51** | **intercept= -0.27** |
| **Gpr158_Kcnj6** | **r = 0.29** | **p-value= 0.00116** | **slope = 0.24** | **intercept= -0.62** |
| Gpr158_Kcnj9 | r = 0.16 | p-value= 0.07680 | slope = 0.09 | intercept= -0.90 |
| Gpr158_Kcnj5 | r = 0.14 | p-value= 0.11410 | slope = 0.02 | intercept= -0.99 |
| Gpr158_Kcnj10 | r = -0.19 | p-value= 0.03654 | slope = -0.09 | intercept= -0.97 |
| Gpr158_Kcnj16 | r = 0.05 | p-value= 0.56775 | slope = -0.01 | intercept= -0.97 |
| Gpr158_Kcnj8 | r = 0.04 | p-value= 0.63862 | slope = 0.00 | intercept= -1.00 |
| **Gpr158_Kcnj11** | **r = 0.24** | **p-value= 0.00718** | **slope = 0.02** | **intercept= -0.98** |
| Gpr158_Kcnj13 | r = 0.03 | p-value= 0.77333 | slope = 0.02 | intercept= -0.94 |
| Gpr158_Kcnk1 | r = 0.10 | p-value= 0.26299 | slope = 0.01 | intercept= -0.89 |
| Gpr158_Kcnk2 | r = 0.19 | p-value= 0.03648 | slope = 0.29 | intercept= -0.66 |
| Gpr158_Kcnk3 | r = 0.13 | p-value= 0.15160 | slope = 0.01 | intercept= -0.93 |
| Gpr158_Kcnk4 | r = 0.09 | p-value= 0.31404 | slope = 0.01 | intercept= -0.98 |
| Gpr158_Kcnk5 | r = -0.00 | p-value= 0.97165 | slope = 0.00 | intercept= -1.00 |
| Gpr158_Kcnk6 | r = -0.16 | p-value= 0.07458 | slope = 0.03 | intercept= -0.98 |
| Gpr158_Kcnk7 | r = 0.04 | p-value= 0.65590 | slope = 0.00 | intercept= -1.00 |
| Gpr158_Kcnk9 | r = 0.01 | p-value= 0.88676 | slope = -0.01 | intercept= -0.88 |
| **Gpr158_Kcnk10** | **r = 0.28** | **p-value= 0.00146** | **slope = 0.23** | **intercept= -0.71** |
| Gpr158_Kcnk12 | r = 0.04 | p-value= 0.62867 | slope = 0.02 | intercept= -0.96 |
| Gpr158_Kcnk13 | r = -0.20 | p-value= 0.02932 | slope = -0.13 | intercept= -0.93 |
| Gpr158_Kcnk15 | r = -0.14 | p-value= 0.13161 | slope = -0.01 | intercept= -1.00 |
| Gpr158_Kcnk18 | r = 0.15 | p-value= 0.10791 | slope = 0.00 | intercept= -1.00 |
| Gpr158_Kcna1 | r = -0.15 | p-value= 0.08963 | slope = -0.00 | intercept= -0.84 |
| Gpr158_Kcna2 | r = 0.08 | p-value= 0.40093 | slope = 0.06 | intercept= -0.85 |
| Gpr158_Kcna3 | r = 0.08 | p-value= 0.37020 | slope = 0.01 | intercept= -0.99 |
| Gpr158_Kcna4 | r = 0.13 | p-value= 0.16175 | slope = 0.03 | intercept= -0.93 |
| Gpr158_Kcna5 | r = 0.12 | p-value= 0.19192 | slope = 0.01 | intercept= -0.99 |
| Gpr158_Kcna6 | r = -0.17 | p-value= 0.05562 | slope = -0.20 | intercept= -0.91 |
| Gpr158_Kcna10 | r = 0.22 | p-value= 0.01463 | slope = 0.01 | intercept= -0.99 |
| Gpr158_Kcnb1 | r = 0.17 | p-value= 0.05694 | slope = 0.21 | intercept= -0.73 |
| Gpr158_Kcnb2 | r = 0.23 | p-value= 0.01034 | slope = 0.77 | intercept= 0.05 |
| Gpr158_Kcnc1 | r = 0.18 | p-value= 0.03973 | slope = -0.03 | intercept= -0.66 |
| Gpr158_Kcnc2 | r = 0.08 | p-value= 0.34917 | slope = 0.24 | intercept= -0.40 |
| Gpr158_Kcnc3 | r = 0.18 | p-value= 0.04065 | slope = 0.11 | intercept= -0.83 |
| Gpr158_Kcnc4 | r = 0.14 | p-value= 0.12761 | slope = -0.16 | intercept= -0.97 |
| Gpr158_Kcnd1 | r = -0.00 | p-value= 0.99242 | slope = -0.10 | intercept= -1.02 |
| Gpr158_Kcnd2 | r = 0.07 | p-value= 0.41218 | slope = 0.24 | intercept= 0.09 |
| Gpr158_Kcnd3 | r = 0.14 | p-value= 0.12508 | slope = 0.24 | intercept= -0.47 |
| Gpr158_Kcng1 | r = 0.15 | p-value= 0.10552 | slope = 0.01 | intercept= -0.99 |
| Gpr158_Kcng2 | r = -0.02 | p-value= 0.83414 | slope = 0.02 | intercept= -0.98 |
| **Gpr158_Kcng3** | **r = 0.29** | **p-value= 0.00130** | **slope = 0.07** | **intercept= -0.95** |
| Gpr158_Kcng4 | r = -0.06 | p-value= 0.52124 | slope = 0.00 | intercept= -0.99 |
| Gpr158_Kcnq1 | r = 0.03 | p-value= 0.75223 | slope = 0.01 | intercept= -0.98 |
| **Gpr158_Kcnq2** | **r = 0.35** | **p-value= 0.00007** | **slope = 0.34** | **intercept= -0.63** |
| Gpr158_Kcnq3 | r = 0.14 | p-value= 0.13065 | slope = 0.26 | intercept= -0.19 |
| Gpr158_Kcnq4 | r = 0.03 | p-value= 0.77334 | slope = -0.01 | intercept= -0.98 |
| Gpr158_Kcnq5 | r = 0.19 | p-value= 0.03414 | slope = 0.39 | intercept= -0.38 |
| Gpr158_Kcnv1 | r = -0.01 | p-value= 0.92806 | slope = -0.02 | intercept= -0.99 |
| **Gpr158_Kcnv2** | **r = 0.24** | **p-value= 0.00805** | **slope = 0.01** | **intercept= -0.99** |
| **Gpr158_Kcns1** | **r = 0.24** | **p-value= 0.00822** | **slope = 0.01** | **intercept= -0.99** |
| Gpr158_Kcns2 | r = 0.10 | p-value= 0.26694 | slope = 0.01 | intercept= -0.99 |
| Gpr158_Kcns3 | r = -0.02 | p-value= 0.85777 | slope = 0.03 | intercept= -0.96 |
| **Gpr158_Kcnh1** | **r = 0.27** | **p-value= 0.00270** | **slope = 0.22** | **intercept= -0.77** |
| Gpr158_Kcnh2 | r = 0.18 | p-value= 0.04297 | slope = 0.05 | intercept= -0.92 |
| Gpr158_Kcnh3 | r = 0.13 | p-value= 0.14350 | slope = 0.01 | intercept= -0.99 |
| Gpr158_Kcnh4 | r = 0.19 | p-value= 0.03450 | slope = 0.03 | intercept= -0.98 |
| **Gpr158_Kcnh5** | **r = 0.29** | **p-value= 0.00092** | **slope = 0.19** | **intercept= -0.69** |
| Gpr158_Kcnh6 | r = 0.11 | p-value= 0.24002 | slope = 0.04 | intercept= -0.95 |
| Gpr158_Kcnh7 | r = 0.15 | p-value= 0.10048 | slope = 0.07 | intercept= -0.33 |
| Gpr158_Kcnh8 | r = 0.14 | p-value= 0.11294 | slope = 0.11 | intercept= -0.87 |
